# Hyperglycemia-responsive circRNA drives diabetic neuropathic pain by competing for PUM2 binding and promoting proNT3/CCL2 in DRG neurons

**DOI:** 10.64898/2026.11.10.765753

**Authors:** Xu Li, Congcong Huang, Gokulapriya Govindarajalu, Bing Wang, Hong-Hong Zhang, Xiaozhou Feng, Dilip Sharma, Ruining Ma, Li Zhang, Bixin Zheng, Jifang Xiao, Marcella de A. Ferreira, Tolga Berkman, Jasmin Cabrera, Xiang Li, Baochun Jiang, Xue-Song Zhang, Steve Davidson, Yuan-Xiang Tao

**Affiliations:** Department of Anesthesiology, New Jersey Medical School, Rutgers, The State University of New Jersey, Newark, NJ07103, USA; Center for Advanced Biotechnology and Medicine, Rutgers University, Piscataway, NJ 08854, USA; Department of Molecular Pathobiology, NYU Pain Research Center, College of Dentistry, New York University, New York, NY10010, USA; Department of Physiology, Pharmacology & Neuroscience, New Jersey Medical School, Rutgers, The State University of New Jersey, Newark, NJ07103, USA; Department of Cell Biology & Molecular Medicine, New Jersey Medical School, Rutgers, The State University of New Jersey, Newark, NJ07103, USA

**Keywords:** *cRps6kb1*, neurotrophin-3 mRNA, proneurotrophin-3, PUM2, dorsal root ganglion, diabetic neuropathic pain

## Abstract

Diabetic neuropathic pain (DNP), one of the most common and debilitating complications of diabetes mellitus (DM), remains poorly managed by conventional analgesics, highlighting the urgent need to explore its etiology-associated mechanisms. Here, we identified a novel conserved circular RNA, cRps6kb1, that is significantly upregulated in dorsal root ganglion (DRG) neurons across multiple mouse models of type 1 DM and type 2 DM, as well as in human donors with DNP. Blocking cRps6kb1 upregulation attenuated the increases in DRG neurotrophin-3 mRNA (*Nt3*), its encoded proneurotrophin-3 (proNT3), and downstream C-C chemokine ligand 2 (CCL2) expression, as well as the persistence of nociceptive hypersensitivity under the conditions of streptozotocin-induced type 1 DM or high-fat diet-fed type 2 DM. Conversely, mimicking cRps6kb1 upregulation increased DRG *Nt3*, proNT3, and CCL2 levels and produced DNP-like symptoms in naive mice. In addition, high glucose markedly increased cRps6kb1 and *Nt3* expression in the cultured DRG neurons from mice and human donors. Mechanistically, upregulated cRps6kb1 competed with the *Nt3* 3’-untranslated region (3-UTR) for the translational repressor PUM2 binding, releasing PUM2 from the *Nt3* 3-UTR and enhancing *Nt3* mRNA translation and stability. Thus, cRps6kb1 is a key contributor to DNP and represents a promising therapeutic target for its treatment.

---

Neuropathic pain is a major unmet health problem caused by trauma, such as peripheral nerve or spinal cord injury, and diseases including diabetes, stroke, and HIV infection^1,2^. Although different forms may share common mechanisms, each has distinct molecular pathways shaped by its underlying etiology^1^. Diabetic peripheral neuropathy is among the most common and debilitating complications of diabetes and a leading cause of chronic neuropathic pain^3^. Diabetic neuropathic pain (DNP) is characterized by bilateral, symmetrical distal limb spontaneous pain, evoked allodynia and hyperalgesia, and/or sensory loss^3^. These symptoms substantially impair quality of life and are often accompanied by sleep disturbances, anxiety, depression, and functional disability, imposing significant socioeconomic burdens on patients and healthcare systems^3,4^. Current DNP treatments frequently provide inadequate pain relief and cause adverse effects because they primarily manage symptoms rather than target pathogenic mechanisms^4,5^. Identifying diabetes-specific molecular targets and elucidating mechanisms driving DNP may therefore enable more effective mechanism-based therapies.

Proneurotrophin-3 (proNT3) is the precursor of neurotrophin-3 (NT3), a member of the nerve growth factor family of neurotrophins^6,7^. Emerging evidence indicates that proNT3 is a biologically active signaling molecule that regulates neuronal excitability and neuroinflammatory responses^8,9^ and contributes to chronic neuropathic pain^10^. Peripheral nerve trauma or chemotherapy upregulates *Nt3* mRNA and proNT3 protein expression in neurons of the injured dorsal root ganglion (DRG), whereas mature NT3 protein remains undetectable under both normal and neuropathic pain conditions^11,12^. Genetic blockade of *Nt3* mRNA/proNT3 upregulation in the injured DRG attenuates nerve trauma- or paclitaxel-induced nociceptive hypersensitivity and increase of DRG C-C-chemokine ligand 2 (CCL2), a key mediator of neuropathic pain^11,12^. Conversely, DRG proNT3 overexpression induces neuropathic pain-like behaviors and elevates DRG CCL2 levels^11,12^. These effects are attenuated by DRG knockdown of tropomyosin receptor kinase C (TrkC)^11,12^, suggesting that proNT3 as a key instigator promotes neuropathic pain through TrkC-mediated CCL2 signaling in primary sensory neurons.

Circular RNAs (circRNAs) are endogenous non-coding RNAs characterized by covalently closed-loop structures lacking 5′ caps and 3′ poly(A) tails, making them highly stable and resistant to exonuclease degradation^13^. They are evolutionarily conserved across species, including mice and humans, and exhibit tissue-, cell-type-, and developmental-stage-specific expression^14,15^. circRNAs regulate gene expression and participate in diverse physiological and pathological processes through mechanisms including miRNA sponging, protein interactions, transcriptional regulation, and peptide translation^16,17^. Although several circRNAs, including *circAnks1a*, *circRNA-Filip1l*, and *circNrip1*, contribute to nerve trauma-induced neuropathic pain through distinct mechanisms^18–20^, circRNAs’ function in DNP remains largely unknown.

Here, we identified a novel, highly conserved circRNA, *cRps6kb1* (*cRps*), with 100% sequence identity between mice and humans. Hyperglycemia selectively upregulates *cRps* specifically in DRG neurons under diabetic conditions. *cRps* promotes DNP by competitively binding to the translational repressor PUM2, releasing it from the 3′-untranslated region (3′-UTR) of *Nt3* mRNA and enhancing *Nt3* mRNA translation and stability, thereby increasing proNT3 and downstream CCL2 expression in diabetic DRG neurons. Thus, *cRps* as an important contributor to DNP is a promising therapeutic target.

## RESULTS

### Identification of *cRps*

We identified *cRps* through deep analysis of our previous circRNA sequencing dataset^21^ and found significantly increased stacked reads of *cRps* in the DRG from the Akita mice, a well-established genetic model of type 1 diabetes mellitus (T1DM)^22^. Further analysis revealed that *cRps* originates from chromatin 11 (86,389,697–86,435,631) and spans the downstream exon 4 to the upstream exon 2 of the *Rps6kb1* gene, forming a 241-nt circular RNA (Fig. 1A). Sanger sequencing verified its back-splicing junction (Fig. 1B).

**Fig. 1.**
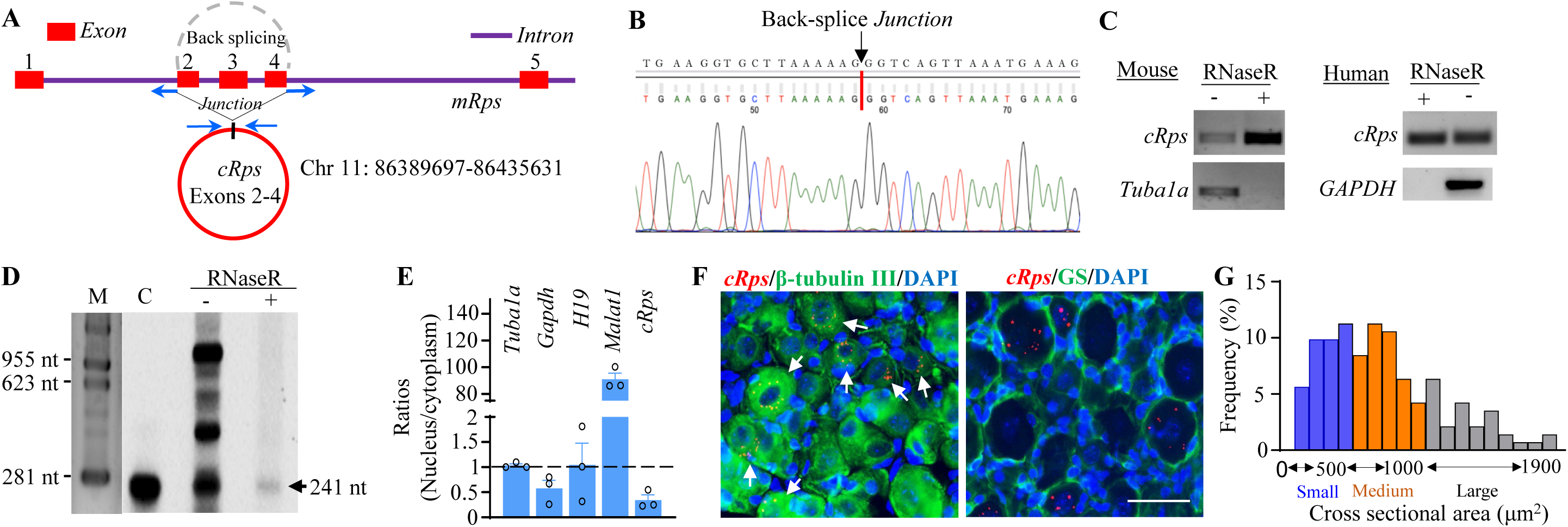
Identification and distribution of *cRps* in the DRG. (A) Schematic of the *cRps* genomic locus derived from linear *Rps6kb1* mRNA (*mRps*); arrow indicates the back-splice junction. (B) RT-PCR validation of the *cRps* back-splice junction using divergent primers and Sanger sequencing. (C) Strand-specific RT-PCR of *cRps* in naïve mouse and human lumbar DRGs with or without RNase R treatment; linear *Tuba1a* and *GAPDH* served as controls. M, DNA ladder. *n* = 3 replicates/species. (D) Northern blot validation of *cRps* in mouse DRGs with or without RNase R treatment. C, full-length *cRps*-expressing plasmid; M, RNA marker. *n* = 3 mice. (E) Subcellular distribution of *cRps*, *Tuba1a*, *Gapdh*, *H19*, and *Malat1* in cultured mouse DRG neurons. *n* = 3 mice. (F) Representative images showing cytoplasmic *cRps* (red) in β-tubulin III-positive neurons, but not GS-positive satellite cells, 4 weeks after STZ injection. Arrows indicate *cRps*-positive neurons (≥3 signal particles/cell). DAPI labels nuclei. Scale bar, 40 μm. (G) Distribution of *cRps*-labeled soma sizes in mouse L4 DRGs 4 weeks after STZ injection: small, 36.6%; medium, 40.9%; large, 22.5%. *n* = 5 mice.

To demonstrate its circular RNA characteristics, we assessed *cRps* stability and and resistance to RNase R digestion. *cRps* isolated from mice and human DRGs remained largely unchanged following RNase R treatment. In contrast, the linear *Tuba1a* and *GAPDH* mRNAs were markedly degraded (Fig. 1C). Following transcriptional arrest, *cRps* exhibited substantially slower degradation and a longer half-life than its linear counterpart, *Rps6kb1* mRNA (*mRps*; Supplementary Fig. 1A). Moreover, reverse transcription with oligo(dT) primers barely detected *cRps*, whereas *mRps* remained efficiently amplified, consistent with the absence of a poly(A) tail in *cRps* (Supplementary Fig. 1B). These findings confirmed the circular nature and stability of *cRps*.

Northern blotting detected *cRps* at the expected size in mouse DRG RNA following RNase R treatment (Fig. 1D). Under normal conditions, *cRps* was also detected in other tissues, including the spinal cord and trigeminal ganglion (Supplementary Fig. 1C). Subcellular fractionation demonstrated that *cRps* was preferentially enriched in the cytoplasm of DRG cells (Fig. 1E). BaseScope in situ hybridization combined with immunohistochemistry further characterized its cellular distribution. *cRps* signals were sparse or undetectable in naïve or vehicle-treated DRGs but were markedly increased 4 weeks after streptozotocin (STZ) injection, a mouse model of T1DM^22^ (Fig. 1F). Signals (≥ 3 particles per cell) were predominantly localized to neuronal cytoplasm rather than adjacent satellite cells (Fig. 1F). Approximately 22.3% of DRG neurons were *cRps*-positive 4 weeks following STZ injection, including 36.6% were small (< 25 µm in diameter), 40.9% were medium (25–35 µm), and 22.5% were large (> 35 µm in diameter) neurons (Fig. 1G). Among *cRps*-positive neurons, 33.4% expressed isolectin B4, 24.1% calcitonin gene-related peptide, 17.7% tyrosine hydroxylase, and 36.7% neurofilament-200 (Supplementary Fig. 1D). Together, these findings establish *cRps* as a stable, cytoplasmic circRNA enriched in diabetic DRG neurons, supporting its potential involvement in DNP.

### *cRps* upregulation in specific response to hyperglycemia in the context of diabetes

We next determined whether STZ injection induced time-dependent *cRps* upregulation in the DRG. *cRps*, but not *mRps*, was significantly increased bilaterally at 2, 3, 4, and 6 weeks after STZ injection (Fig. 2A-B). Consistently, the numbers of *cRps*-labelled neurons, number of labelled particles in cytoplasm and total labelled particles per positive neuron were also elevated in the DRGs 4 weeks post-STZ injection (Fig. 2C-E). In contrast, *cRps* expression remained unchanged in other tissues examined following STZ injection (Supplementary Fig. 2A–C). *cRps* was similarly upregulated in DRGs (not in the trigeminal ganglion) of the Akita mice (Fig. 2F; Supplementary Fig. 2D) and non-obese diabetic (NOD) mice (a spontaneous mouse model of T1DM^22^; Fig. 2G) as well as high-fat diet-fed mice (a mouse model of type 2 diabetes mellitus (T2DM)^22^; Fig. 2H) and db/db mice (a genetic mouse model of T2DM^22^; Fig. 2I). Importantly, *cRps*, but not *mRps*, was markedly increased in DRGs from human donors with DNP, accompanied by increased *cRps*-positive neurons and cytoplasmic particles (Fig. 2J–L). *cRps* upregulation positively correlated with nociceptive hypersensitivity (Supplementary Fig. 2E–L) and elevated blood glucose levels in diabetic mice (Supplementary Fig. 3A–E) and human donors. High glucose also selectively increased *cRps* in cultured mouse and human DRG neurons, whereas two other pain-associated circRNAs, *circNRIP1*^18^ and *circPlekha*^19^, were unaffected (Fig. 2M–N; Supplementary Fig. 3F). In contrast, *cRps* expression was unchanged in models of nerve trauma-, chemotherapy-, inflammatory-, or surgical pain (Supplementary Fig. 3G–L). Together, these findings demonstrate that *cRps* upregulation is specifically associated with hyperglycemia in DRG neurons under DNP conditions.

**Fig. 2.**
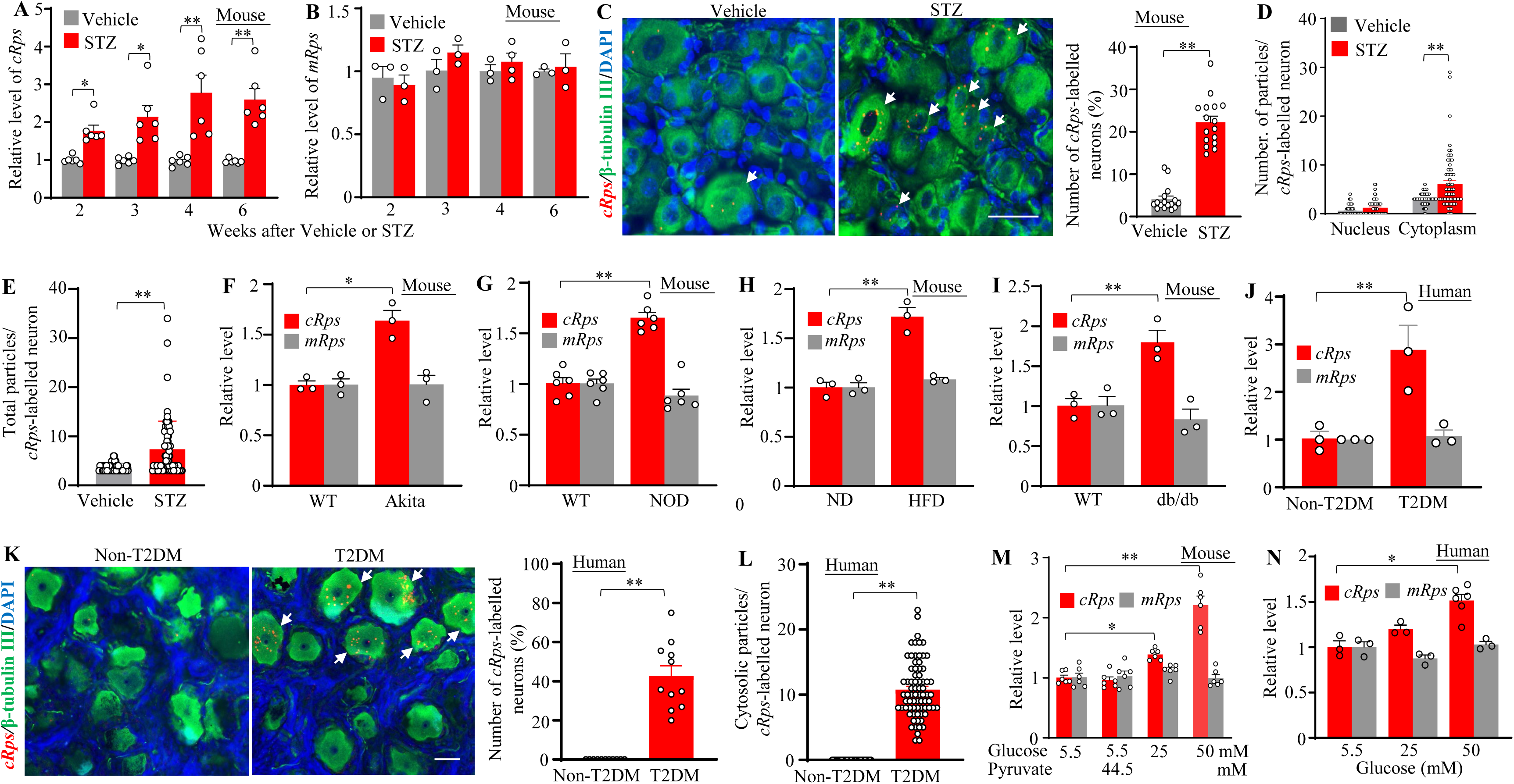
Upregulation of *cRps* expression in DRG neurons of mice and humans under diabetic conditions. (A, B) *cRps* and linear *Rps6kb1* mRNA (*mRps*) expression in lumbar DRGs of male mice 2, 3, 4, and 6 weeks after STZ or vehicle injection. *n* = 3–6 mice/time point/group. \**P* < 0.05, \*\**P* < 0.01, two-way ANOVA with Tukey’s post hoc test. (C–E) Representative images and quantification of *cRps*-positive lumbar DRG neurons (C), nuclear and cytoplasmic *cRps* signal particles (D), and total particles per neuron (E) 4 weeks after STZ or vehicle injection. *n* = 5 mice/group. \*\**P* < 0.01, paired Student’s *t*-test. Scale bar, 40 μm. (F–I) *cRps* and *mRps* expression in lumbar DRGs from Akita (F), NOD (G), HFD-fed (15 weeks; H), and db/db (I) mice and their respective controls. *n* = 3–6 mice/group. \**P* < 0.05, \*\**P* < 0.01, unpaired Student’s *t*-test. (J) *cRps* and *mRps* expression in lumbar DRGs from donors with painful type 2 diabetes (T2DM) and non-diabetic controls. *n* = 3 donors/group. \*\**P* < 0.01, unpaired Student’s *t*-test. (K, L) Representative images and quantification of *cRps*-positive human DRG neurons (K) and cytoplasmic *cRps* signal particles per neuron (L) in T2DM and non-diabetic donors. *n* = 3 donors/group. \*\**P* < 0.01; paired *t*-test (K), unpaired *t*-test (L). Scale bar, 40 μm. (M, N) *cRps* and *mRps* expression in cultured mouse (M) and human (N) DRG neurons exposed to different glucose concentrations. *n* = 3–6 replicates/concentration. \**P* < 0.05, \*\**P* < 0.01, one-way ANOVA with Tukey’s post hoc test.

### Blocking DRG *cRps* upregulation alleviates DNP

To determine whether DRG *cRps* upregulation contributes to DNP, we blocked STZ-induced DRG *cRps* upregulation by microinjecting AAV5-cRps-shRNA into unilateral L3/L4 DRGs of T1DM male mice, with AAV5-Scr-shRNA as control. Because patients with DNP typically seek treatment only after the onset of DNP, we assessed the effect of AAV5-cRps-shRNA on established DNP by microinjecting the virus into the DRGs 3 days after STZ or vehicle administration. This experimental design was based on the 3–4 weeks required for AAV5-mediated transgene expression in the DRG^23–25^. AAV5-cRps-shRNA selectively suppressed *cRps* upregulation in the ipsilateral (not contralateral) DRGs 35 days post-STZ and attenuated established mechanical, heat, and cold hypersensitivity at 21–35 days, but not 14 days, after microinjection (Fig. 3A–E; Supplementary Fig. 4A–D). It also reduced stimulation-independent spontaneous pain (evidenced by a robust preference for the lidocaine-paired chamber) and ipsilateral dorsal horn neuronal/glial hyperactivity (documented by the increases in the levels of p-ERK1/2 and GFAP), respectively, 35 days after microinjection (Supplementary Fig. 4E–G). Similar effects were observed in female mice that received the same AAV5 microinjection 3 days after STZ administration (Supplementary Fig. 5A–K), indicating that DRG *cRps* upregulation is required for maintaining STZ-induced DNP regardless of sex.

**Fig. 3.**
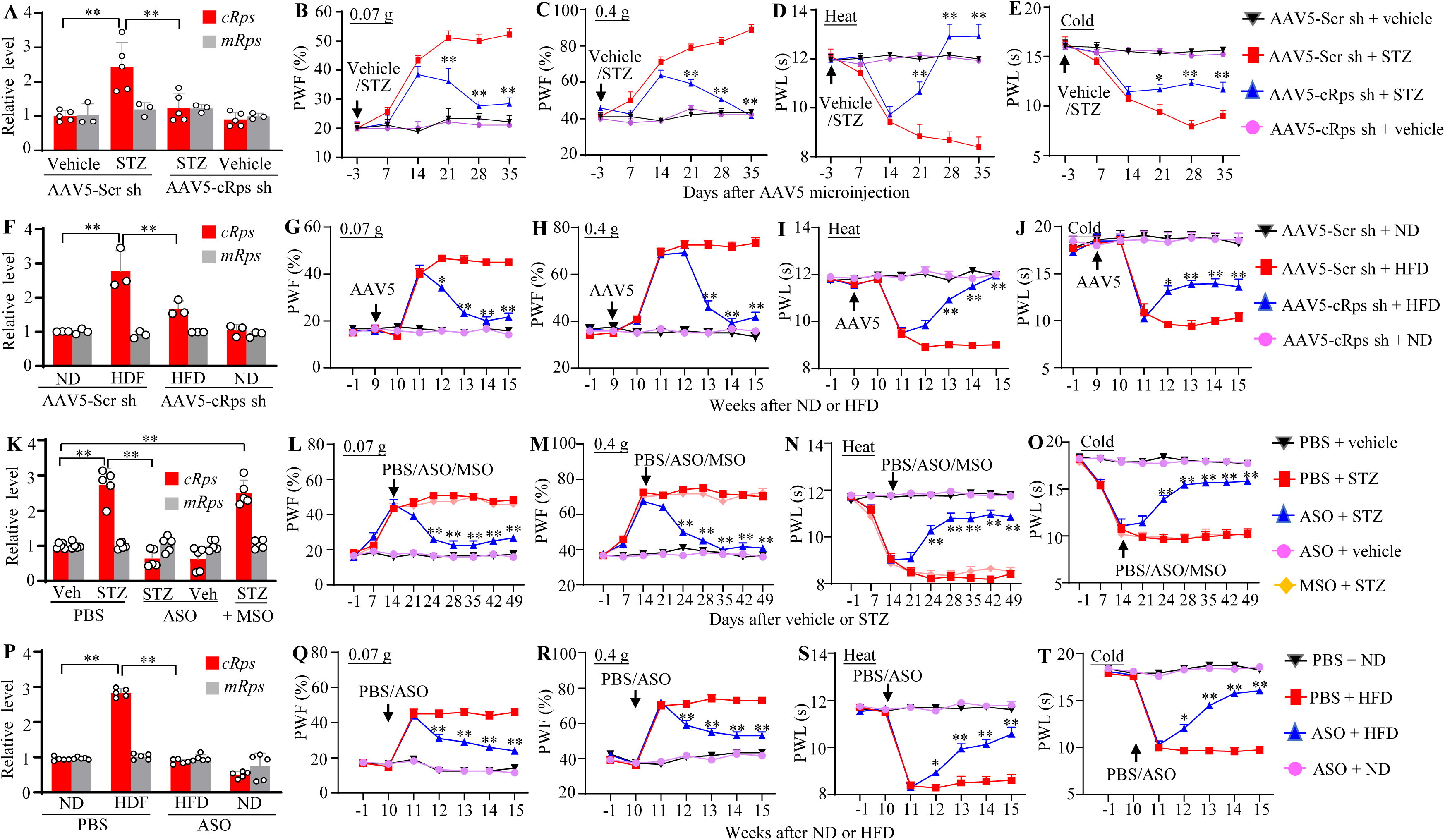
Blocking *cRps* upregulation in the DRG alleviates established diabetic neuropathic pain in male mice with T1DM or T2DM. (A) Expression of *cRps* and *mRps* in ipsilateral L3/L4 DRGs 35 days after unilateral AAV5-cRps sh or scrambled control (AAV5-Scr sh) injection in STZ- or vehicle-treated mice. *n* = 3–5 replicates (2 mice/replicate)/group. \*\**P* < 0.01, two-way ANOVA with Tukey’s post hoc test. (B–E) Effects of AAV5-cRps sh or AAV5-Scr sh injection 3 days after STZ or vehicle treatment on ipsilateral paw withdrawal frequency (PWF) to 0.07-g and 0.4-g von Frey stimuli and paw withdrawal latency (PWL) to heat and cold stimuli. *n* = 9 mice/group. \**P* < 0.05, \*\**P* < 0.01 versus STZ + AAV5-Scr sh at corresponding time points; three-way repeated-measures ANOVA with Tukey’s post hoc test. (F) *cRps* and *mRps* expression in ipsilateral L3/L4 DRGs after 15 weeks of normal (ND) or high-fat diet (HFD); AAV5-cRps sh or AAV5-Scr sh was injected 9 weeks after diet initiation. *n* = 3 replicates (2 mice/replicate)/group. \*\**P* < 0.01, two-way ANOVA with Tukey’s post hoc test. (G–J) Effects of AAV5-cRps sh or AAV5-Scr sh injection after 9 weeks of ND or HFD on ipsilateral PWF to 0.07-g and 0.4-g von Frey stimuli and PWL to heat and cold stimuli. *n* = 12 mice/group. \**P* < 0.05, \*\**P* < 0.01 versus HFD + AAV5-Scr sh at corresponding time points; three-way repeated-measures ANOVA with Tukey’s post hoc test. (K) *cRps* and *mRps* expression in bilateral L3/L4 DRGs 49 days after STZ or vehicle treatment and 35 days after intrathecal (i.t.) PBS, *cRps*-specific antisense oligonucleotide (ASO; 100 μg), or missense control (MSO; 100 μg) administration. *n* = 5 replicates (5 mice/replicate)/group. \*\**P* < 0.01, two-way ANOVA with Tukey’s post hoc test. (L–O) Effects of i.t. PBS, ASO, or MSO administered 2 weeks after STZ or vehicle treatment on left-side PWF to 0.07-g and 0.4-g von Frey stimuli and PWL to heat and cold stimuli. *n* = 12 mice/group. \*\**P* < 0.01 versus STZ + PBS at corresponding time points; three-way repeated-measures ANOVA with Tukey’s post hoc test. (P) *cRps* and *mRps* expression in bilateral L3/L4 DRGs after 15 weeks of ND or HFD; ASO (100 μg) or PBS was administered i.t. at week 10. *n* = 5 mice/group. \*\**P* < 0.01, two-way ANOVA with Tukey’s post hoc test. (Q–T) Effects of i.t. PBS or ASO (100 μg) administered after 10 weeks of ND or HFD on left-side PWF to 0.07-g and 0.4-g von Frey stimuli and PWL to heat and cold stimuli. *n* = 12 mice/group. \*\**P* < 0.01 versus HFD + PBS at corresponding time points; three-way repeated-measures ANOVA with Tukey’s post hoc test.

We next examined a diet-induced obesity model of T2DM^22^. AAV5-cRps-shRNA or control virus was microinjected into unilateral L3/L4 DRGs 9 weeks after initiation of high-fat diet (HFD) or normal diet (ND) feeding. One week later (week 10), HFD-fed mice received a low-dose intraperitoneal (i.p.) injection of STZ, whereas ND-fed mice received a vehicle injection via the same route. Nociceptive hypersensitizes developed by week 11 and persisted for at least through week 15. Similar to the findings in the STZ-induced T1DM model, AAV5-cRps-shRNA suppressed DRG *cRps* upregulation 15 weeks post-HFD feeding and attenuated established mechanical allodynia and heat/cold hyperalgesia at weeks 12–15, but not week 11, after HFD feeding (Fig. 3F–J; Supplementary Fig. 6A–D). It also reduced spontaneous pain and ipsilateral dorsal horn neuronal/glial hyperactivity (Supplementary Fig. 6E–G). Thus, DRG *cRps* upregulation contributes to DNP maintenance in both T1DM and T2DM models.

To independently validate these findings, we developed a *cRps*-specific antisense oligonucleotide (ASO). Intrathecal (i.t.) administration of *cRps* ASO, but not PBS or missense oligonucleotide, after DNP onset (that is, 14 days after STZ injection) suppressed *cRps* upregulation in the bilateral L3/4 DRGs 49 days post-STZ injection (Fig. 3K). This microinjection also markedly mitigated mechanical, heat and cold nociceptive hypersensitivities 24, 28, 35, 42 and 49 days, but not 14 and 21 days, post-STZ on both sides (Fig. 3L-O; Supplementary Fig. 7A-C). *cRps* ASO abolished spontaneous pain as well as neuronal/glial hyperactivity in the bilateral spinal dorsal horns at 49 days post-STZ injection (Supplementary Fig. 7D–F). Similar analgesic effects were observed following i.t. *cRps* ASO treatment at 10-week post-initiation of HDF feeding in T2DM mice (Fig. 3P–T; Supplementary Fig. 8A–F). Collectively, these results further establish a role for DRG *cRps* in maintaining DNP.

Importantly, neither AAV5-cRps-shRNA nor *cRps* ASO altered basal mechanical, heat, or cold responses, locomotor function, fasting glucose, HbA1c, or intraepidermal nerve fiber density in control or diabetic mice (Fig. 3B–E, 3G–J, 3L-O and 3Q-T; Supplementary Fig. 4B– D, 5B-H, 6B-D, 7A-C, 8A-C and 9A–N; Supplementary Table 1). In HFD-fed mice, body weight, glucose tolerance, and insulin resistance were also unaffected (Supplementary Fig. 10A– F). Thus, inhibition of DRG *cRps* selectively alleviates established DNP without disrupting acute nociception, normal somatosensation, locomotion, or the metabolic and peripheral neuropathological features of diabetes.

### DRG *cRps* overexpression produces diabetic neuropathic pain-like symptoms

Is DRG *cRps* upregulation sufficient for DNP? To answer this question, we overexpressed full-length *cRps* in unilateral L3/L4 DRGs of naïve male mice by AAV5-cRps microinjection, using AAV5-GFP as control. AAV5-cRps selectively increased *cRps*, but not linear *mRps*, expression 8 weeks after microinjection (Fig. 4A). *cRps* overexpression produced ipsilateral mechanical, heat, and cold nociceptive hypersensitivity beginning 4 weeks after microinjection and persisting through 8 weeks (Fig. 4B–E), accompanied by spontaneous pain and neuronal/glial hyperactivity in the ipsilateral spinal dorsal horn 8 weeks post-microinjection (Fig. 4F–H). Similar effects were observed in female mice (Fig. 4I–P). Neither AAV5 microinjection altered contralateral basal nociceptive responses or locomotor function (Fig. 4B-D and 4J-L; Supplementary Table 1). These findings clearly demonstrate that DRG *cRps* upregulation leads to DNP-like symptoms in both male and female mice in the absence of diabetes.

**Fig. 4.**
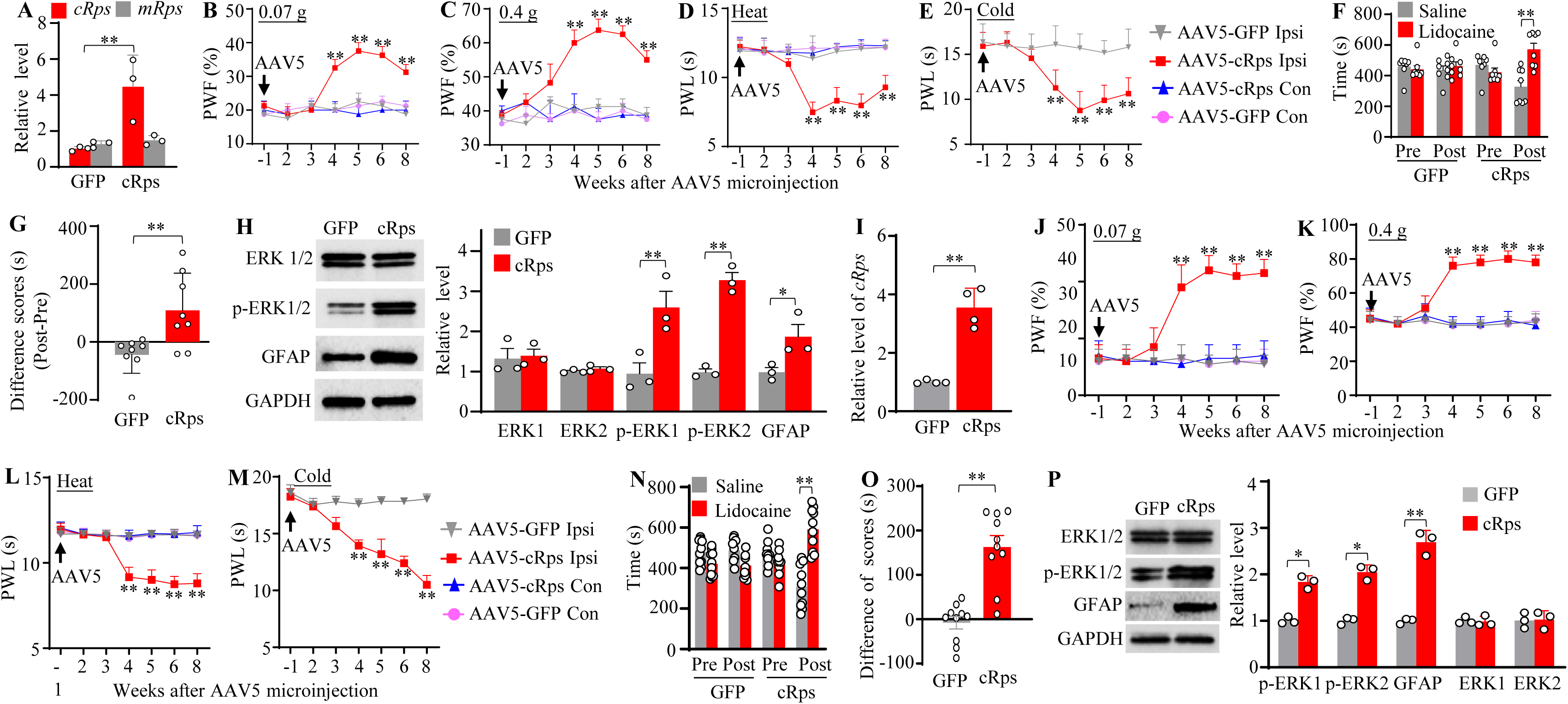
DRG overexpression of *cRps* produces diabetic neuropathic pain–like symptoms. (A) Expression of *cRps* and *mRps* in ipsilateral L3/L4 DRGs 8 weeks after unilateral AAV5-cRps or AAV5-GFP injection in male mice. *n* = 3 replicates (2 mice/replicate)/group. \*\**P* < 0.01, two-tailed unpaired Student’s *t*-test. (B–E) Paw withdrawal frequency (PWF) to 0.07-g and 0.4-g von Frey stimuli and paw withdrawal latency (PWL) to heat and cold stimuli at indicated weeks post-injection in male mice, assessed ipsilaterally (Ipsi) and contralaterally (Con). *n* = 8 mice/group. \*\**P* < 0.01 versus AAV5-GFP at corresponding time points; two-way repeated-measures ANOVA with Tukey’s post hoc test. (F, G) Conditioned place preference (CPP) assessment of spontaneous pain 7 weeks post-injection in male mice. Pre, preconditioning; Post, post-conditioning. *n* = 8 mice/group. \*\**P* < 0.01; two-way repeated-measures ANOVA with Tukey’s post hoc test (F) or two-tailed unpaired Student’s *t*-test (G). (H) p-ERK1/2, ERK1/2, and GFAP levels in ipsilateral L3/L4 dorsal horns 8 weeks post-injection in male mice. *n* = 3 mice/group. \*\**P* < 0.01, two-tailed unpaired Student’s *t*-test. (I) *cRps* and *mRps* expression in ipsilateral L3/L4 DRGs 8 weeks after AAV5-cRps or AAV5-GFP injection in female mice. *n* = 4 replicates (2 mice/replicate)/group. \*\**P* < 0.01, two-tailed unpaired Student’s *t*-test. (J–M) PWF to 0.07-g and 0.4-g von Frey stimuli and PWL to heat and cold stimuli at the indicated weeks post-injection in female mice, assessed ipsilaterally and contralaterally. *n* = 10 mice/group. \*\**P* < 0.01 versus AAV5-GFP at corresponding time points; two-way repeated-measures ANOVA with Tukey’s post hoc test. (N, O) CPP assessment of spontaneous pain 7 weeks post-injection in female mice. Pre, preconditioning; Post, post-conditioning. *n* = 10 mice/group. \*\**P* < 0.01; two-way repeated-measures ANOVA with Tukey’s multiple-comparisons test (N) or two-tailed unpaired Student’s *t*-test (O). (P) p-ERK1/2, ERK1/2, and GFAP levels in ipsilateral L3/L4 dorsal horns 8 weeks post-injection in female mice. *n* = 3 mice/group. \*\**P* < 0.01, two-tailed unpaired Student’s *t*-test.

### *cRps*-controlled *Nt3* mRNA and its encoded proNT3 protein in diabetic DRG neurons as key drivers of DNP

To identify downstream targets of DRG *cRps* upregulation under DNP conditions, we performed high-throughput RNA sequencing of the microinjected L3/4 DRGs 35 days after vehicle or STZ administration in mice receiving AAV5-cRps-shRNA or AAV5-Scr-shRNA microinjection. Among 21,554 detected genes, 1,887 were differentially expressed in STZ-treated mice, including 231 upregulated and 1,656 downregulated genes (Supplementary Fig. 11A). Of these, blocking *cRps* upregulation by DRG AAV5-cRps-shRNA microinjection reversed the expression of 143 upregulated and 813 downregulated genes (Supplementary Fig. 11B), which were enriched in biological processes related to development, anatomical structure, and morphogenesis (Supplementary Fig. 11C).

Among the most strongly *cRps*-regulated genes, *Nt3* is essential for neuronal development, survival, and function^6^. Consistent with RNA sequencing results (Supplementary Fig. 11B), *Nt3* mRNA and proNT3 protein increased progressively in bilateral DRGs at 2, 3, 4, and 6 weeks following STZ injection and were accompanied by increased numbers of *Nt3* mRNA-positive neurons 4 weeks post-STZ injection (Fig. 5A-C). Similar increases occurred in the bilateral DRGs 15 weeks after initiation of HFD feeding (Fig. 5D-E) as well as in the DRGs of NOD mice, db/db mice and human donors with DNP (Fig. 5F-I). High glucose also increased *Nt3* mRNA in cultured mouse and human DRG neurons (Fig. 5J–K). As reported previously^11,12^, mature NT3 remained undetectable in the DRGs post-STZ or post-HFD/ND feeding in mice and from human donors with or without DM (Supplementary Fig. 12A-C).

**Fig. 5.**
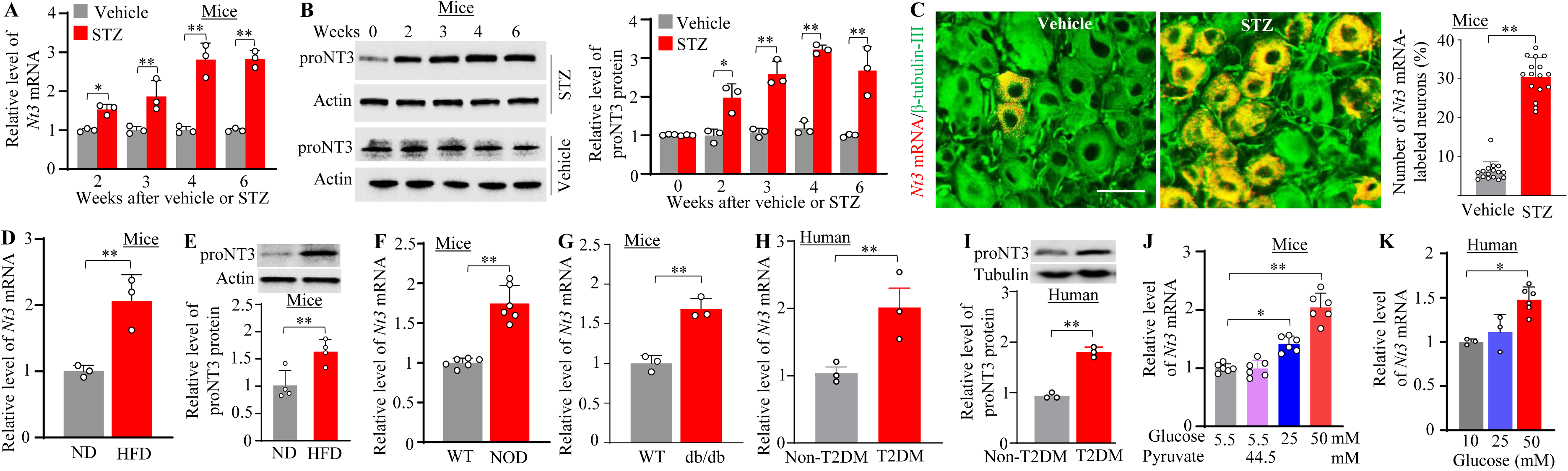
Increased *Nt3* mRNA and proNT3 protein expression in DRG of mice and humans under diabetic conditions. (A, B) *Nt3* mRNA and proNT3 protein expression in lumbar DRGs of male mice 0–6 weeks after STZ or vehicle injection. *n* = 3 mice/time point/group. \**P* < 0.05, \*\**P* < 0.01, two-way ANOVA with Tukey’s post hoc test. (C) Representative images and quantification of *Nt3* mRNA-positive lumbar DRG neurons 4 weeks after injection. *n* = 5 mice/group. \*\**P* < 0.01, paired Student’s *t*-test. Scale bar, 40 μm. (D, E) *Nt3* mRNA and proNT3 protein levels in lumbar DRGs after 15 weeks of ND or high-fat diet (HFD). *n* = 4 mice/group. \*\**P* < 0.01, unpaired Student’s *t*-test. (F, G) *Nt3* mRNA expression in lumbar DRGs of NOD (F) and db/db (G) male mice and respective WT controls. *n* = 3–6 mice/group. \*\**P* < 0.01, unpaired Student’s *t*-test. (H, I) *Nt3* mRNA and proNT3 protein levels in lumbar DRGs from donors with painful type 2 diabetes (T2DM) and non-diabetic controls. *n* = 3 donors/group. \*\**P* < 0.01, unpaired Student’s *t*-test. (J, K) *Nt3* mRNA expression in cultured mouse (J) and human (K) DRG neurons exposed to different glucose concentrations. *n* = 3–6 replicates/concentration. \**P* < 0.05, \*\**P* < 0.01, one-way ANOVA with Tukey’s post hoc test..

To examine whether DRG proNT3 increase is required for established DNP, we blocked *Nt3* mRNA increase in the unilateral L3/L4 DRGs using *Nt3*-specific siRNA microinjection 21 days after STZ injection (Fig. 6A). This treatment significantly attenuated mechanical allodynia and heat/cold hyperalgesia on the ipsilateral, but not contralateral, side starting day 24 through day 28 post-STZ and alleviated neuronal/glial hyperactivity in the ipsilateral L3/4 spinal dorsal horns 28 days post-STZ (Fig. 6B-E; Supplementary Fig. 13A-E). Consistent results were obtained using specific *Nt3* gene knockdown in the unilateral L3/L4 DRGs of male and female *Nt3*^f/f^ mice following AAV5-Cre microinjection 21 days before STZ or vehicle injection (Supplementary Fig. 14A-K and 15A-K). In the STZ-treated both male and female mice, this knockdown also mitigated the spontaneous pain (Supplementary Figs. 14I-J and 15I-J). Neither intervention altered basal nociceptive responses in the vehicle-treated mice or locomotor function (Fig. 6B-E and Supplementary Fig. 13A-C, Fig. 14B-H and 15B-H; Supplementary Tabel 1).

**Fig. 6.**
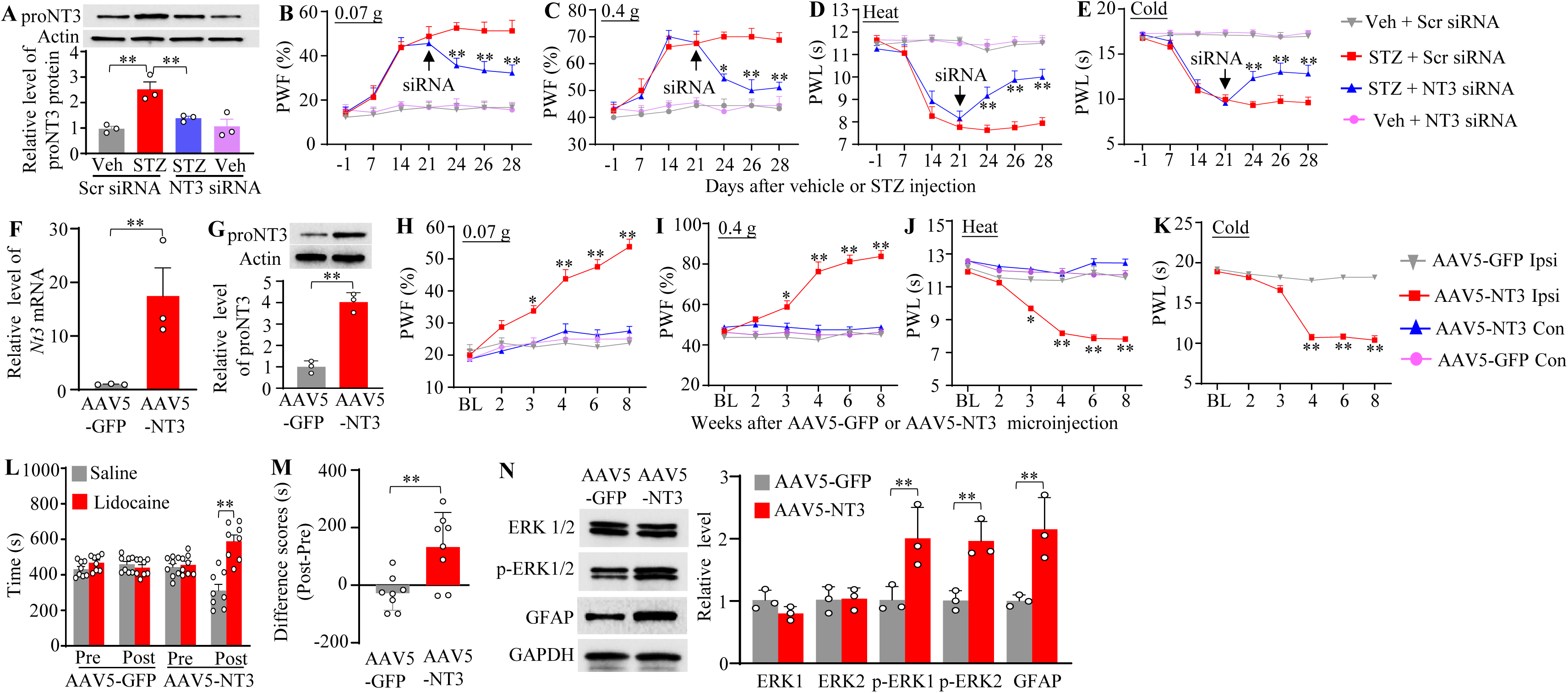
Role of DRG proNT3 in the STZ-induced diabetic neuropathic pain. (A) ProNT3 protein levels in ipsilateral L3/L4 DRGs 28 days after STZ or vehicle injection and 7 days after NT3-specific or scrambled control siRNA injection. *n* = 3 replicates (2 mice/replicate)/group. \*\**P* < 0.01, one-way ANOVA with Tukey’s post hoc test. (B–E) Effects of NT3 or scrambled siRNA injected 21 days after STZ or vehicle treatment on ipsilateral paw withdrawal frequency (PWF) to 0.07-g and 0.4-g von Frey stimuli and paw withdrawal latency (PWL) to heat and cold stimuli. *n* = 8 mice/group. \**P* < 0.05, \*\**P* < 0.01 versus STZ + Scr siRNA at corresponding time points; three-way repeated-measures ANOVA with Tukey’s post hoc test. (F, G) *Nt3* mRNA and proNT3 protein levels in ipsilateral L3/L4 DRGs 8 weeks after AAV5-NT3 or AAV5-GFP injection. *n* = 3 replicates (2 mice/replicate)/group. \*\**P* < 0.01, unpaired Student’s *t*-test. (H–K) Effects of AAV5-NT3 or AAV5-GFP on ipsilateral and contralateral PWF to 0.07-g and 0.4-g von Frey stimuli and PWL to heat and cold stimuli at indicated weeks post-injection. *n* = 8 mice/group. \**P* < 0.05, \*\**P* < 0.01 versus AAV5-GFP at corresponding time points; two-way repeated-measures ANOVA with Tukey’s post hoc test. (L, M) Conditioned place preference (CPP) assessment of spontaneous pain 7 weeks after AAV5-NT3 or AAV5-GFP injection. Pre, preconditioning; Post, post-conditioning. *n* = 8 mice/group. \*\**P* < 0.01; unpaired Student’s *t*-test (L) or two-way repeated-measures ANOVA with Tukey’s post hoc test (M). (N) p-ERK1/2, ERK1/2, and GFAP levels in ipsilateral L3/L4 dorsal horns 8 weeks after AAV5-NT3 or AAV5-GFP injection. *n* = 3 mice/group. \*\**P* < 0.01, unpaired Student’s *t*-test.

We next tested whether DRG proNT3 increase was sufficient to induce DNP. Full-length *Nt3* was overexpressed in the unilateral L3/L4 DRGs of naïve male mice using DRG microinjection of AAV5-NT3 (Fig. 6F-G). Compared with AAV5-GFP, AAV5-NT3 produced persistent ipsilateral mechanical, heat, and cold hypersensitivity beginning 3–4 weeks after microinjection and lasting at least 8 weeks (Fig. 6H–K). proNT3 overexpression also induced spontaneous pain and neuronal/astrocytic hyperactivity in the ipsilateral L3/4 spinal dorsal horn 8 weeks after AAV5-NT3 microinjection (Fig. 6L-N). Similar effects were observed in female mice (Supplementary Fig. 16A-I). Neither viral microinjection altered locomotor function (Supplementary Table 1).

Together, these findings establish DRG proNT3 as a critical player in DNP, consistent with its previously demonstrated role in nerve trauma- and chemotherapy-induced neuropathic pain^11,12^.

### *cRps* upregulation is responsible for proNT3 increase in diabetic DRG neurons

Consistent with our RNA-sequencing findings (Supplementary Fig. 11B), DRG-specific knockdown of *cRps* with AAV5-cRps shRNA, but not AAV5-Scr shRNA, attenuated the STZ-induced increases in *Nt3* mRNA and proNT3 protein, as well as their downstream *Ccl2* mRNA and CCL2 protein, in the microinjected L3/L4 DRGs 35 days after microinjection (Fig. 7A-B and Supplementary Fig. 17A-B). Similar inhibitory effects were observed 15 weeks after HFD feeding (Fig. 7C-D and Supplementary Fig. 17C-D). Consistently, i.t. *cRps* ASO, but not PBS, abolished the increases of *Nt3* mRNA, proNT3 protein, *Ccl2* mRNA and CCL2 protein in bilateral L3/L4 DRGs 49 days after STZ injection or 15 weeks HFD feeding (Fig. 7E-H and Supplementary Fig. 17E-H). In cultured DRG neurons, *cRps* knockdown by AAV5-cRps shRNA transduction markedly suppressed high glucose (50 mM)-induced increases in *Nt3* mRNA and proNT3 protein levels (Fig. 7I-J). Together, these findings suggest that *cRps* upregulation contributes to *Nt3*/proNT3 induction and downstream *Ccl2* mRNA and CCL2 expression in diabetic DRG neurons and links hyperglycemia to this pathway in both T1DM and T2DM.

**Fig. 7.**
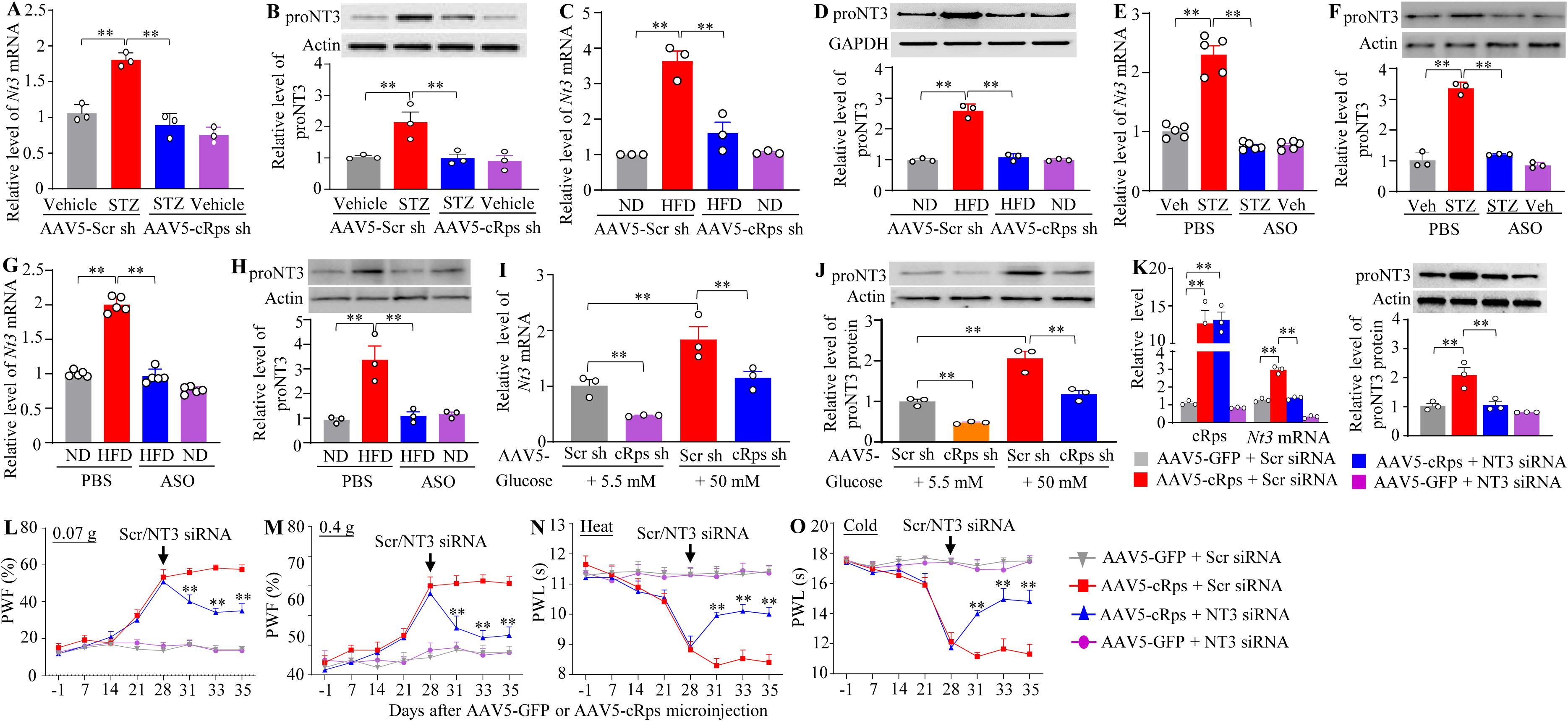
*cRps* upregulation contributes to increased *Nt3* mRNA and proNT3 Protein expression in the DRG under diabetic neuropathic pain conditions. (A-D) *Nt3* mRNA (A, C) and proNT3 protein (B, D) levels in ipsilateral L3/L4 DRGs following AAV5-cRps sh or AAV5-Scr sh injection in STZ- or vehicle-treated mice (A, B; DRGs collected 35 days post-injection) or ND- and HFD-fed mice (C, D; injection at week 9, collection at week 15). *n* = 3–5 replicates (2 mice/replicate)/group. \*\**P* < 0.01, two-way ANOVA with Tukey’s post hoc test. (E–H) *Nt3* mRNA (E, G) and proNT3 protein (F, H) levels in bilateral L3/L4 DRGs after intrathecal (i.t.) ASO, MSO, or PBS administration in STZ- or vehicle-treated mice (E, F; 2 weeks after treatment, collection on day 49) or ND- and HFD-fed mice (G, H; administration at week 10, collection at week 15). *n* = 3–5 replicates (2 mice/replicate)/group. \*\**P* < 0.01, two-way ANOVA with Tukey’s post hoc test. (I, J) *Nt3* mRNA (I) and proNT3 protein (J) levels in cultured mouse DRG neurons exposed to 5.5 or 50 mM glucose and transduced with AAV5-cRps sh or AAV5-Scr sh for 3 days. *n* = 3 replicates/group. \*\**P* < 0.01, one-way ANOVA with Tukey’s post hoc test. (K) *Nt3* mRNA and proNT3 protein levels in ipsilateral L3/L4 DRGs 35 days after AAV5-cRps or AAV5-GFP injection, followed by NT3 or scrambled (Scr) siRNA co-injection on day 28. *n* = 3 replicates (2 mice/replicate)/group. \*\**P* < 0.01, one-way ANOVA with Tukey’s post hoc test. (L–O) Effects of NT3 or Scr siRNA co-injection on day 28 after AAV5-cRps or AAV5-GFP injection on ipsilateral paw withdrawal frequency (PWF) to 0.07-g (L) and 0.4-g (M) von Frey stimuli and paw withdrawal latency (PWL) to heat (N) and cold (O) stimuli. *n* = 9 mice/group. \*\**P* < 0.01 versus AAV5-cRps + Scr siRNA at corresponding time points; two-way repeated-measures ANOVA with Tukey’s post hoc test.

To further establish proNT3 as a functional downstream mediator of *cRps*, we examined whether DRG-specific *Nt3* knockdown could alleviate *cRps* overexpression-induced nociceptive hypersensitivity. Specific NT3 siRNA or control scrambled siRNA was microinjected into the ipsilateral L3/L4 DRGs 28 days after unilateral microinjection of AAV5-cRps or AAV5-GFP in naïve male mice. AAV5-cRps, but not AAV5-GFP, increased *cRps*, *Nt3* mRNA, proNT3 protein, *Ccl2* mRNA, and CCL2 protein in the microinjected DRGs 35 days after AAV5 microinjection (Fig. 7K and Supplementary Fig. 18A), accompanied by ipsilateral mechanical allodynia, heat- and cold-induced hyperalgesia from days 28 to 35 after AAV5 microinjection, and increased neuronal/glial hyperactivity in the L3/L4 dorsal horn on day 35 post-AAV5 microinjection (Fig. 7L-O and Supplementary Fig. 18B-E). Importantly, NT3 siRNA markedly attenuated these molecular and behavioral changes, indicating that proNT3 is required for *cRps* overexpression-induced DRG CCL2 increase and nociceptive hypersensitivity. These effects appeared to be *cRps*-specific, as AAV5-cRps-induced increases in *Nt3* mRNA and proNT3 protein were abolished by co-transduction of AAV5-cRps shRNA in DRG neurons (Supplementary Fig. 18F).

Collectively, our findings indicate that proNT3 and its downstream effector CCL2 are key mediators of the contribution of *cRps* upregulation to DNP in DRG neurons.

### *cRps* competitively binds to PUM2 to promote the expression of *Nt3* mRNA and proNT3 in diabetic DRG

Finally, we investigated how *cRps* upregulation increased *Nt3* mRNA and proNT3 levels in diabetic DRGs. RNA-binding proteins (RBPs) play a critical role in RNA biogenesis by regulating various aspects of RNA expression and metabolism^18,26,27^. To search for *cRps*-interacting RBPs, we conducted the comprehensive identification of RNA-binding proteins by mass spectrometry (ChIRMS) assay coupled with the analysis of PANTHER Classification System database. Approximately 11 candidate RBPs were identified to interact with *cRps* in cultured DRG neurons (cRps/mRps ratio ≥1.5; protein coverage >10%; Supplementary Table 2). Among these, pumilio homolog 2 (PUM2) was chosen as a potential *cRps* partner because a PUM2-binding element (PBE; UGUANAUA) was identified in the *Nt3* 3′-UTR using the RBPSuite web server.

Given that PUM2 represses target gene expression by binding to mRNA 3′-UTRs^28,29^, we hypothesized that *cRps* upregulation competitively sequesters PUM2, thereby relieving PUM2-mediated destabilization of *Nt3* mRNA. Consistent with this hypothesis, *cRps*, but not *mRps*, interacted with PUM2 in cultured DRG neurons (Fig. 8A). RIP assays further showed that PUM2 bound both *cRps* and the *Nt3* 3′-UTR in the ipsilateral L3/4 DRGs of AAV5-Scr shRNA– microinjected mice 4 weeks after vehicle injection (Fig. 8B). STZ treatment increased *cRps*-PUM2 association by 2.7-fold, while reducing PUM2 binding to the *Nt3* 3′-UTR by 71%, in the ipsilateral L3/4 DRGs of AAV5-Scr shRNA–microinjected mice 4 weeks after STZ injection (Fig. 8B). However, these changes were absent following the blockade of *cRps* upregulation in the ipsilateral L3/4 DRGs of AAV5-cRps shRNA–microinjected mice 4 weeks after STZ injection (Fig. 8B). Notably, STZ injection did not alter the basal expression levels of *Pum2* mRNA and PUM2 protein in the ipsilateral L3/4 DRGs from 2 to 6 weeks post-injection (Supplementary Fig. 19A-B). These findings suggest that *cRps* upregulation promotes PUM2 sequestration and release from *Nt3* mRNA, resulting in stabilization of *Nt3* mRNA in diabetic DRG neurons.

**Fig. 8.**
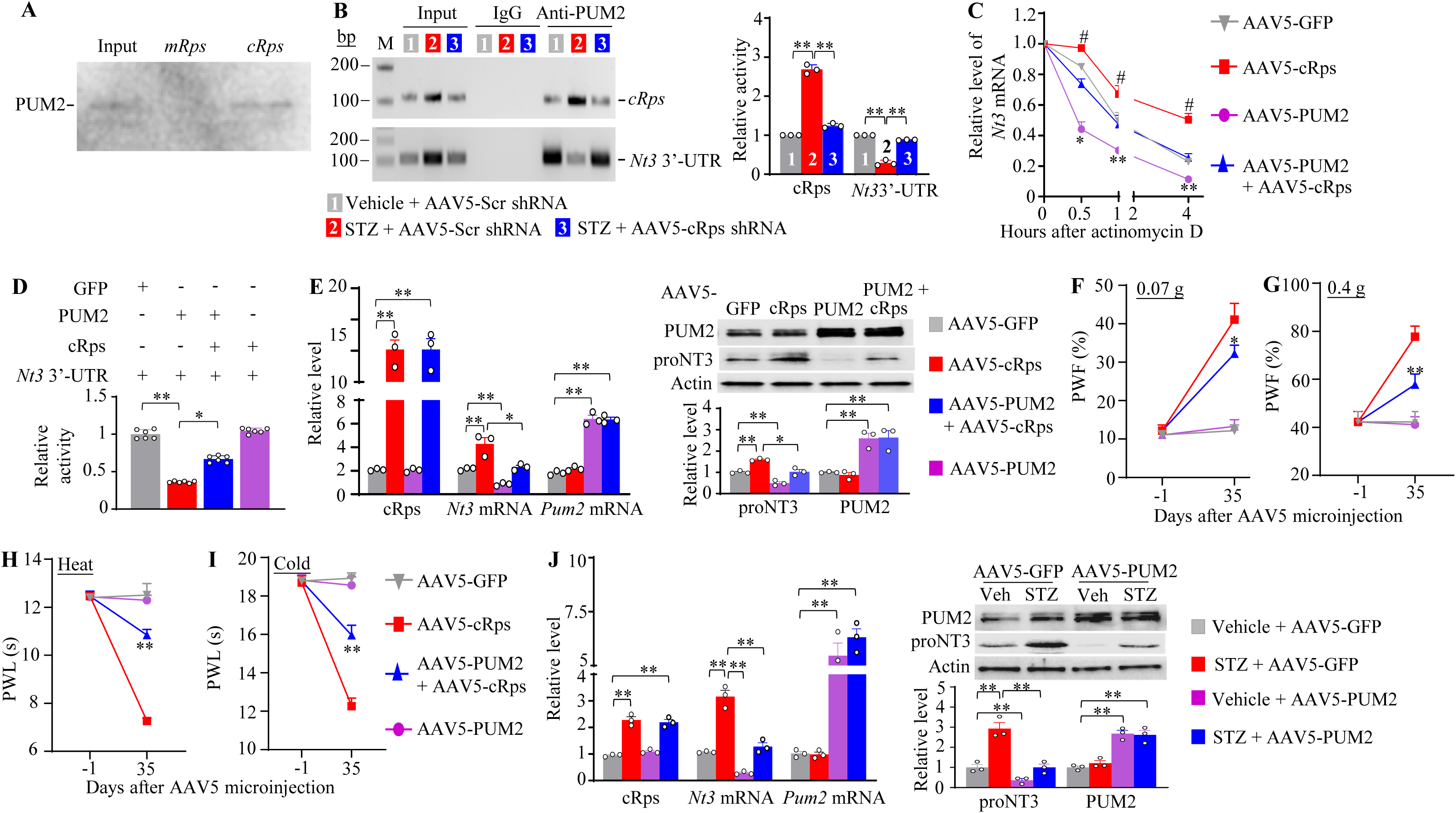
cRps competes with *Nt3* 3’-UTR for PUM2 binding to promote *Nt3* mRNA and proNT3 protein expression in diabetic DRGs. (A) PUM2 pulldown by labeled exogenous *cRps*, but not *mRps*, in cultured DRG neurons. *n* = 3 replicates. (B) *cRps* and *Nt3* 3′-UTR enrichment in PUM2 immunoprecipitates from ipsilateral L3/L4 DRGs 4 weeks after AAV5-cRps shRNA or scrambled (Scr) shRNA injection in STZ- or vehicle-treated male mice. *n* = 3 replicates (45 mice)/group. \*\**P* < 0.01, two-way ANOVA with Tukey’s post hoc test. (C) *Nt3* mRNA stability in AAV5-transduced CAD cells treated with actinomycin D (5 µg/ml) for 0–4 h, measured by qRT-PCR. *n* = 3 replicates/group/time point. \**P* < 0.05, \*\**P* < 0.01, #*P* < 0.05 versus AAV5-GFP at corresponding time points; two-way ANOVA with Tukey’s post hoc test. (D) *Nt3* 3′-UTR activity in HEK293T cells transfected with vectors expressing *Gfp*, *cRps*, *Pum2*, or the *Nt3* 3′-UTR. *n* = 6 replicates/group. \**P* < 0.05, \*\**P* < 0.01, one-way ANOVA with Tukey’s post hoc test. (E) *cRps*, *Nt3* mRNA, *Pum2* mRNA, PUM2 protein, and proNT3 protein levels in ipsilateral L3/L4 DRGs 35 days after injection of the indicated AAV5 vectors. *n* = 3 replicates (6 mice)/group/assay. \**P* < 0.05, \*\**P* < 0.01, one-way ANOVA with Tukey’s post hoc test. (F–I) Effects of AAV5-PUM2, AAV5-cRps, AAV5-GFP, or combined AAV5-PUM2/AAV5-cRps injection on ipsilateral paw withdrawal frequency (PWF) to 0.07-g (F) and 0.4-g (G) von Frey stimuli and paw withdrawal latency (PWL) to heat (H) and cold (I). *n* = 10 mice/group. \**P* < 0.05, \*\**P* < 0.01 versus AAV5-cRps at corresponding time points; two-way repeated-measures ANOVA with Tukey’s post hoc test. (J) *cRps*, *Nt3* mRNA, *Pum2* mRNA, PUM2 protein, and proNT3 protein levels in ipsilateral L3/L4 DRGs 35 days after the indicated AAV5 injections in STZ- or vehicle-treated male mice. *n* = 3 replicates (6 mice)/group/assay. \*\**P* < 0.01, one-way ANOVA with Tukey’s post hoc test.

Our *in vitro* RNA stability assay further showed that PUM2 overexpression by transducing AAV5-PUM2 significantly increased the decay rate of *Nt3* mRNA at 0.5, 1, and 4 hours after transcriptional arrest with actinomycin D (ActD) treatment in cultured CAD cells (Fig. 8C). In contrast, *cRps* overexpression by co-transducing AAV5-cRps prevented this effect (Fig. 8C). AAV5-cRps transduction alone also markedly slowed basal decay of *Nt3* mRNA at 0.5, 1, and 4 hours after ActD treatment in CAD cells (Fig. 8C). Moreover, we conducted a luciferase reporter assay, in which the *Nt3* 3′-UTR was cloned downstream of a luciferase reporter gene, to further verify the blocking role of *cRps* in PUM2-triggered instability of the *Nt3* 3′-UTR in CAD cells. Transfection of the vector expressing PUM2 produced a significant decrease in the luciferase reporter activity (Fig. 8D), suggesting that PUM2 overexpression destabilizes the *Nt3* mRNA. This decrease was noticeably reversed in the presence of co-transfection of the vector expressing *cRps* (Fig. 8D). Similarly, AAV5-PUM2 transduction in cultured DRG neurons substantially reduced *Nt3* mRNA and proNT3 protein levels (Supplementary Fig. 19C). This reduction was completely rescued by co-transduction of AAV5-cRps (Supplementary Fig. 19C). Together, these *in vitro* and *in vivo* findings strongly support a model in which *cRps* as a competitor binds to PUM2, releasing it from the *Nt3* 3′-UTR and thereby stabilizing *Nt3* mRNA and increasing proNT3 expression in diabetic DRG neurons.

Our *in vivo* experiments further demonstrated that DRG PUM2 overexpression in the ipsilateral L3/4 DRGs, achieved by AAV5-PUM2 microinjection, significantly attenuated the AAV5-cRps-induced increases in *Nt3* mRNA and proNT3 protein levels in the microinjected DRGs 5 weeks after viral microinjection (Fig. 8E). Importantly, this overexpression also mitigated AAV5-cRps-induced mechanical allodynia and heat/cold hyperalgesia on the ipsilateral, but not contralateral, side (Fig. 8F-I and Supplementary Fig. 19D-F), as well as neuronal/glial hyperactivity in the ipsilateral L3/4 dorsal horn (Supplementary Fig. 19G) at 5 weeks after viral microinjection. Similarly, microinjection of AAV5-PUM2 into the unilateral L3/4 DRGs 3 days after STZ or vehicle administration, completely attenuated the STZ-induced increases in *Nt3* mRNA and proNT3 protein levels in the microinjected DRGs (Fig. 8J), as well as mechanical allodynia and heat/cold hyperalgesia on the ipsilateral, but not contralateral, side (Supplementary Fig. 20A-G) and neuronal/glial hyperactivity in the ipsilateral L3/4 dorsal horn (Supplementary Fig. 20H), 5 weeks after viral microinjection. Given the co-expression of *cRps*, *Pum2*, and *Nt3* mRNAs in DRG neurons of different sizes (Supplementary Fig. 21), these findings further indicate that PUM2 functions as a negative regulator in *cRps*-triggered expression of *Nt3* mRNA and proNT3 protein in diabetic DRG neurons and in *cRps* contribution to DNP.

## DISCUSSION

DNP has been studied for decades, yet its molecular mechanisms remain incompletely understood. Here, we identified *cRps* as a unique hyperglycemia-responsive circRNA selectively upregulated in DRG neurons from diabetic mice and human donors with DNP. *cRps* contributes to the maintenance of both T1DM- and T2DM-induced DNP by competing with the *Nt3* 3′-UTR for PUM2 binding, thereby releasing PUM2 from *Nt3* 3′-UTR, stabilizing *Nt3* mRNA, and increasing proNT3 and downstream CCL2 expression. These findings identify *cRps* as a key contributor to DNP.

Unlike circRNAs previously implicated in neuropathic pain, including *circNrip1*^18^, *circRNA-Filip1l*^19^ and *circAnks1a*^20^, *cRps* exhibits remarkable specificity for DNP. Although expressed at low basal levels in mouse and human DRGs, *cRps* was consistently upregulated across multiple T1DM and T2DM models, as well as in human donors with DNP. In contrast, its expression was unchanged in models of peripheral nerve trauma, chemotherapy, chronic inflammation, or surgical pain. *cRps* upregulation was also restricted to the DRG and was selectively induced by high glucose in cultured mouse and human DRG neurons, suggesting a direct association with diabetic hyperglycemia. The mechanisms underlying this selective response remain unknown. Because RBPs regulate circRNA biogenesis through back-splicing^29,30^, future studies should determine whether specific RBPs promote *cRps* formation under hyperglycemic conditions. In addition, examining *cRps* expression in diabetic patients with and without DNP^31,32^ will clarify whether its upregulation is specifically associated with DNP, non-painful diabetic peripheral neuropathy, diabetes itself, or their combination.

Mechanistically, *cRps* promotes *Nt3* mRNA stability and proNT3 expression by competing with the *Nt3* 3′-UTR for PUM2 binding. PUM2 is an evolutionarily conserved RNA-binding protein^33^ that generally represses translation and promotes mRNA decay by binding Pumilio-binding elements (PBEs) within target 3′-UTRs^28,34^. We found that the *Nt3* 3′-UTR contains functional PBEs through which PUM2 promotes *Nt3* mRNA instability. PUM2 overexpression enhanced *Nt3* mRNA degradation, whereas *cRps* overexpression attenuated this effect. Consistent with this mechanism, diabetic conditions increased the association of *cRps* with PUM2 while reducing PUM2 binding to the *Nt3* 3′-UTR, without altering total PUM2 expression. Because *cRps*, *Pum2* mRNA, and *Nt3* mRNA are co-expressed in individual DRG neurons, these findings support a model in which hyperglycemia-induced *cRps* sequesters PUM2 and promotes its release from *Nt3* 3’-UTR, thereby stabilizing *Nt3* mRNA and enhancing its translation. It is worth noting that transcription factors and epigenetic modifications may also contribute to the increase in DRG *Nt3* mRNA under DNP conditions. Future studies are needed to determine whether these mechanisms interact with *cRps*/PUM2 and contribute to the increase in *Nt3* mRNA in diabetic DRG neurons.

Our findings further establish a *cRps*–PUM2–Nt3/proNT3–CCL2 pathway in DRG neurons under DNP conditions. proNT3 is regarded as a key player in nerve trauma- or chemotherapy-induced neuropathic pain, by promoting TrkC-mediated expression of CCL2, a key endogenous contributor of neuropathic pain^35^, in DRG neurons^10^. In this study, *Nt3* mRNA and proNT3 protein were increased in DRG neurons across multiple diabetic mouse models and in human donors with DNP, as well as in cultured DRG neurons exposed to high glucose. DRG-specific *Nt3* knockdown attenuated STZ-induced nociceptive hypersensitivity, whereas *Nt3* overexpression in naïve mice produced mechanical, thermal, and cold hypersensitivity and spontaneous pain. Consistent with previous studies^11,12^, mature NT3 protein was undetectable in the DRG under normal and diabetic conditions, supporting a predominant role for proNT3 rather than mature NT3 in DNP. Importantly, genetic or ASO-mediated inhibition of *cRps* reduced *Nt3* mRNA, proNT3, and CCL2 expression and alleviated established nociceptive hypersensitivity in both T1DM and T2DM models. Conversely, *cRps* overexpression increased these molecules and produced pain hypersensitivity, effects that were reversed by DRG *Nt3* knockdown. DRG-specific PUM2 overexpression similarly suppressed *cRps*- or diabetes-induced increases in *Nt3* and proNT3 and reduced nociceptive hypersensitivity. Together, these findings indicate the *cRps*-triggered the PUM2–Nt3/proNT3–CCL2 pathway in diabetic DRG neurons contributes, at least in part, to DNP persistence. It should be acknowledged that *cRps* also regulated many other genes based on our RNA sequencing analysis. Thus, the involvement of *cRps*-triggered proNT3-indepndent mechanisms in DNP could be ruled out.

In summary, we identified a novel hyperglycemia-responsive *cRps*-triggered mechanism by which *Nt3* mRNA stability was increased and proNT3 and CCL2 levels were elevated in diabetic DRG neurons. Importantly, a single i.t. administration of *cRps*-specific ASO produced sustained analgesia for more than 25 days in mice with established T1DM- or T2DM-associated nociceptive hypersensitivity without altering acute pain, normal somatosensation, or locomotor function. Given its complete sequence conservation between mice and humans and the clinical precedent for i.t. ASO administration^36,37^, *cRps* represents a promising therapeutic target for DNP.

## MATERIALS AND METHODS

### Animals

Adult male and female CD1 and C57BL/6J mice were obtained from Charles River Laboratories, and *Nt3*^f/f^, Akita, and db/db mice from Jackson Laboratory. NOD/ShiLtJ mice were bred and housed at Rutgers under an approved IACUC protocol (eIACUC #201900017). Mice were maintained under standard conditions with a 12-h light/dark cycle and ad libitum food and water. All procedures were IACUC-approved and complied with NIH and International Association for the Study of Pain guidelines. Animals were randomly assigned, investigators were blinded, and efforts were made to minimize suffering and optimize sample sizes.

### Human DRG tissues

Human lumbar DRG tissues from de-identified organ donors were obtained from AnaBios Corporation (San Diego, CA), along with anonymized medical histories, including diabetes status and pain symptoms. Samples from non-diabetic donors served as controls, whereas those from donors with diabetes and pain symptoms comprised the experimental group.

### STZ-induced T1DM mouse model and glucose measurement

CD1 or C57BL/6J mice (7–8 weeks old) were fasted overnight before receiving a single intraperitoneal injection of streptozotocin (STZ; 165 mg/kg; Sigma) in sodium citrate buffer (pH 4.5), as previously described ^38,39^. Control mice received an equivalent volume of vehicle (0.1 mL/10 g body weight). To prevent acute hypoglycemia, mice received 10% glucose solution instead of drinking water for 24 h after injection. Blood glucose was measured using a OneTouch Ultra glucose meter (LifeScan) 3 days later. Mice with STZ-induced blood glucose levels >270 mg/dL were considered diabetic and included in subsequent experiments.

### High-fat diet (HFD)-induced T2DM mouse model

The HFD-induced T2DM model was established as previously described ^40,41^, with minor modifications. C57BL/6J mice were fed a normal diet (ND) or high-fat diet (HFD; D12492; Research Diets, New Brunswick, NJ) for 10 weeks, followed by overnight fasting and a single i.p. injection of STZ (75 mg/kg) or vehicle (sodium citrate buffer, pH 4.5; 0.1 mL/10 g body weight). Blood glucose and body weight were monitored every 1–2 weeks. HFD-fed mice with blood glucose levels >270 mg/dL were considered diabetic and included in subsequent experiments.

### Glucose or insulin tolerance tests

Glucose and insulin tolerance tests were performed 1–2 weeks after STZ injection in ND- or HFD-fed mice following overnight fasting with free access to water. Baseline blood glucose was measured from the tail vein using a OneTouch Ultra glucose meter, followed by i.p. injection of glucose (1 g/kg) for the glucose tolerance test or insulin (0.25 U/kg) for the insulin tolerance test. Blood glucose was measured at 30, 60, 90, and 120 min post-injection. Glucose tolerance was assessed by plotting blood glucose levels over time and calculating the area under the curve (AUC).

### Blood HbA1c measurements

Blood HbA1c levels were measured using a mouse-compatible assay kit according to the manufacturer’s instructions. Anticoagulated whole blood was processed, and HbA1c levels were determined using the provided calibration method and expressed as a percentage of total hemoglobin.

### Other mouse pain models

Neuropathic pain was induced by L4 spinal nerve ligation (SNL) or sciatic nerve chronic constriction injury (CCI) under isoflurane anesthesia, as previously described ^23–25^. For SNL, the L4 spinal nerve was ligated with 7–0 silk and transected distally; for CCI, the sciatic nerve was loosely ligated at four sites (1-mm spacing). Sham surgeries omitted nerve ligation or transection. Chemotherapy-induced neuropathy was induced by i.p. paclitaxel (PTX; 4 mg/kg) on days 0, 2, 4, and 6, using 50% Cremophor EL and 50% ethanol as the vehicle ^23–25^. Inflammatory pain was induced by plantar injection of complete Freund’s adjuvant (CFA; 20 μL) or intra-articular injection of mono-iodoacetate (MIA; 5 μL, 50 μg/μL in 0.9% saline) ^23–25^. Incisional pain was induced by a 0.8-cm longitudinal incision through the skin, fascia, and muscle of the plantar hind paw ^23–25^.

### Behavioral tests

Mice were habituated to the testing environment for 1–2 h daily for 2–3 days before baseline testing. Mechanical, heat, and cold sensitivity were assessed sequentially at 1-h intervals. Conditional place preference (CPP) was tested at least 5 weeks after viral microinjection, and locomotor function was assessed before tissue collection.

Mechanical sensitivity: Mice were acclimated for 30 min in Plexiglas chambers on a wire-mesh platform. Calibrated von Frey filaments (0.07 and 0.4 g) were applied to the plantar surfaces of both hind paws 10 times at 5-minute intervals. Paw withdrawal frequency (PWF) was calculated as the percentage of positive responses.

Heat sensitivity: Paw withdrawal latency (PWL) was measured using a radiant heat source applied to the plantar surface of each hind paw. Three to four trials were performed at 5-min intervals, with a 20-s cutoff to prevent tissue damage.

Cold sensitivity: PWL was measured on a temperature-monitored cold plate maintained at 0°C. Each hind paw was tested three times at 10-min intervals, with a 20-s cutoff.

Spontaneous pain: CPP was assessed using a two-chamber apparatus with distinct visual and tactile cues (Med Associates). Following 15-min preconditioning, mice underwent 3 days of conditioning, pairing intrathecal (i.t.) saline (5 μL) with one chamber and lidocaine (0.4%, 5 μL) with the opposite chamber, 6 h apart. The injection order was reversed daily. On test day, mice were allowed to explore both chambers freely for 15 min. CPP was quantified as the difference between test and preconditioning times spent in the lidocaine-paired chamber.

Locomotor function was evaluated using placing, grasping, and righting reflex tests. Each reflex was assessed five times at 5-min intervals, and performance was quantified as the number of normal responses.

### DRG microinjection

DRG microinjection was performed as previously described, with minor modifications ^23–25^. Under 2% isoflurane anesthesia, the L3/L4 articular processes were removed to expose the DRGs. siRNA (40–80 μM) or viral solution (4–9 × 10¹² vg/mL; 1 μL/DRG) was injected at 0.1 μL/min using a glass micropipette, which remained in place for 10 min to minimize backflow. The surgical site was irrigated with sterile saline and sutured closed.

### Intrathecal injection

Under 2% isoflurane anesthesia, mice underwent intrathecal injection at the L4–L5 level. Oligonucleotide solution or PBS (1 μL) was slowly injected via a glass micropipette, which remained in place for 5 min to minimize reflux. The surgical site was rinsed with sterile saline and closed with wound clips.

### Plasmid constructs and virus production

Full-length *Nt3*, *Pum2*, and *cRps* cDNAs were amplified from mouse DRG RNA using the Platinum Taq High Fidelity PCR Kit (Invitrogen/Thermo Fisher Scientific) and cloned into pAAV-CMV-circ (*cRps*) or pHpa-trs-SK (*Nt3* and *Pum2*) vectors following restriction digestion. For *cRps* knockdown, shRNA targeting the unique back-splice junction was inserted into pAAV-shRNA-EF1α-EYFP. AAV5 particles carrying the respective constructs were packaged and purified using the AAVpro Purification Kit (Takara), and viral titers were determined using the AAVpro Titration Kit according to the manufacturer’s instructions. Primer and shRNA sequences are provided in Supplementary Table 3.

### CAD cell or mouse DRG neuronal culture and transfection/transduction

CAD cells and primary mouse DRG neurons were prepared as previously described ^23–25^. CAD cells were cultured in DMEM/F-12 supplemented with 8% FBS and 1% antibiotics. DRGs from mice aged ≥4 weeks were enzymatically dissociated using dispase and collagenase, mechanically triturated, and plated on poly-D-lysine-coated six-well plates (1.5–4 × 10⁵ cells/well) in supplemented Neurobasal medium. On day 2, cells were transduced with AAV (4– 10 μL; ≥6–9 × 10¹² vg/mL), transfected with plasmid DNA (300–500 ng), or treated with siRNA (100 nM) using Lipofectamine 3000. Cells were harvested 3 days later for subsequent analyses.

#### Human DRG neuronal culture

Human lumbar DRGs were obtained from consented, de-identified cadaveric donors through Network for Hope (Cincinnati, OH). The University of Cincinnati IRB determined that tissue procurement and use did not constitute human subjects research (Study ID: 2015-5302). Primary neuronal cultures were prepared as previously described ^42^, with minor modifications. Briefly, DRGs were cleaned in ice-cold NMDG-aCSF, minced, and enzymatically digested with papain (45 U, 35 min, 37°C) followed by collagenase (4.5 mg, 35 min, 37°C). Tissues were triturated, filtered through a 100-μm strainer, centrifuged, and resuspended in Neurobasal A supplemented with B27 Plus, penicillin/streptomycin, 2 mM GlutaMAX, and 5% FBS. Cells were plated on poly-D-lysine (0.1 mg/mL)/collagen (0.2 mg/mL)-coated plates or coverslips and maintained at 37°C with 5% CO₂. The medium was replaced every other day, and neurons recovered for at least 24 h before experiments.

### Quantitative real-time reverse transcription (RT)-PCR assay

Total RNA from mouse/human tissues and cultured cells was extracted using the RNeasy Mini Kit (Qiagen) and treated with DNase I (New England Biolabs) to remove genomic DNA. Human DRG RNA was also obtained from Clontech Laboratories. RNA purity was assessed spectrophotometrically (A260/A280, 1.97–2.08). cDNA was synthesized using the Omniscript Reverse Transcription Kit (Qiagen) with gene-specific or oligo(dT) primers. qRT-PCR was performed using SYBR Green Master Mix on a Bio-Rad CFX96 system. Relative expression was calculated using the 2⁻ΔΔCt method, normalized to day 0 for time-course studies or the corresponding control for treatment comparisons. No-template controls showed no amplification. Primer sequences are provided in Supplementary Table 3.

### Single-cell RT-PCR assay

Freshly dissociated adult mouse DRG neurons were classified by diameter 4 h after plating as large (>35 μm), medium (25–35 μm), or small (<25 μm). Individual neurons were collected into lysis buffer using a micromanipulator under an inverted microscope. Single-cell reverse transcription and PCR were performed using the Signosis Single-Cell RT-PCR Assay Kit according to the manufacturer’s instructions. PCR products were resolved on 2% agarose gels and visualized by ethidium bromide staining. Primer sequences are listed in Supplementary Table 3.

### Nuclear and cytoplasmic RNA fractionation

Nuclear and cytoplasmic fractions of primary cultured mouse DRG neurons were isolated using the PARIS Kit (Invitrogen) according to the manufacturer’s instructions. Briefly, neurons were rinsed with ice-cold PBS, lysed in fractionation buffer, incubated on ice for 30 min, and centrifuged at 15,000 × g for 15 min at 4°C. RNA was extracted from each fraction, and gene/transcript expression was quantified by qRT-PCR as described above.

### *In vitro* RNA stability assay

*Nt3* mRNA stability was assessed by treating cultured cells with actinomycin D (ActD; 5 μg/mL) to inhibit transcription. Cells were harvested at 0, 0.5, 1, 2, and 4 h, and *Nt3* mRNA levels were quantified by qRT-PCR. Transcript abundance was normalized to the 0-h level and expressed as the percentage of RNA remaining.

### BaseScope® *in situ* hybridization and co-immunohistochemistry assays

Mice were perfused with ice-cold 4% paraformaldehyde (PFA), and bilateral L3/L4 DRGs were post-fixed for 4–6 h at 4°C and cryoprotected in 30% sucrose overnight. Human lumbar DRGs (AnaBios) were fixed in 4% PFA overnight at 4°C and similarly cryoprotected. Tissues were embedded and sectioned at 20 μm using a cryostat.

BaseScope in situ hybridization (ISH) was performed using the RED Detection Reagent Kit (ACD) according to the manufacturer’s protocol, with minor modifications. Sections were pretreated with hydrogen peroxide and Protease Plus, then hybridized with mouse/human *cRps* probes (REF: 713121, ACD) at 40°C for 2 days. Signal amplification was performed using AMP 1–8, followed by Fast RED detection (RED-B, 1:60). *Ppib* and bacterial *DapB* probes served as positive and negative controls, respectively.

Following ISH, sections were immunostained to identify neuronal and non-neuronal populations using antibodies against β-tubulin III (1:200; EMD Millipore), glutamine synthetase (GS; 1:200; Sigma-Aldrich), NF200 (1:100; Sigma), calcitonin gene-related peptide (CGRP; 1:100; Abcam), or biotinylated isolectin B4 (IB4; 1:100; Sigma). Sections were blocked with 4% normal donkey serum and 0.3% Triton X-100, incubated with primary antibodies overnight at 4°C and appropriate fluorescent secondary antibodies (1:200; Jackson ImmunoResearch) for 1 h at room temperature, then mounted with DAPI-containing VECTASHIELD.

Experiments included at least three independent replicates. Images were acquired using a Leica DMI4000 fluorescence microscope and analyzed with ImageJ. Cells containing ≥3 *cRps* ISH particles were considered *cRps*-positive. The percentage of *cRps*-positive neurons among β-tubulin III-positive neurons, *cRps* particles per positive neuron, and nuclear versus cytoplasmic particle counts were quantified ^43,44^.

### Immunofluorescence staining of PGP9.5 in hind paw skin

Mouse hind-paw plantar skin was fixed in 4% paraformaldehyde at 4°C, cryoprotected, embedded in OCT, and cryosectioned. Sections were permeabilized, blocked, and incubated overnight at 4°C with anti-PGP9.5 antibody (1:1,000; Sigma-Aldrich), followed by fluorescent secondary antibody and DAPI staining. Images were acquired under identical settings, and intraepidermal nerve fiber (IENF) density was quantified as the number of PGP9.5-positive fibers crossing the dermal–epidermal junction per unit epidermal length.

### Northern blotting analysis

A digoxigenin (DIG)-labeled cRNA probe targeting the mouse *cRps* back-splice junction was synthesized from mouse DRG cDNA using T7 RNA polymerase and DIG-RNA labeling mix (Roche), then DNase I-treated and purified. Primer sequences are listed in Supplementary Table 3.

Northern blotting was performed using the NorthernMax Kit (Invitrogen). Total RNA (2 μg for CAD cells; 30 μg for mouse DRGs, with or without RNase R treatment) was denatured, separated on 2% agarose gels, and transferred to positively charged nylon membranes. After UV crosslinking, membranes were hybridized overnight at 68°C with the DIG-labeled probe, washed with SSC buffer, and developed using CDP-Star substrate. Signals were visualized with a ChemiDoc XRS system and quantified using Image Lab (Bio-Rad).

### Western blotting analysis

Tissues were homogenized in ice-cold lysis buffer containing protease and phosphatase inhibitors. Equal amounts of protein were separated by SDS-PAGE and transferred to PVDF membranes. After blocking with 5% nonfat milk in TBST for 2 h, membranes were incubated overnight at 4°C with primary antibodies against NT3 (1:250; Alomone Labs), PUM2 (1:2,000; Abcam), actin (1:2,500; Santa Cruz Biotechnology), phospho-ERK1/2 (Thr202/Tyr204; 1:800; Cell Signaling Technology), Iba1 (1:1,000; Cell Signaling Technology), ERK1/2 (1:800; Cell Signaling Technology), GFAP (1:1,000; Cell Signaling Technology), or GAPDH (1:2,000; Santa Cruz Biotechnology) at the indicated dilutions. Following incubation with HRP-conjugated secondary antibodies, signals were developed using Clarity Western ECL Substrate (Bio-Rad) and visualized with a ChemiDoc XRS system. Band intensities were quantified using Image Lab, with GAPDH or β-actin as loading controls.

### *In vitro* RNA transcription, circularization, and purification

*In vitro* transcription and circularization were performed as previously described ^45^, with minor modifications. Full-length *cRps* RNA was transcribed from T7-DNA templates using the RiboMAX Large RNA Production System (Promega) and Biotin RNA Labeling Mix (Roche), with 400 mM GMP, at 37°C for 4 h. After DNase I treatment (30 min, 37°C), RNA was purified using Micro Bio-Spin P-30 columns (Bio-Rad). Linear RNA was circularized with T4 RNA ligase 1 (NEB) overnight at 16°C. Circular and linear products were separated on a 2% denaturing agarose gel, visualized by ethidium bromide staining, excised, and purified using the Zymoclean Gel RNA Recovery Kit.

### Comprehensive identification of RNA-binding proteins by mass spectrometry assay

RNA pull-down assays were performed using the Pierce Magnetic RNA-Protein Pull-Down Kit (Thermo Fisher Scientific). Biotinylated full-length sense *cRps* (2 μg) and linear *mRps* (2 μg; negative control) were synthesized, purified, and incubated with DRG lysates overnight at 4°C. RNA–protein complexes were captured using streptavidin magnetic beads, and bound proteins were eluted and separated by SDS-PAGE. Proteins were identified by LC-MS/MS using a Q Exactive mass spectrometer, with spectra searched against the *Mus musculus* Swiss-Prot database using MASCOT in Proteome Discoverer. The interaction between *cRps* and PUM2 was validated by Western blotting with an anti-PUM2 antibody.

### Luciferase reporter assay

The full-length *Nt3* mRNA 3′-UTR was PCR-amplified and cloned into the KpnI and NheI sites of the psiCHECK vector (Promega), as previously described ^43,44^. Recombinant clones were verified by sequencing. CAD cells were co-transfected with *cRps*- or *Pum2*-expressing plasmids (300 ng), reporter constructs with or without the *Nt3* 3′-UTR (300 ng), and pRL-TK (10 ng) using Lipofectamine 3000. After 48 h, luciferase activity was measured using the Dual-Luciferase Reporter Assay System (Promega). Relative reporter activity was calculated by normalizing firefly to Renilla luciferase activity. Experiments were independently repeated three times.

### RNA Immunoprecipitation (RIP) Assay

RNA immunoprecipitation (RIP) was performed using the Magna RIP Kit (Millipore) according to the manufacturer’s instructions. DRGs were homogenized in ice-cold PBS, centrifuged, and lysed in RIP buffer containing protease and RNase inhibitors. Lysates were incubated overnight at 4°C with magnetic protein A/G beads conjugated to anti-PUM2 antibody (2 μg; Abcam) or control IgG. After six washes, immunoprecipitated RNA was purified by proteinase K treatment and phenol/chloroform extraction. Input samples (10–20%) were retained, and *cRps*, the *Nt3* 3′-UTR, and *mRps* intronic sequences were quantified by RT-qPCR. Primer sequences are listed in Supplementary Table 3.

### RNA sequencing

Four weeks after STZ or vehicle treatment, ipsilateral L3/L4 DRGs from mice pre-injected with AAV5-scrambled shRNA or AAV5-*cRps* shRNA were harvested and pooled (two mice/sample). Total RNA (1.2 μg/sample) underwent rRNA depletion using the Ribo-Zero kit (Illumina), followed by stranded RNA library preparation without poly-A selection using the TruSeq Stranded Total RNA Kit. Experiments were performed in triplicate, and libraries were sequenced on an Illumina HiSeq 2500 platform (2 × 150-bp paired-end), yielding at least 60 million reads per sample.

### Bioinformatics analysis

Sequencing reads were quality-filtered, depleted of residual rRNA using RNAcentral, and aligned to the mouse reference genome using STAR. Candidate circRNAs were identified by back-splice junction reads and annotated by genomic location and length. Differential expression was analyzed using DESeq in R with default parameters. Potential PUM2-binding sites in the *Nt3* 3′-UTR and *cRps* were predicted using IntaRNA, LncRNA, TargetScan, RBPSuite, and LncTar.

### Statistical analysis

Animals and cells were randomly assigned to experimental groups. Sample sizes were based on pilot studies, previous reports, and power analyses (90% power, α = 0.05). Data are presented as mean ± SEM from at least three independent experiments. Normality was assumed but not formally tested. Statistical analyses included two-tailed paired Student’s *t*-tests or one-, two-, or three-way ANOVA, followed by Tukey’s post hoc test when appropriate. Analyses were performed using GraphPad Prism, with *P* < 0.05 considered significant.

### Data availability

The raw data are available upon request for academic research purposes, subject to access restrictions. Researchers may request access by contacting the corresponding author, Dr. Yuan-Xiang Tao. Access is contingent upon approval by the applicant’s home institution and execution of a data-sharing agreement. Requests will be acknowledged within four weeks, and a decision regarding access will be made within 12 weeks, subject to institutional review and completion of the required agreement. Approved investigators will be granted access to the requested data for 12 months. Extensions may be considered upon approval by the corresponding author and the applicant’s institution. The full, uncropped gel and blot images are provided with this paper.

## ACKNOWLEDGMENTS

This work was supported by NIH grants R01DK142785 and R61NS145978 to Y.X.T.

## AUTHOR CONTRIBUTIONS

Y.X.T. conceived the project and supervised all experiments. X.L., C.H., G.G., B.W., and H.H.Z. contributed substantially to study design, execution, and data acquisition, analysis, and interpretation. X.L., C.C., D.D., B.Z., M.A.F., T.B., J.M., and X.L. prepared animal models and conducted behavioral experiments. X.L., C.H., G.G., B.W., H.H.Z., X.F., D.S., R.M., L.Z., B.Z., J.X, and B.J. performed molecular, biochemical, and morphological experiments. X.L., C.H., H.H.Z., and J.X. conducted bioinformatic analyses, while X.L., C.H., and X.F. performed mass spectrometry and data analysis. G.G. and X.S.Z. assisted with tissue collection from NOD/ShiLtJ mice. S.D. collected human DRGs and performed DRG cultures. X.L., C.H., G.G., B.W., H.H.Z., X.F., D.S., R.M., L.Z., B.Z., M.A.F., T.B., J.C., and Y.X.T. analyzed the data. X.L., B.W, and Y.X.T. drafted the manuscript, and Y.X.T. finalized it. All authors reviewed and discussed the manuscript.

## COMPETING FINANCIAL INTERESTS

The authors declare no compete financial interests.

## ETHICS AND CONSENT TO PARTICIPATE DECLARATIONS

Not applicable.

## SUPPLEMENTARY MATERIALS

Figs. S1 to S21 Table S1-S3

## Notes

### Competing Interest Statement

The authors have declared no competing interest.

## References

1. Campbell, J.N. & Meyer, R.A. Mechanisms of neuropathic pain. Neuron 52, 77–92 (2006).

2. van, H.O., Austin, S.K., Khan, R.A., Smith, B.H. & Torrance, N. Neuropathic pain in the general population: a systematic review of epidemiological studies. Pain 155, 654–662 (2014).

3. Feldman, E.L., et al. Diabetic neuropathy. Nat Rev Dis Primers 5, 42 (2019).

4. Waldfogel, J.M., et al. Pharmacotherapy for diabetic peripheral neuropathy pain and quality of life: A systematic review. Neurology 88, 1958–1967 (2017).

5. Jang, H.N. & Oh, T.J. Pharmacological and Nonpharmacological Treatments for Painful Diabetic Peripheral Neuropathy. Diabetes Metab J 47, 743–756 (2023).

6. Gibon, J. & Barker, P.A. Neurotrophins and Proneurotrophins: Focus on Synaptic Activity and Plasticity in the Brain. Neuroscientist 23, 587–604 (2017).

7. Teng, K.K., Felice, S., Kim, T. & Hempstead, B.L. Understanding proneurotrophin actions: Recent advances and challenges. Dev Neurobiol 70, 350–359 (2010).

8. Yano, H., Torkin, R., Martin, L.A., Chao, M.V. & Teng, K.K. Proneurotrophin-3 is a neuronal apoptotic ligand: evidence for retrograde-directed cell killing. J Neurosci 29, 14790–14802 (2009).

9. Blum, R. & Konnerth, A. Neurotrophin-mediated rapid signaling in the central nervous system: mechanisms and functions. Physiology (Bethesda) 20, 70–78 (2005).

10. Shang, H., Iyer, R., Wang, B. & Tao, Y.X. EXPRESS: Precursor signaling redefined: proneurotrophin-3, not neurotrophin-3, in primary sensory neurons as an instigator in neuropathic pain. Mol Pain, 17448069261460645 (2026).

11. Shang, H., et al. Contribution of proneurotrophin-3 to nerve trauma-induced neuropathic pain through promoting TrkC-mediated increase of CCL2 in primary sensory neurons. Brain Behav Immun, 106256 (2026).

12. Sharma, D., et al. Proneurotrophin-3 contributes to chemotherapy-induced neuropathic pain through TrkC-mediated CCL2 elevation in DRG neurons. EMBO Rep (2025).

13. Chen, L.L. The biogenesis and emerging roles of circular RNAs. Nat. Rev. Mol. Cell Biol 17, 205–211 (2016).

14. Xia, S., et al. Comprehensive characterization of tissue-specific circular RNAs in the human and mouse genomes. Brief Bioinform 18, 984–992 (2017).

15. Rybak-Wolf, A., et al. Circular RNAs in the Mammalian Brain Are Highly Abundant, Conserved, and Dynamically Expressed. Mol Cell 58, 870–885 (2015).

16. Gu, X., et al. The Roles of Circular RNAs in Ischemic Stroke through Modulating Neuroinflammation. J Integr Neurosci 23, 87 (2024).

17. Hu, T., Li, D., Fan, T., Zhao, X. & Chen, Z. Circular RNA PUM1 performs as a competing endogenous RNA of microRNA-340-5p to mediate DEAD-box helicase 5 to mitigate cerebral ischemia-reperfusion injury. Bioengineered 13, 11564–11578 (2022).

18. Feng, X., et al. Circular RNA circNrip1 Interacts with SYNCRIP to Promote Neuropathic Pain by Stabilizing Tlr2 mRNA in Primary Sensory Neurons. Adv Sci (Weinh), e19740 (2026).

19. Pan, Z., et al. MicroRNA-1224 Splicing CircularRNA-Filip1l in an Ago2-Dependent Manner Regulates Chronic Inflammatory Pain via Targeting Ubr5. J Neurosci 39, 2125–2143 (2019).

20. Zhang, S.B., et al. CircAnks1a in the spinal cord regulates hypersensitivity in a rodent model of neuropathic pain. Nat. Commun 10, 4119 (2019).

21. Zhang, H.H., et al. Circular RNA profile in diabetic peripheral neuropathy: analysis of coexpression networks of circular RNAs and mRNAs. Epigenomics 12, 843–857 (2020).

22. O’Brien, P.D., Sakowski, S.A. & Feldman, E.L. Mouse models of diabetic neuropathy. ILAR. J 54, 259–272 (2014).

23. Li, Y., et al. N(6)-Methyladenosine Demethylase FTO Contributes to Neuropathic Pain by Stabilizing G9a Expression in Primary Sensory Neurons. Adv. Sci. (Weinh. ) 7, 1902402 (2020).

24. Zhao, J.Y., et al. DNA methyltransferase DNMT3a contributes to neuropathic pain by repressing Kcna2 in primary afferent neurons. Nat. Commun 8, 14712 (2017).

25. Zhao, X., et al. A long noncoding RNA contributes to neuropathic pain by silencing Kcna2 in primary afferent neurons. Nat. Neurosci 16, 1024–1031 (2013).

26. Zhang, Y., et al. RNA-binding protein SYNCRIP contributes to neuropathic pain through stabilising CCR2 expression in primary sensory neurones. Br J Anaesth 133, 1028–1041 (2024).

27. Zhou, Y., et al. Circular RNA circ-FIRRE interacts with HNRNPC to promote esophageal squamous cell carcinoma progression by stabilizing GLI2 mRNA. Cancer Sci 114, 3608–3622 (2023).

28. Goldstrohm, A.C., Hall, T.M.T. & McKenney, K.M. Post-transcriptional Regulatory Functions of Mammalian Pumilio Proteins. Trends Genet 34, 972–990 (2018).

29. Das, A., Sinha, T., Shyamal, S. & Panda, A.C. Emerging Role of Circular RNA-Protein Interactions. Noncoding. RNA 7(2021).

30. Liu, C.X. & Chen, L.L. Circular RNAs: Characterization, cellular roles, and applications. Cell 185, 2016–2034 (2022).

31. Didangelos, T., Doupis, J. & Veves, A. Painful diabetic neuropathy: clinical aspects. Handb Clin Neurol 126, 53–61 (2014).

32. Tesfaye, S., et al. Painful diabetic peripheral neuropathy: consensus recommendations on diagnosis, assessment and management. Diabetes Metab Res Rev 27, 629–638 (2011).

33. White, E.K., Moore-Jarrett, T. & Ruley, H.E. PUM2, a novel murine puf protein, and its consensus RNA-binding site. RNA 7, 1855–1866 (2001).

34. Padmanabhan, K. & Richter, J.D. Regulated Pumilio-2 binding controls RINGO/Spy mRNA translation and CPEB activation. Genes Dev 20, 199–209 (2006).

35. Biber, K. & Boddeke, E. Neuronal CC chemokines: the distinct roles of CCL21 and CCL2 in neuropathic pain. Front Cell Neurosci 8, 210 (2014).

36. Scoles, D.R. & Pulst, S.M. Oligonucleotide therapeutics in neurodegenerative diseases. RNA. Biol 15, 707–714 (2018).

37. Wurster, C.D., et al. Intrathecal administration of nusinersen in adolescent and adult SMA type 2 and 3 patients. J Neurol 266, 183–194 (2019).

38. O’Neill, J., et al. Differences in mouse models of diabetes mellitus in studies of male reproduction. Int. J. Androl 33, 709–716 (2010).

39. Wen, C.H., et al. Effect of intrathecal NIS-lncRNA antisense oligonucleotides on neuropathic pain caused by nerve trauma, chemotherapy, or diabetes mellitus. Br J Anaesth 130, 202–216 (2023).

40. Coppey, L.J., Shevalye, H., Obrosov, A., Davidson, E.P. & Yorek, M.A. Determination of peripheral neuropathy in high-fat diet fed low-dose streptozotocin-treated female C57Bl/6J mice and Sprague-Dawley rats. J Diabetes Investig 9, 1033–1040 (2018).

41. Parilla, J.H., Willard, J.R., Barrow, B.M. & Zraika, S. A Mouse Model of Beta-Cell Dysfunction as Seen in Human Type 2 Diabetes. J Diabetes Res 2018, 6106051 (2018).

42. Ford, Z.K., et al. Cannabinoid Receptor 1 Expression in Human Dorsal Root Ganglia and CB13-Induced Bidirectional Modulation of Sensory Neuron Activity. Front Pain Res (Lausanne) 2, 721332 (2021).

43. Du, S., et al. A nerve injury-specific long noncoding RNA promotes neuropathic pain by increasing Ccl2 expression. J Clin Invest 132, e153563 (2022).

44. Wang, B., et al. A sensory neuron-specific long non-coding RNA reduces neuropathic pain by rescuing KCNN1 expression. Brain 146, 3866–3884 (2023).

45. Liu, C.X., et al. Structure and Degradation of Circular RNAs Regulate PKR Activation in Innate Immunity. Cell 177, 865–880 e821 (2019).

