## Supplementary Figures and Tables for "Hyperglycemia-responsive circRNA drives diabetic neuropathic pain by competing for PUM2 binding and promoting proNT3/CCL2 in DRG neurons"

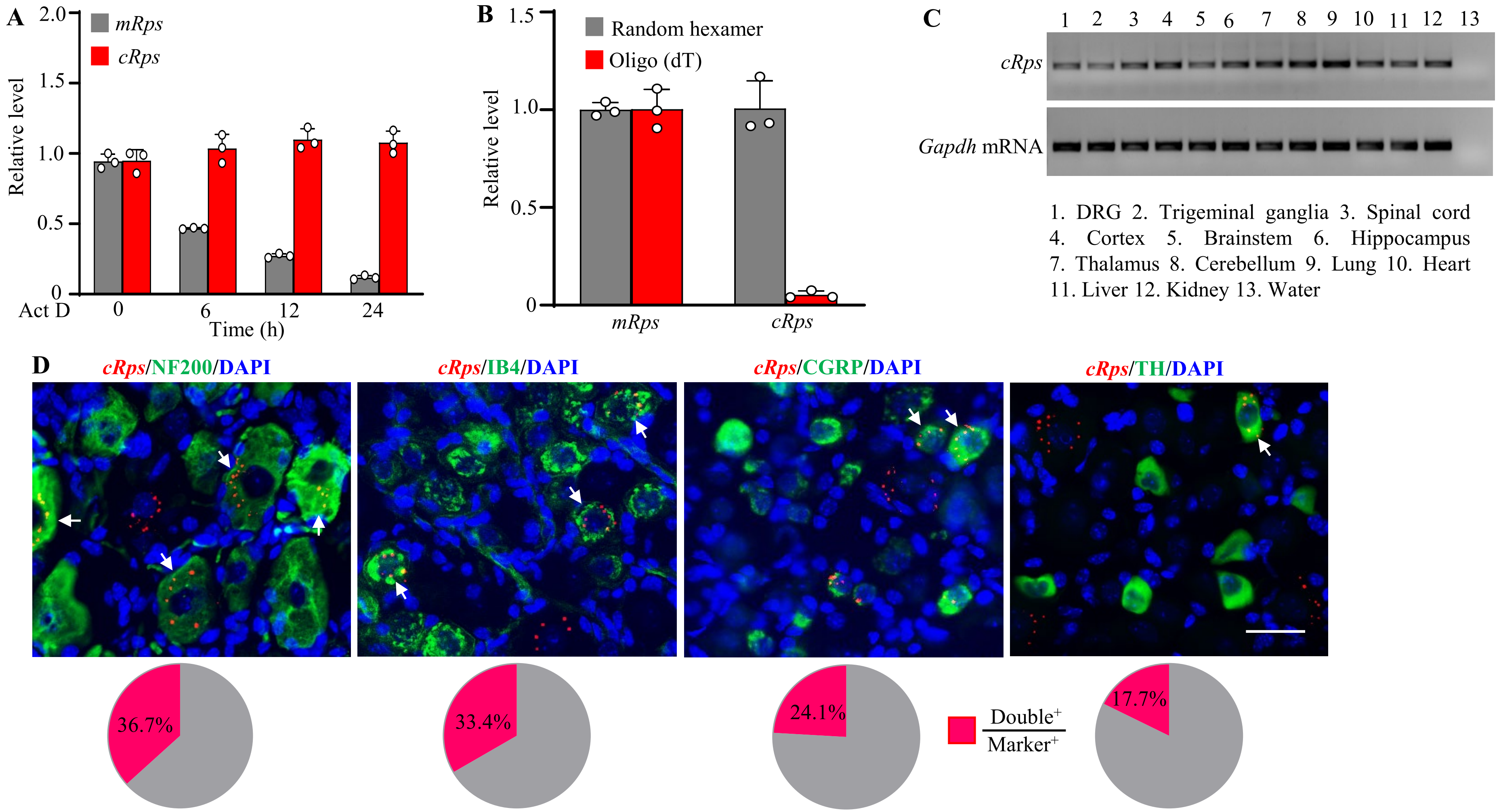

**Supplementary Fig. 1** (A) Expression levels of *cRps* and *mRps* in the cultured DRG neurons at different hours (h) after treatment with actinomycin D (ActD). n = 3 replicates/time point. (B) Expression levels of *cRps* and *mRps* in cultured DRG neurons were analyzed by quantitative PCR following reverse transcription (RT) using either random hexamer primers (RH) or oligo (dT) primers (OD). n = 3 replicates/RT. (C) Expression level of *cRps* in different tissues as indicated in normal mice. *Gapdh* mRNA is used as an internal control. (D) Co-localization (arrows) of *cRps* with neurofilament 200 (NF200), isolectin B4 (IB4), calcitonin gene-related peptide (CGRP) and tyrosine hydroxylase (TH) in DRG neurons of mice 4 weeks after STZ injection. *n* = 3 mice. Scale bar: 40 µm.

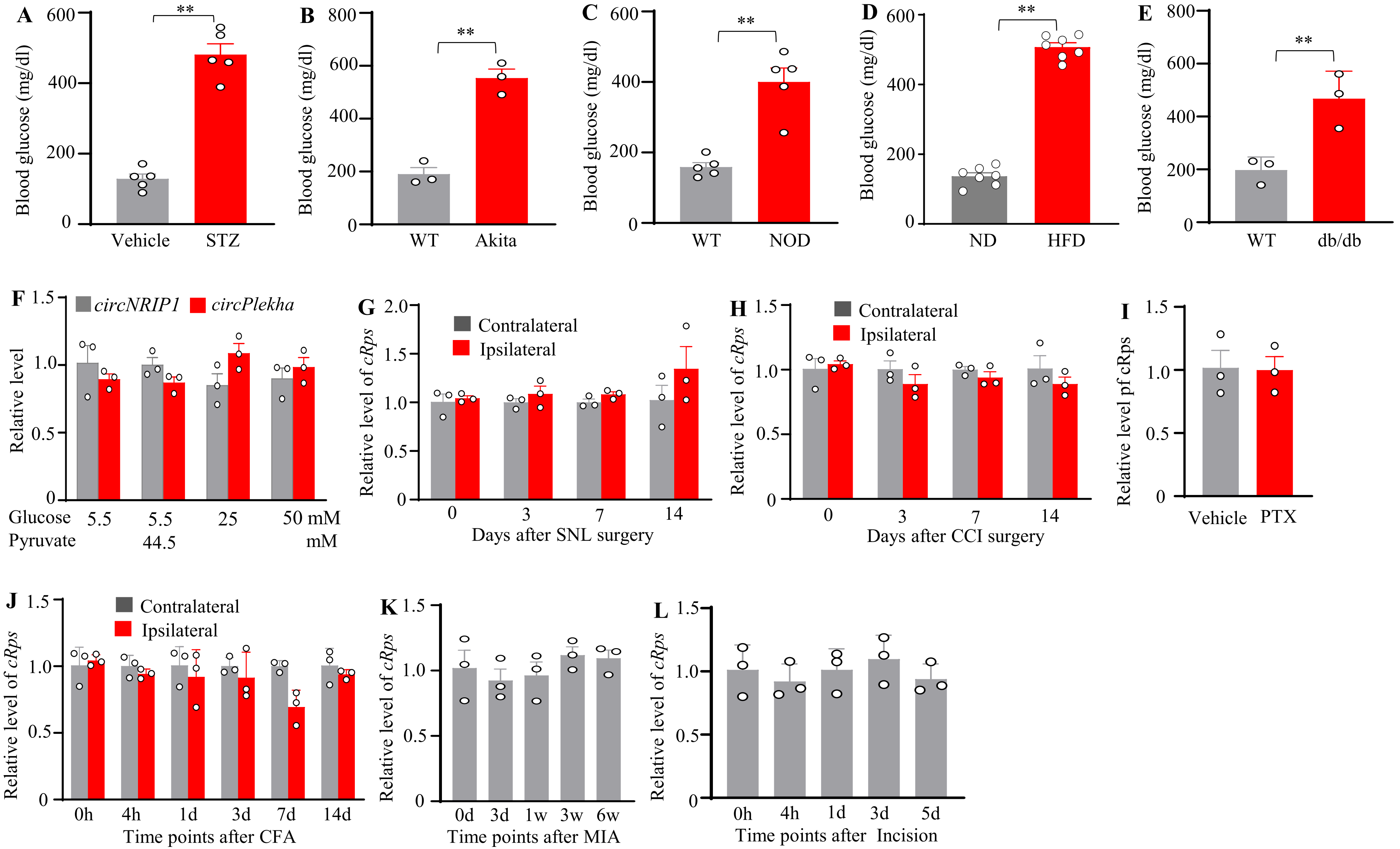

A
Blood glucose (mg/dl)
**
Vehicle
STZ
B
Blood glucose (mg/dl)
**
WT
Akita
C
Blood glucose (mg/dl)
**
WT
NOD
D
Blood glucose (mg/dl)
**
ND
HFD
E
Blood glucose (mg/dl)
**
WT
db/db
F
circNRIP1
circPlekha
Relative level
Glucose 5.5 5.5 25 50 mM
Pyruvate 44.5 mM
G
Contralateral
Ipsilateral
Relative level of cRps
Days after SNL surgery
H
Contralateral
Ipsilateral
Relative level of cRps
Days after CCI surgery
I
Relative level pf cRps
Vehicle
PTX
J
Contralateral
Ipsilateral
Relative level of cRps
0h 4h 1d 3d 7d 14d
Time points after CFA
K
Relative level of cRps
0d 3d 1w 3w 6w
Time points after MIA
L
Relative level of cRps
0h 4h 1d 3d 5d
Time points after Incision

**Supplementary Fig. 2.** (A, B) Expression levels of *cRps* (A) and *mRps* (B) in the ipsilateral L3/4 dorsal horn 2, 3, 4 and 6 weeks following STZ or vehicle injection. n = 3-6 mice/time point. Two-way ANOVA followed by Tukey's post hoc test. (C) Expression level of *cRps* in different tissues as indicated in mice 4 weeks following STZ or vehicle injection. $n$ = 5 mice/time point/group. Two-tailed paired Student's t test. (D) Expression level of *cRps* in the trigeminal ganglion from the Akita mice and the corresponding wild type (WT) mice. n = 3 mice/genotyping. Two-tailed paired Student's t test. (E-H) Behavioral tests showed the increases in paw withdraw frequency (PWF) to 0.07-g (E) and 0.4-g (F) von Frey filament stimuli and reductions in pain withdrawal latency (PWL) to heat (G) and cold (H) stimuli on the right and left sides at the different weeks as indicated after STZ or vehicle injection. n = 12 mice/group. ** $P < 0.01$ verse the vehicle group at the corresponding time points, two-way repeated-measures ANOVA followed by Tukey's post hoc test. (I-L) Behavioral tests showed the increases in paw withdraw frequency (PWF) to 0.07-g (I) and 0.4-g (J) von Frey filament stimuli and reductions in pain withdrawal latency (PWL) to heat (K) and cold (L) stimuli on the right and left sides at the different weeks as indicated after the initiation of normal diet (ND) or high-fat diet (HFD). n = 7 mice/group. **$P < 0.05$, ** $P < 0.01$ verse the vehicle group at the corresponding time points, two-way repeated-measures ANOVA followed by Tukey's post hoc test.

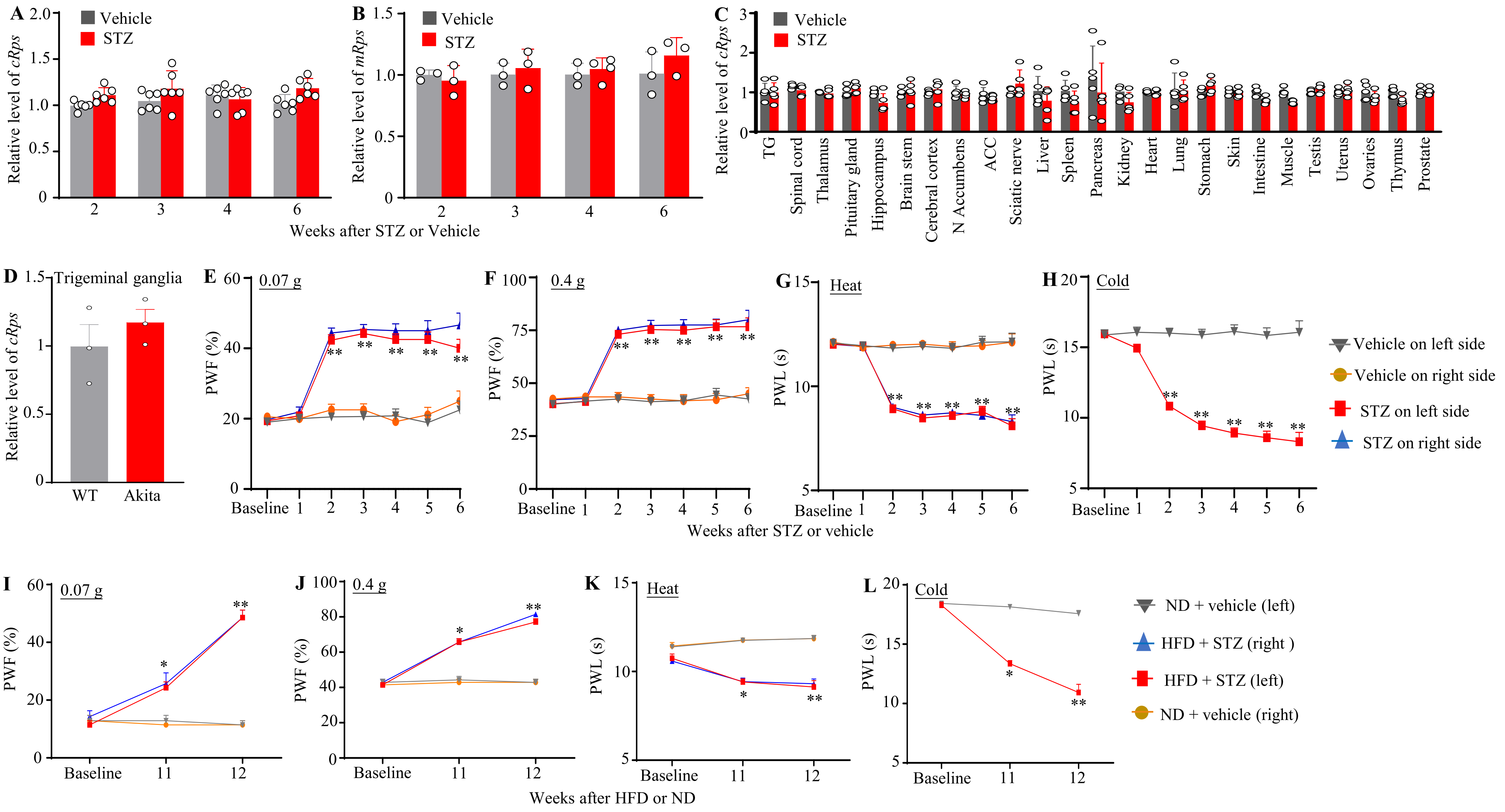
A
B
C
Vehicle
STZ
Relative level of cRps
Relative level of mRps
Weeks after STZ or Vehicle
TG
Spinal cord
Thalamus
Pituitary gland
Hippocampus
Brain stem
Cerebral cortex
N Accumbens
ACC
Sciatic nerve
Liver
Spleen
Pancreas
Kidney
Heart
Lung
Stomach
Skin
Intestine
Muscle
Testis
Uterus
Ovaries
Thymus
Prostate
D
Trigeminal ganglia
WT
Akita
E
F
G
H
0.07 g
0.4 g
Heat
Cold
PWF (%)
PWL (s)
Baseline
Weeks after STZ or vehicle
Vehicle on left side
Vehicle on right side
STZ on left side
STZ on right side
I
J
K
L
Weeks after HFD or ND
ND + vehicle (left)
HFD + STZ (right )
HFD + STZ (left)
ND + vehicle (right)

**Supplementary Fig.3**. (A-E) Blood glucose levels in STZ-induced T1DM male mice and the corresponding control vehicle-treated male mice (4 weeks after STZ or vehicle injection; A), Akita male mice and the corresponding wild type (WT) male mice (B), NOD (non-obese diabetic) male mice and the corresponding WT male mcie (C), HFD-induced T2DM male mice and the corresponding control ND-fed male mice (15 weeks after feeding; D), and db/db male mice and the corresponding WT male mice (E). n = 3-7 mice/treatment or genotyping. $^{**}P < 0.01$, two-tailed paired Student's $t$ test. (F) Expression levels of *circNRIP1* and *circPlekha* in the cultured DRG neurons exposued to the different concentrations of glucose. n = 3 replicates/concentration. One-way ANOVA followed post hoc Tukey's test. (G-L) Expression level of *cRps* in the ipsilateral and contralateral L4 DRGs on indicated days after SNL (G), in the ipsilateral and contralateral L3/4 DRGs on indicated days after CCI (H), in the bilateral L3/4 DRGs 14 days after paclitaxel (PTX) injection (I), in the ipsilateral and contralateral L3/4 DRGs on the indicated days after CFA injection (J), in the ipsilateral L3/4 DRGs on the indicated time points after MIA injection (K) and in the ipsilateral L3/4 DRGs on the indicated time points after paw incision (L). n = 6 mice/model/time point. One-wan ANOVA followed post hoc Tukey's test.

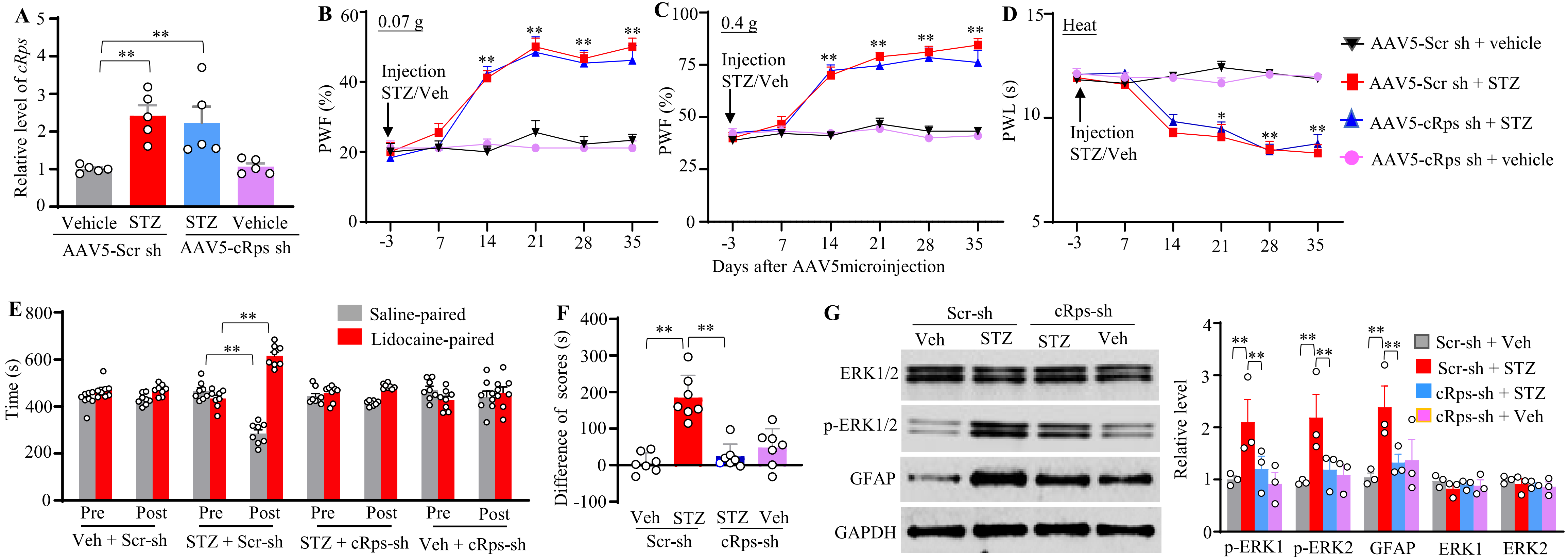

**Supplementary Fig. 4.** (A) Expression level of *cRps* in contralateral L3/L4 DRGs from STZ- or vehicle-treated male mice 35 days after unilateral microinjection of AAV5 expressing a *cRps*-specific shRNA (AAV5-cRps sh) or a control scrambled shRNA (AAV5-Scr sh) into the L3/L4 DRGs. $n = 5$ replicates (2 mice/replicate)/group. $^{**}P < 0.01$, two-way ANOVA followed by Tukey's post hoc test. (B-D) Effect of post-microinjection of AAV5-cRps sh or AAV5-Scr sh into unilateral L3/L4 DRGs 3 days after STZ or vehicle injection on paw withdrawal frequency (PWF) to 0.07-g (B) and 0.4-g (C) von Frey filament stimuli and paw withdrawal latency (PWL) to heat stimulation (D) on the indicated days after viral microinjection on the contralateral side in male mice. $n = 9$ mice/group. $^{*}P < 0.05$, $^{**}P < 0.01$ versus the vehicle plus AAV5-Scr sh group at the corresponding time points; three-way repeated-measures ANOVA followed by Tukey's post hoc test. (E, F) Effect of post-microinjection of AAV5-cRps sh (cRps-sh) or AAV5-Scr sh (Scr-sh) into unilateral L3/L4 DRGs 3 days after STZ or vehicle injection on spontaneous ongoing pain, assessed using the conditioned place preference (CPP) paradigm 28 days after viral microinjection in male mice. Pre, preconditioning; Post, post-conditioning. $n = 7$-$8$ mice/group. $^{**}P < 0.01$, two (F)- or three (E)-way repeated-measures ANOVA followed by Tukey's post hoc test. (G) Effects of unilateral L3/L4 DRG microinjection of AAV5-cRps-sh (cRps-sh) or AAV5-Scr sh (Scr-sh) on levels of p-ERK1/2, ERK1/2, and GFAP in the ipsilateral L3/4 dorsal horn 35 days after viral microinjection in male mice. $n = 3$ mice/group. $^{**}P < 0.01$, 2-way ANOVA followed by Tukey's post hoc test.

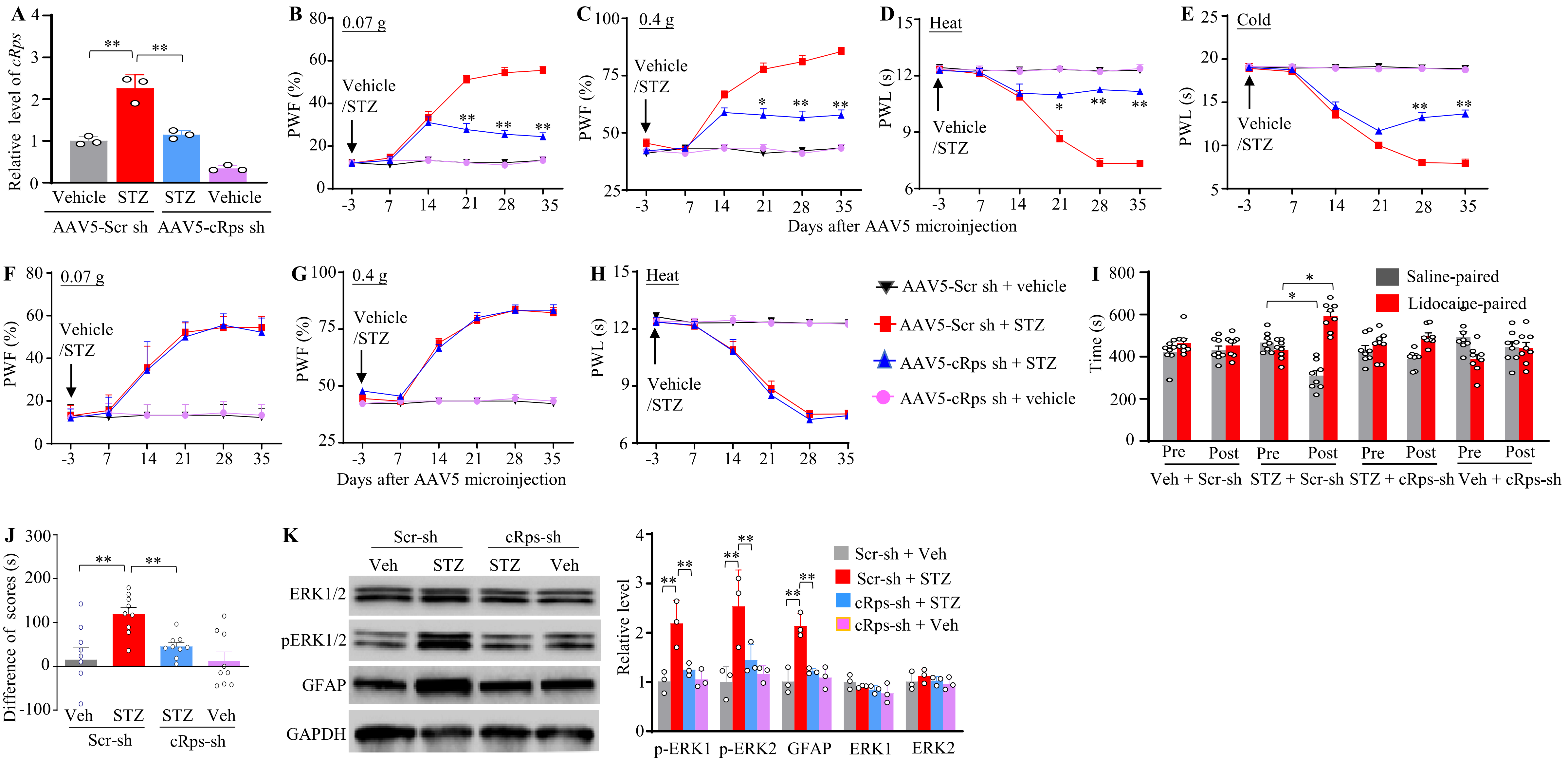
A
Relative level of cRps
Vehicle STZ STZ Vehicle
AAV5-Scr sh AAV5-cRps sh
B 0.07 g
C 0.4 g
D Heat
E Cold
PWF (%)
PWL (s)
Vehicle /STZ
Days after AAV5 microinjection
F 0.07 g
G 0.4 g
H Heat
AAV5-Scr sh + vehicle
AAV5-Scr sh + STZ
AAV5-cRps sh + STZ
AAV5-cRps sh + vehicle
I
Time (s)
Saline-paired
Lidocaine-paired
Pre Post
Veh + Scr-sh STZ + Scr-sh STZ + cRps-sh Veh + cRps-sh
J
Difference of scores (s)
Veh STZ STZ Veh
Scr-sh cRps-sh
K
Scr-sh cRps-sh
Veh STZ STZ Veh
ERK1/2
pERK1/2
GFAP
GAPDH
Relative level
Scr-sh + Veh
Scr-sh + STZ
cRps-sh + STZ
cRps-sh + Veh
p-ERK1 p-ERK2 GFAP ERK1 ERK2

**Supplementary Fig. 5**. **Blocking *cRps* upregulation in the DRG alleviates established diabetic neuropathic pain in female mice with T1DM**. (A) Expression level of *cRps* in the ipsilateral L3/L4 DRGs from STZ- or vehicle-treated female mice 35 days after unilateral microinjection of AAV5 expressing a *cRps*-specific shRNA (AAV5-cRps sh) or a scrambled control shRNA (AAV5-Scr sh) into the L3/L4 DRGs. $n = 6$ mice. $^{**}P < 0.01$, two-way ANOVA followed by Tukey's post hoc test. (B–H) Effect of post-microinjection of AAV5-cRps sh or AAV5-Scr sh into unilateral L3/L4 DRGs 3 days after STZ or vehicle injection on paw withdrawal frequency (PWF) to 0.07-g (B and F) and 0.4-g (C and G) von Frey filament stimuli and paw withdrawal latency (PWL) to heat (D and H) and cold (E) stimuli on the indicated days after viral microinjection on the ipsilateral (B-E) and contralateral (F-H) sides. $n = 9$ mice/group. $^{*}P < 0.05$, $^{**}P < 0.01$ versus the STZ plus AAV5-Scr sh group at the corresponding time points; three-way repeated-measures ANOVA followed by Tukey's post hoc test. (I, J) Effect of post-microinjection of AAV5-cRps sh (cRps-sh) or AAV5-Scr sh (Scr-sh) into unilateral L3/L4 DRGs 3 days after STZ or vehicle injection on spontaneous ongoing pain, assessed using the conditioned place preference (CPP) paradigm 28 days after viral microinjection in female mice. Pre, preconditioning; Post, post-conditioning. $n = 7\text{-}8$ mice/group. $^{**}P < 0.01$, two (J)- or three (I)-way repeated-measures ANOVA followed by Tukey's post hoc test. (K) Effects of unilateral L3/L4 DRG microinjection of AAV5-cRps-sh (cRps-sh) or AAV5-Scr sh (Scr-sh) on levels of p-ERK1/2, ERK1/2, and GFAP in the ipsilateral L3/4 dorsal horn 35 days after viral microinjection in female mice. $n = 3$ mice/group. $^{**}P < 0.01$, 2-way ANOVA followed by Tukey's post hoc test. .

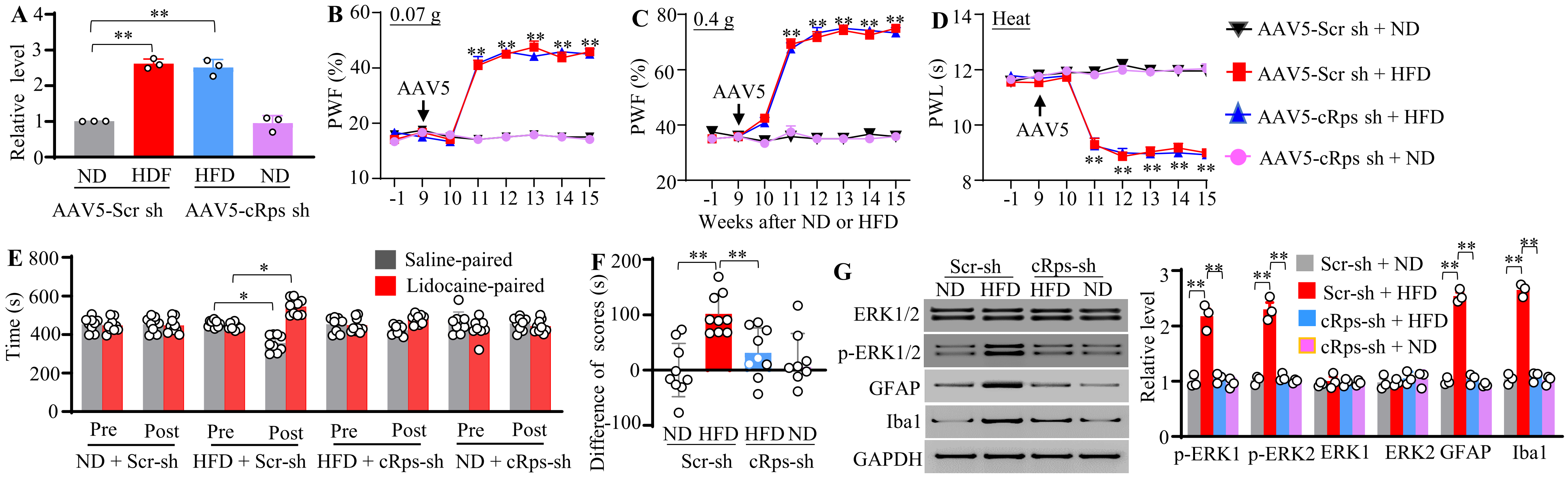

**Supplementary Fig. 6.** (A) Expression level of *cRps* in the contralateral L3/L4 DRGs from male mice fed a normal diet (ND) or high-fat diet (HFD) for 15 weeks. AAV5-cRps sh or AAV5-Scr sh was unilaterally microinjected into the L3/L4 DRGs 9 weeks after the initiation of dietary feeding. *n* = 3 replicates (2 mice/replicate)/group. $^{**}P < 0.01$, two-way ANOVA followed by Tukey's post hoc test. (B-D) Effect of post-microinjection of AAV5-cRps sh or AAV5-Scr sh into unilateral L3/L4 DRGs 9 weeks after the initiation of ND or HFD feeding on paw withdrawal frequency (PWF) to 0.07-g (B) and 0.4-g (C) von Frey filament stimuli and paw withdrawal latency (PWL) to heat (D) stimulation on the indicated weeks after viral microinjection on the contralateral side. *n* = 12 mice/group. $^{**}P < 0.01$ versus the ND plus AAV5-Scr sh group at the corresponding time points; three-way repeated-measures ANOVA followed by Tukey's post hoc test. (E, F). Effect of post-microinjection of AAV5-cRps sh (cRps-sh) or AAV5-Scr sh (Scr-sh) into unilateral L3/L4 DRGs 9 weeks after the initiation of ND or HFD feeding on spontaneous ongoing pain, assessed using the conditioned place preference (CPP) paradigm 14 weeks after the initiation of ND or HFD feeding in male mice. Pre, preconditioning; Post, post-conditioning. *n* = 9 mice/group. $^{*}P < 0.05$, $^{**}P < 0.01$, two- (F) or three-(E) way ANOVA followed by Tukey's post hoc test. (G) Effects of unilateral L3/L4 DRG microinjection of AAV5-cRps-sh (cRps-sh) or AAV5-Scr sh (Scr-sh) on levels of p-ERK1/2, ERK1/2, and GFAP in the ipsilateral L3/4 dorsal horn 15 weeks after the initiation of ND or HFD feeding in male mice. n = 3 mice)/treatment. $^{**}P < 0.01$ by two-way ANOVA followed by Tukey post hoc test.

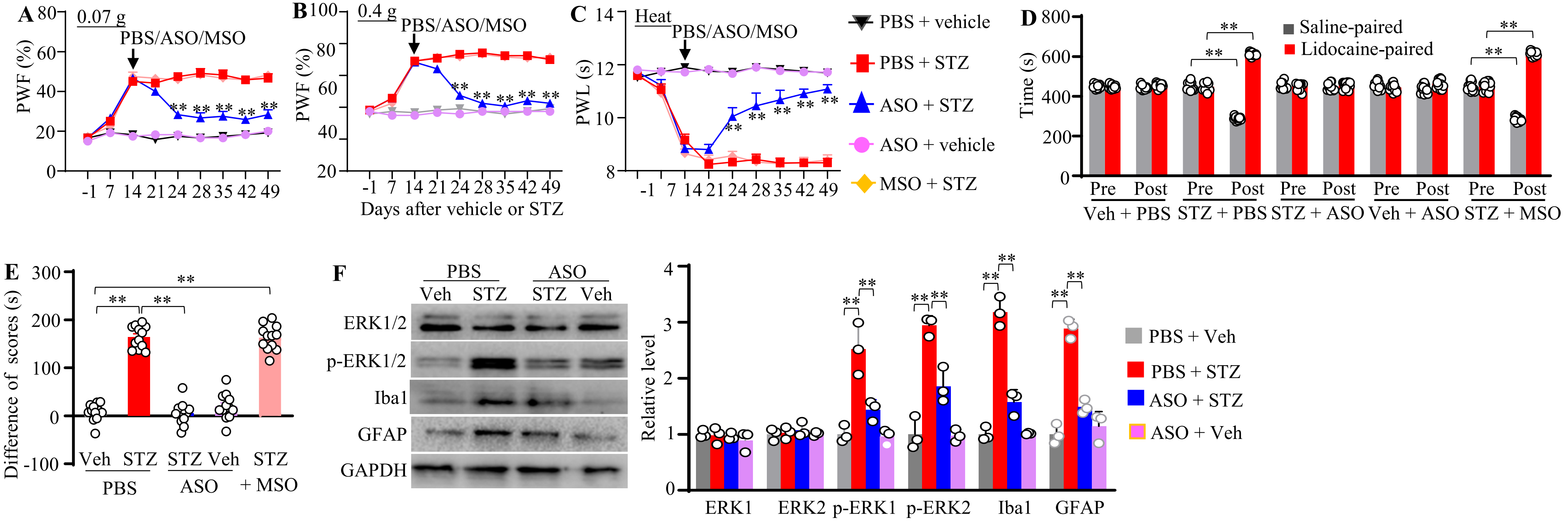

**Supplementary Fig. 7**. (A-C) Effect of intrathecal administration of PBS, ASO (100 µg), or MSO (100 µg) 2 weeks after STZ or vehicle injection on paw withdrawal frequency (PWF) to 0.07-g (A) and 0.4-g (B) von Frey filament stimuli and PWL to heat (C) stimulation on the indicated days after STZ or vehicle injection on the right side. *n* = 12 mice/group. $^{**}P < 0.01$ versus the STZ plus PBS group at the corresponding time points; three-way repeated-measures ANOVA followed by Tukey's post hoc test. (D, E) Effect of intrathecal administration of PBS, ASO (100 µg), or MSO (100 µg) 2 weeks after STZ or vehicle injection on spontaneous ongoing pain, assessed using the conditioned place preference (CPP) paradigm 42 days after STZ or vehicle injection in male mice. Pre, preconditioning; Post, post-conditioning. *n* = 12 mice /group. $^{**}P < 0.01$, two- (E) or three-(D) way repeated-measures ANOVA followed by Tukey's post hoc test. (F) Effect of intrathecal administration of PBS or ASO (100 µg) 2 weeks after STZ or vehicle injection on levels of p-ERK1/2, ERK1/2, Iba1, and GFAP in the ipsilateral L3/4 dorsal horn 49 days after STZ or vehicle injection. *n* = 3 mice/group.$^{**}P < 0.01$, 2-way ANOVA followed by Tukey's post hoc test.

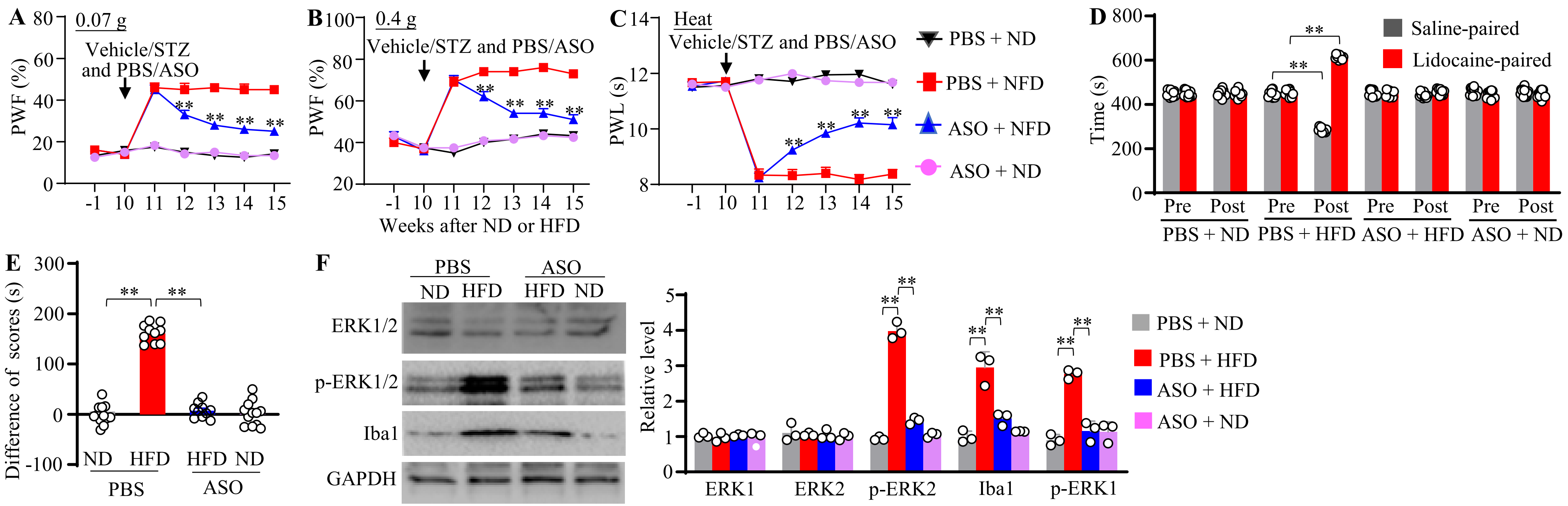

**Supplementary Fig. 8**. (A-C) Effect of intrathecal administration of PBS or ASO (100 µg) 10 weeks after initiation of ND or HFD feeding on paw withdrawal frequency (PWF) to 0.07-g (A) and 0.4-g (B) von Frey filament stimuli and paw withdrawal latency (PWL) to heat (C) stimulation on the indicated weeks after initiation of ND or HFD feeding on the right side. *n* = 12 mice/group. ***P* < 0.01 versus the HFD plus PBS group at the corresponding time points; three-way repeated-measures ANOVA followed by Tukey's post hoc test. (D, E) Effect of intrathecal administration of PBS or ASO (100 µg) 10 weeks after initiation of ND or HFD feeding on spontaneous ongoing pain, assessed using the conditioned place preference (CPP) paradigm 14 weeks after initiation of ND or HFD feeding in male mice. Pre, preconditioning; Post, post-conditioning. *n* = 12 mice /group. ***P* < 0.01, two- (E) or three-(D) way repeated-measures ANOVA followed by Tukey's post hoc test. (F) Effect of intrathecal administration of PBS or ASO (100 µg) 10 weeks after initiation of ND or HFD feeding on levels of p-ERK1/2, ERK1/2 and Iba1 in the ipsilateral L3/4 dorsal horn 15 weeks after initiation of ND or HFD feeding. *n* = 3 mice/group.***P* < 0.01, 2-way ANOVA followed by Tukey's post hoc test.

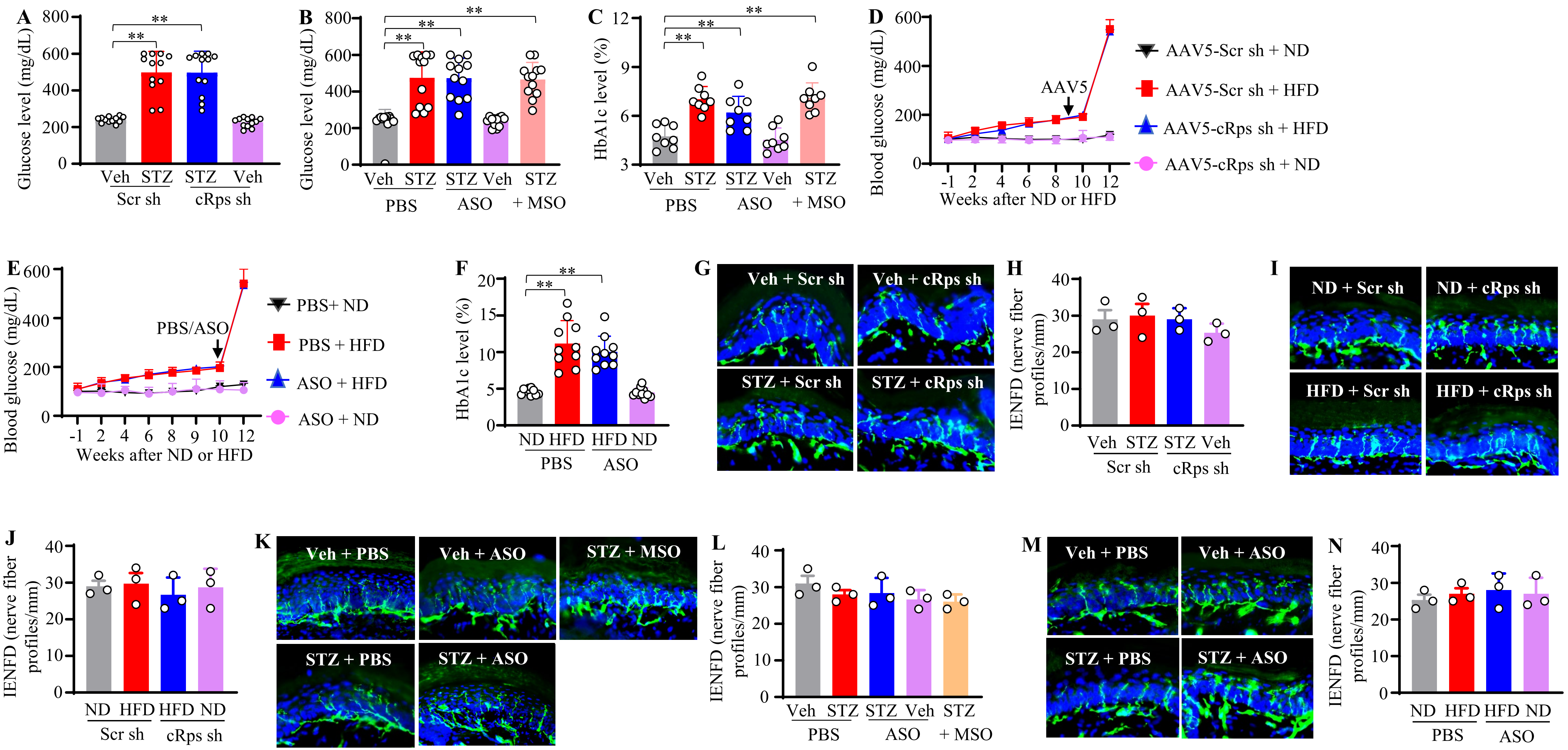
A
Glucose level (mg/dL)
Veh STZ STZ Veh
Scr sh cRps sh
B
Glucose level (mg/dL)
Veh STZ STZ Veh STZ
PBS ASO + MSO
C
HbA1c level (%)
Veh STZ STZ Veh STZ
PBS ASO + MSO
D
Blood glucose (mg/dL)
AAV5
Weeks after ND or HFD
AAV5-Scr sh + ND
AAV5-Scr sh + HFD
AAV5-cRps sh + HFD
AAV5-cRps sh + ND
E
Blood glucose (mg/dL)
PBS/ASO
Weeks after ND or HFD
PBS+ ND
PBS + HFD
ASO + HFD
ASO + ND
F
HbA1c level (%)
ND HFD HFD ND
PBS ASO
G
Veh + Scr sh
Veh + cRps sh
STZ + Scr sh
STZ + cRps sh
H
IENFD (nerve fiber profiles/mm)
Veh STZ STZ Veh
Scr sh cRps sh
I
ND + Scr sh
ND + cRps sh
HFD + Scr sh
HFD + cRps sh
J
IENFD (nerve fiber profiles/mm)
ND HFD HFD ND
Scr sh cRps sh
K
Veh + PBS
Veh + ASO
STZ + MSO
STZ + PBS
STZ + ASO
L
IENFD (nerve fiber profiles/mm)
Veh STZ STZ Veh STZ
PBS ASO + MSO
M
Veh + PBS
Veh + ASO
STZ + PBS
STZ + ASO
N
IENFD (nerve fiber profiles/mm)
ND HFD HFD ND
PBS ASO

**Supplementary Fig. 9**. (A) Blood glucose levels 35 days after unilateral microinjection of AAV5-cRps sh or AAV5-Scr sh into the L3/4 DRGs of male mice receiving STZ or vehicle injection 3 days before viral microinjection. n = 12 mice/group. $^{**}P < 0.01$, 2-way ANOVA followed by Tukey's post hoc test. (B, C) Levels of blood glucose (B) and HbA1c (C) 49 days after STZ or vehicle injection in male mice receiving intrathecal administration of PBS, ASO (100 µg), or MSO (100 µg) 2 weeks post-STZ or vehicle injection. n = 12 mice/group. $^{**}P < 0.01$, 2-way ANOVA followed by Tukey's post hoc test. (D) Blood glucose levels at the different weeks as indicated after initiation of ND or HFD feeding in male mice receiving unilateral microinjection of AAV5-cRps sh or AAV5-Scr sh into the L3/4 DRGs 9 weeks after initiation of ND or HFD feeding. n = 12 mice/group. three-way ANOVA followed by Tukey's post hoc test. (E, F) Level of blood glucose at the different weeks as indicated after initiation of ND or HFD feeding (E) and level of HbA1c 12 weeks after initiation of ND or HFD feeding (F) in male mice receiving intrathecal administration of PBS or ASO (100 µg) 10 weeks after initiation of ND or HFD feeding. n = 10 mice/group. $^{**}P < 0.01$, 2-way ANOVA followed by Tukey's post hoc test. (G, H) Representative photomicrographs (G) and quantification of intraepidermal nerve fiber density (IENFD; H) 35 days after unilateral microinjection of AAV5-cRps sh or AAV5-Scr sh into the L3/4 DRGs of male mice. They received STZ or vehicle injection 3 days before viral microinjection. IENFD in paw plantar side was stained with PGP 9.5. n = 3 mice/group. 2-way ANOVA followed by Tukey's post hoc test. (I, J) Representative photomicrographs (I) and quantification of intraepidermal nerve fiber density (IENFD; J) 15 weeks after initiation of ND or HFD feeding in male mice. They received unilateral microinjection of AAV5-cRps sh or AAV5-Scr sh into the L3/4 DRGs 9 weeks after initiation of ND or HFD feeding. n = 3 mice/group. 2-way ANOVA followed by Tukey's post hoc test. (K, L) Representative photomicrographs (K) and quantification of intraepidermal nerve fiber density (IENFD; L) 49 days days after STZ or vehicle injection in male mice receiving intrathecal administration of PBS, ASO (100 µg), or MSO (100 µg) 2 weeks post-STZ or vehicle injection. n = 3 mice/group. 2-way ANOVA followed by Tukey's post hoc test. (M, N) Representative photomicrographs (M) and quantification of intraepidermal nerve fiber density (IENFD; N) 15 weeks after initiation of ND or HFD feeding in male mice receiving intrathecal administration of PBS or ASO (100 µg) 10 weeks after initiation of ND or HFD feeding. n = 3 mice/group. 2-way ANOVA followed by Tukey's post hoc test.

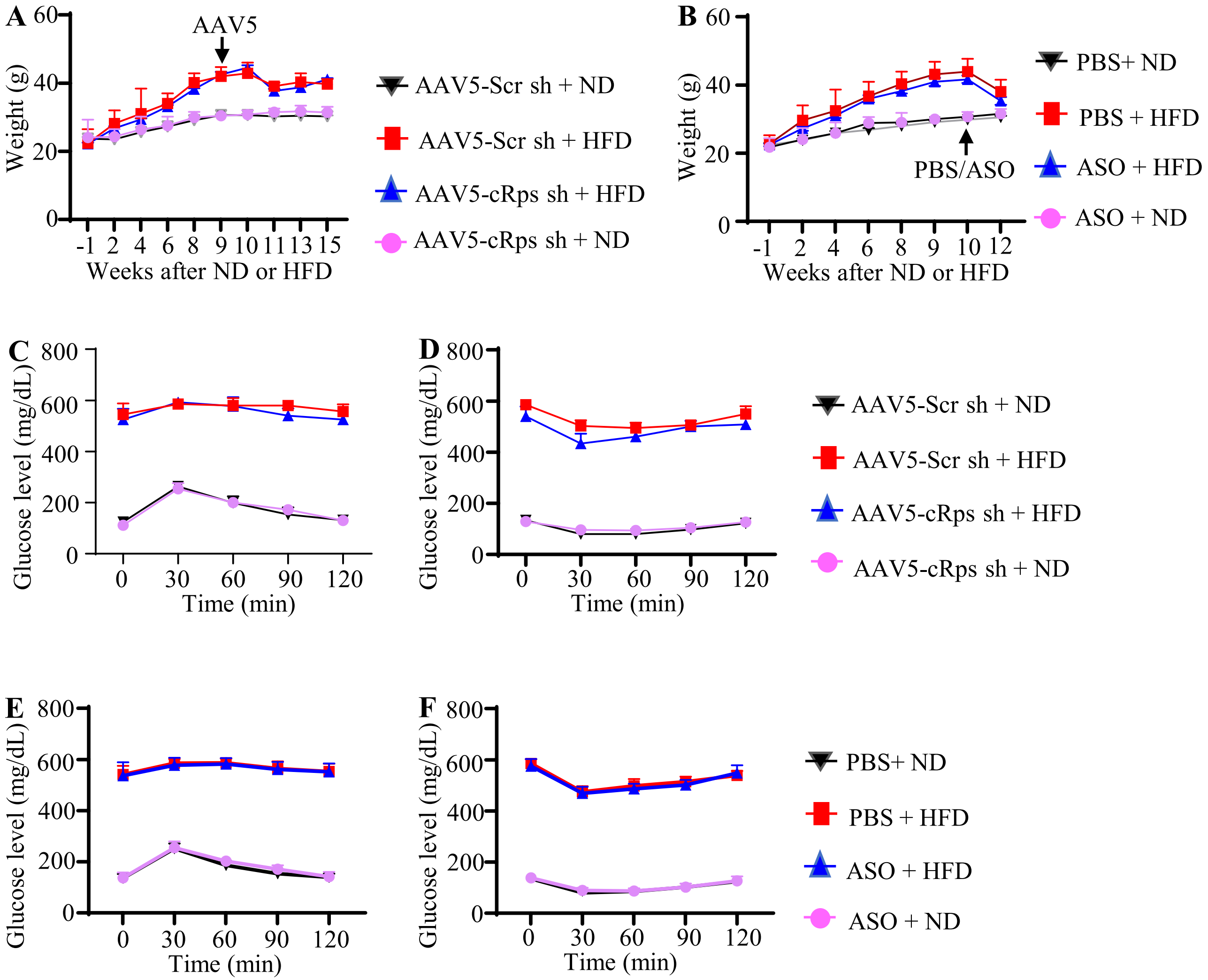

**Supplementary Fig. 10**. (A) Body weights at the different weeks after initiation of ND or HFD feeding in male mice receiving unilateral microinjection of AAV5-cRps sh or AAV5-Scr sh 9 weeks after initiation of ND or HFD feeding. n = 12 mice/group. Three-way ANOVA followed by Tukey's post hoc test. (B) Body weights at the different weeks after initiation of ND or HFD feeding in male mice receiving intrathecal administration of PBS or ASO (100 µg) 10 weeks after initiation of ND or HFD feeding. n = 10 mice/group. n = 10 mice/group. Three-way ANOVA followed by Tukey's post hoc test. (C, D) Glucose tolerance test (C) and insulin tolerance test (D) were performed 12 weeks after initiation of ND or HFD feeding (that is, 2 weeks after STZ or vehicle injection) in male mice receiving unilateral microinjection of AAV5-cRps sh or AAV5-Scr sh 9 weeks after initiation of ND or HFD feeding. n = 7-12 mice/group. Three-way ANOVA followed by Tukey's post hoc test. (E, F) Glucose tolerance test (E) and insulin tolerance test (F) were performed 12 weeks after initiation of ND or HFD feeding (that is, 2 weeks after STZ or vehicle injection) in male mice receiving intrathecal administration of PBS or ASO (100 µg) 10 weeks after initiation of ND or HFD feeding. n = 12 mice/group. Three-way ANOVA followed by Tukey's post hoc test. .

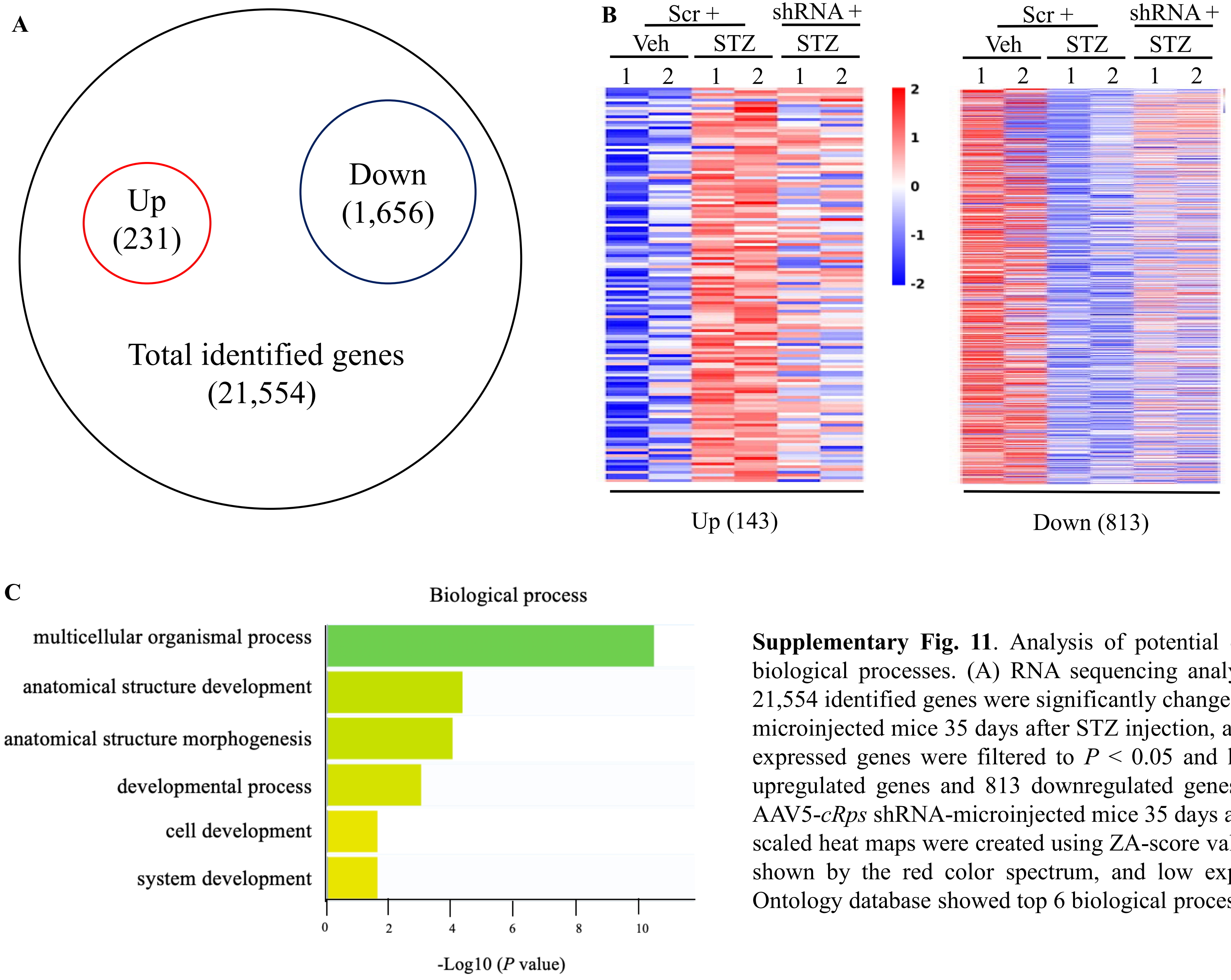

**Supplementary Fig. 11**. Analysis of potential downstream targets of *cRps* and their involvement in biological processes. (A) RNA sequencing analysis revealed that about 1,887 genes out of a total of 21,554 identified genes were significantly changed in the ipsilateral L3/4 DRG from the AAV5-Scr RNA-microinjected mice 35 days after STZ injection, as compared to that after vehicle injection. Differentially expressed genes were filtered to $P < 0.05$ and $\log_2$fold-change. (B) Heatmaps showed that about 143 upregulated genes and 813 downregulated genes were reversed in the ipsilateral L3/4 DRG from the AAV5-*cRps* shRNA-microinjected mice 35 days after STZ injection. n = 2 replicates (1 and 2)/group. The scaled heat maps were created using ZA-score values obtained from RNA sequencing. High expression is shown by the red color spectrum, and low expression is shown in blue. (C) Analysis of the Gene Ontology database showed top 6 biological process functions of these differentially expressed genes.

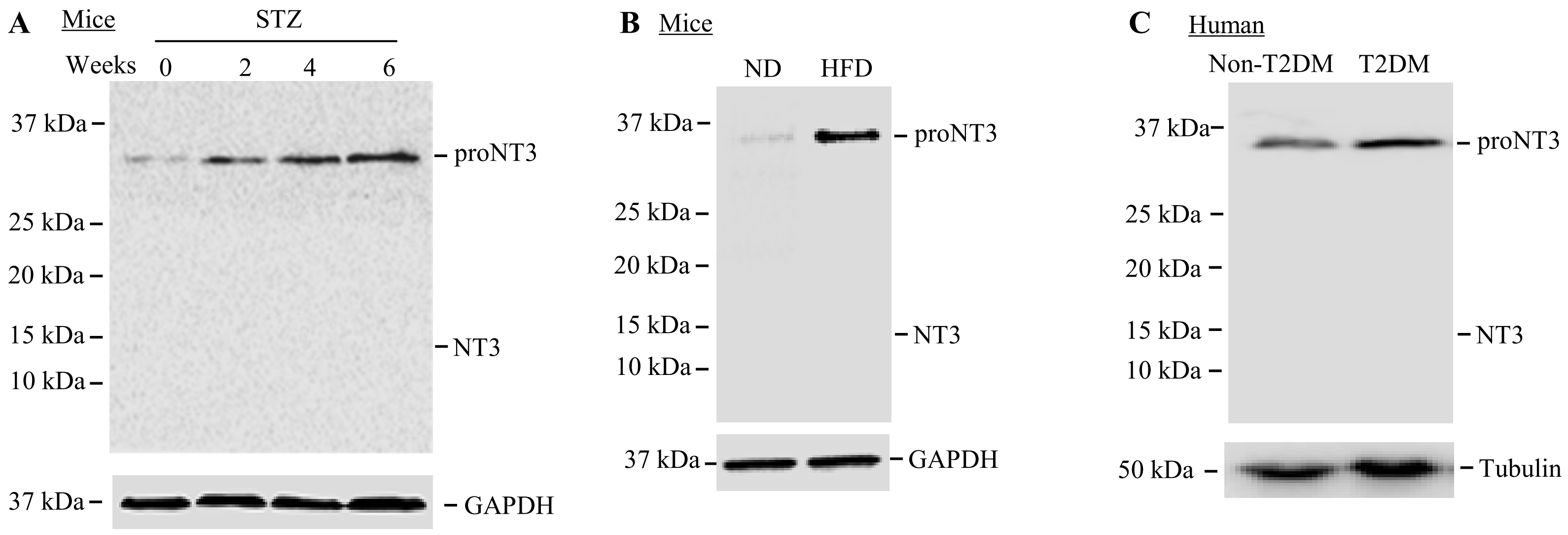

**Supplementary Fig. 12**. proNT3 (~32 kDa) , but not mature NT3 (~14.5 kDa), is detected in the DRG from mice and human donors. The anti-NT3 antibody (ANT-003, Alomone Labs) used has been demonstrated to detect both proNT3 and mature NT3 in previous studies[11,12]. (A) proNT3 expression is time-dependently upregulated in the bilateral DRGs following intraperitoneal injection of streptozotocin (STZ). Mature NT3 is not detected in the bilateral L3/4 DRG at 0, 2, 4, or 6 weeks post-STZ. (B) proNT3 expression is upregulated in the bilateral DRGs 15 weeks after high-fat diet (HFD) feeding. Mature NT3 is not detected in the bilateral DRGs 15 weeks after normal diet (ND)- or HFD-feeding. (C) proNT3 expression is upregulated in the DRG from human donors with T2DM and pain symptoms. Mature NT3 is not detected in the bilateral DRGs from human donors with or without T2DM.

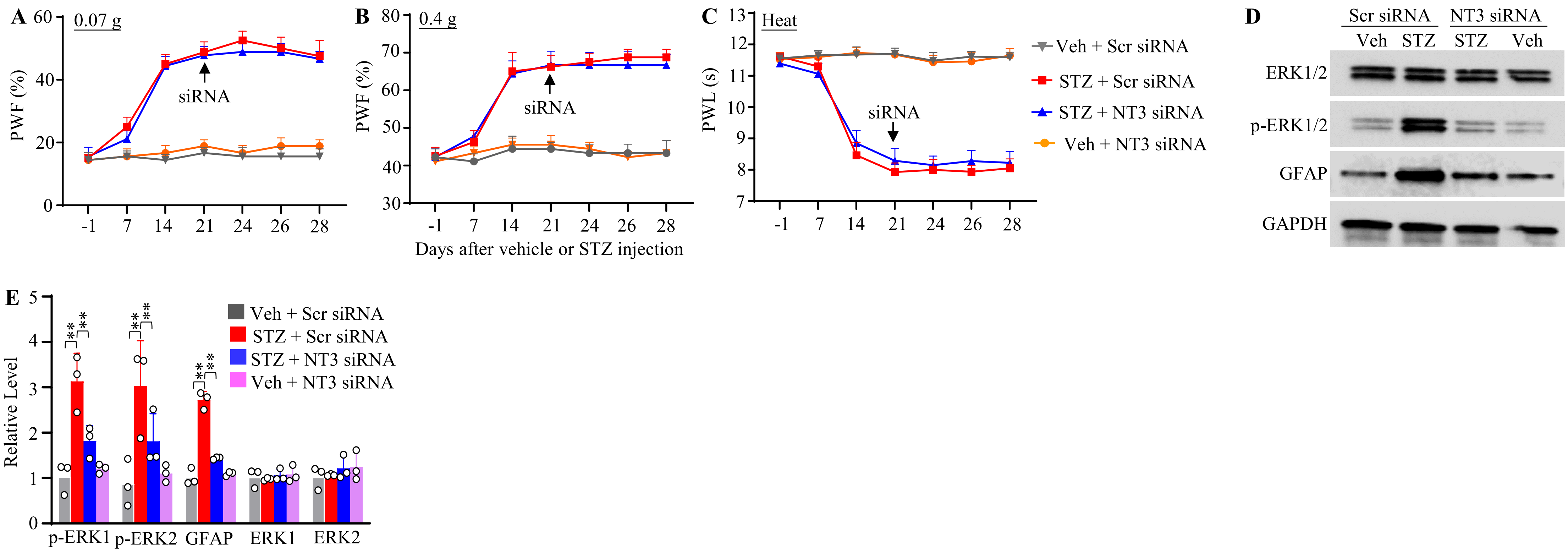

**Supplementary Fig. 13** (A-C) Effect of unilateral L3/L4 DRG microinjection of NT3 siRNA or Scr siRNA 21 days after STZ or vehicle injection on paw withdrawal frequency (PWF) to 0.07-g (A) and 0.4-g (B) von Frey filament stimuli and paw withdrawal latency (PWL) to heat (C) stimulation at the indicated days after STZ or vehicle injection on the contralateral side in male mice. *n* = 8 mice/group. Three-way repeated-measures ANOVA followed by Tukey's post hoc test. (D. E) Effect of unilateral L3/L4 DRG microinjection of NT3 siRNA or Scr siRNA 21 days after STZ or vehicle injection on levels of p-ERK1/2, ERK1/2, and GFAP in the ipsilateral L3/4 dorsal horn 28 days after STZ or vehicle injection in male mice. *n* = 3 mice/group. $^{**}P < 0.01$, 2-way ANOVA followed by Tukey's post hoc test.

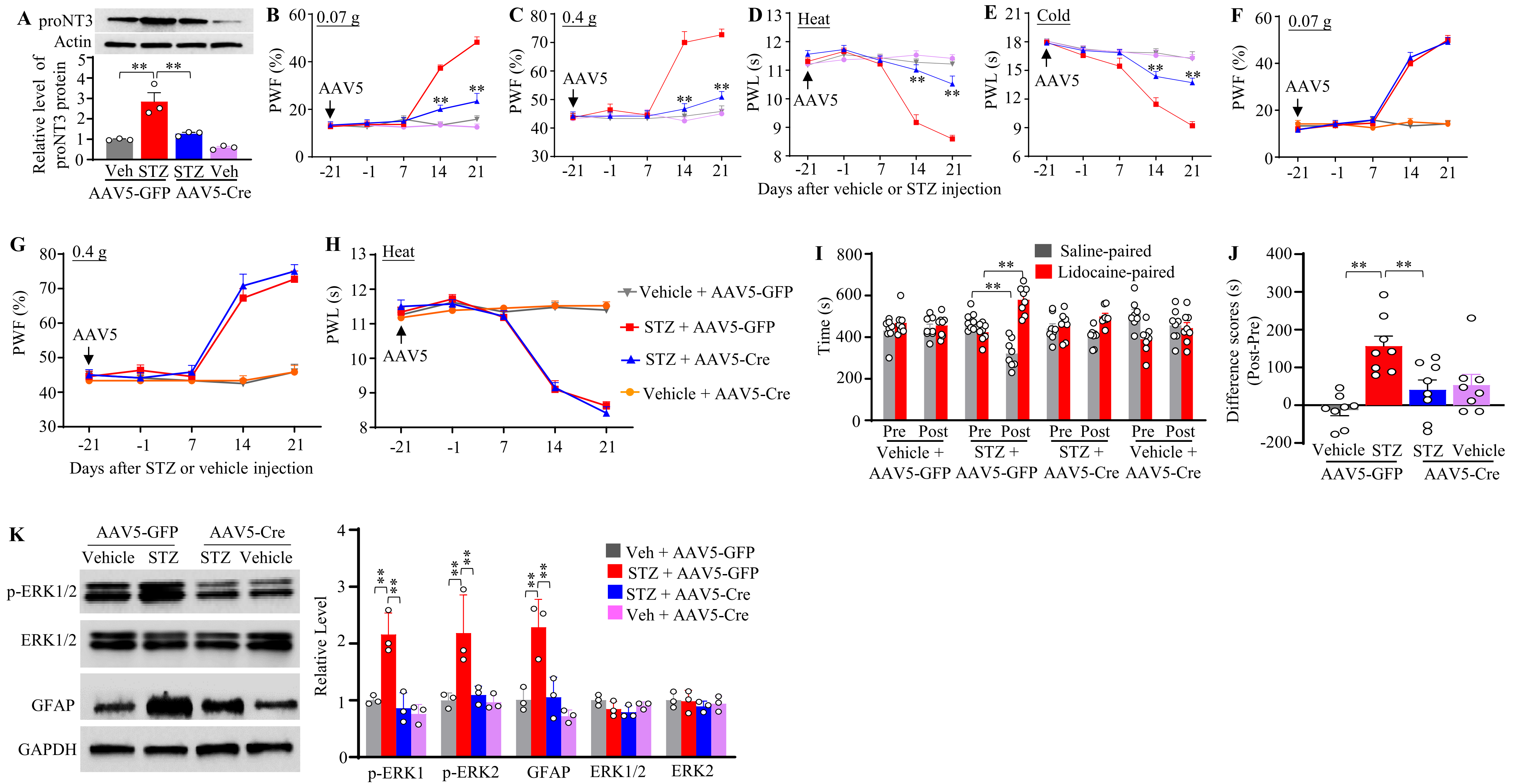
A
proNT3
Actin
Relative level of proNT3 protein
Veh STZ STZ Veh
AAV5-GFP AAV5-Cre
B 0.07 g
PWF (%)
AAV5
C 0.4 g
PWF (%)
AAV5
D Heat
PWL (s)
AAV5
Days after vehicle or STZ injection
E Cold
PWL (s)
AAV5
F 0.07 g
PWF (%)
AAV5
G 0.4 g
PWF (%)
AAV5
Days after STZ or vehicle injection
H Heat
PWL (s)
AAV5
Vehicle + AAV5-GFP
STZ + AAV5-GFP
STZ + AAV5-Cre
Vehicle + AAV5-Cre
I
Saline-paired
Lidocaine-paired
Time (s)
Pre Post
Vehicle + AAV5-GFP
STZ + AAV5-GFP
STZ + AAV5-Cre
Vehicle + AAV5-Cre
J
Difference scores (s) (Post-Pre)
Vehicle STZ STZ Vehicle
AAV5-GFP AAV5-Cre
K
AAV5-GFP AAV5-Cre
Vehicle STZ STZ Vehicle
p-ERK1/2
ERK1/2
GFAP
GAPDH
Relative Level
Veh + AAV5-GFP
STZ + AAV5-GFP
STZ + AAV5-Cre
Veh + AAV5-Cre
p-ERK1 p-ERK2 GFAP ERK1/2 ERK2

**Supplementary Fig. 14**. (A) Expression level of proNT3 protein in the ipsilateral L3/L4 DRGs 21 days after STZ or vehicle injection in male *Nt3*$^{f/f}$ mice receiving pre-microinjection of AAV5-Cre or AAV5-GFP into the unilateral L3/L4 DRGs 21 days before STZ or vehicle injection. $n = 3$ replicates (2 mice/replicate)/group. $^{**}P < 0.01$, two-way ANOVA followed by Tukey's post hoc test. (B–H) Effects of microinjection of AAV5-Cre or AAV5-GFP 21 days before STZ or vehicle injection on PWF to 0.07-g (B and F) and 0.4-g (C and G) von Frey filament stimuli and PWL to heat (D and H) and cold (E) stimuli at the indicated days after STZ or vehicle injection on the ipsilateral (B-E) and contralateral (F-H) sides in male *Nt3*$^{f/f}$ mice. $n = 12$ mice/group. $^{**}P < 0.01$ versus the STZ plus AAV5-GFP group at the corresponding time points; three-way repeated-measures ANOVA followed by Tukey's post hoc test. (I and J) Effects of microinjection of AAV5-Cre or AAV5-GFP 21 days before STZ or vehicle injection on spontaneous ongoing pain, assessed using the CPP paradigm 14 days post-STZ or vehicle injection in male *Nt3*$^{f/f}$ mice. Pre, preconditioning; Post, post-conditioning. $n = 8$ mice/group. $^{**}P < 0.01$, two- (J) or three- (I) way ANOVA followed by Tukey's post hoc test. (K) Effects of microinjection of AAV5-Cre or AAV5-GFP 21 days before STZ or vehicle injection on levels of p-ERK1/2, ERK1/2, and GFAP in the ipsilateral L3/4 dorsal horn of male *Nt3*$^{f/f}$ mice 21 days after STZ or vehicle injection. $n = 3$ mice/group. $^{**}P < 0.01$, 2-way ANOVA followed by Tukey's post hoc test.

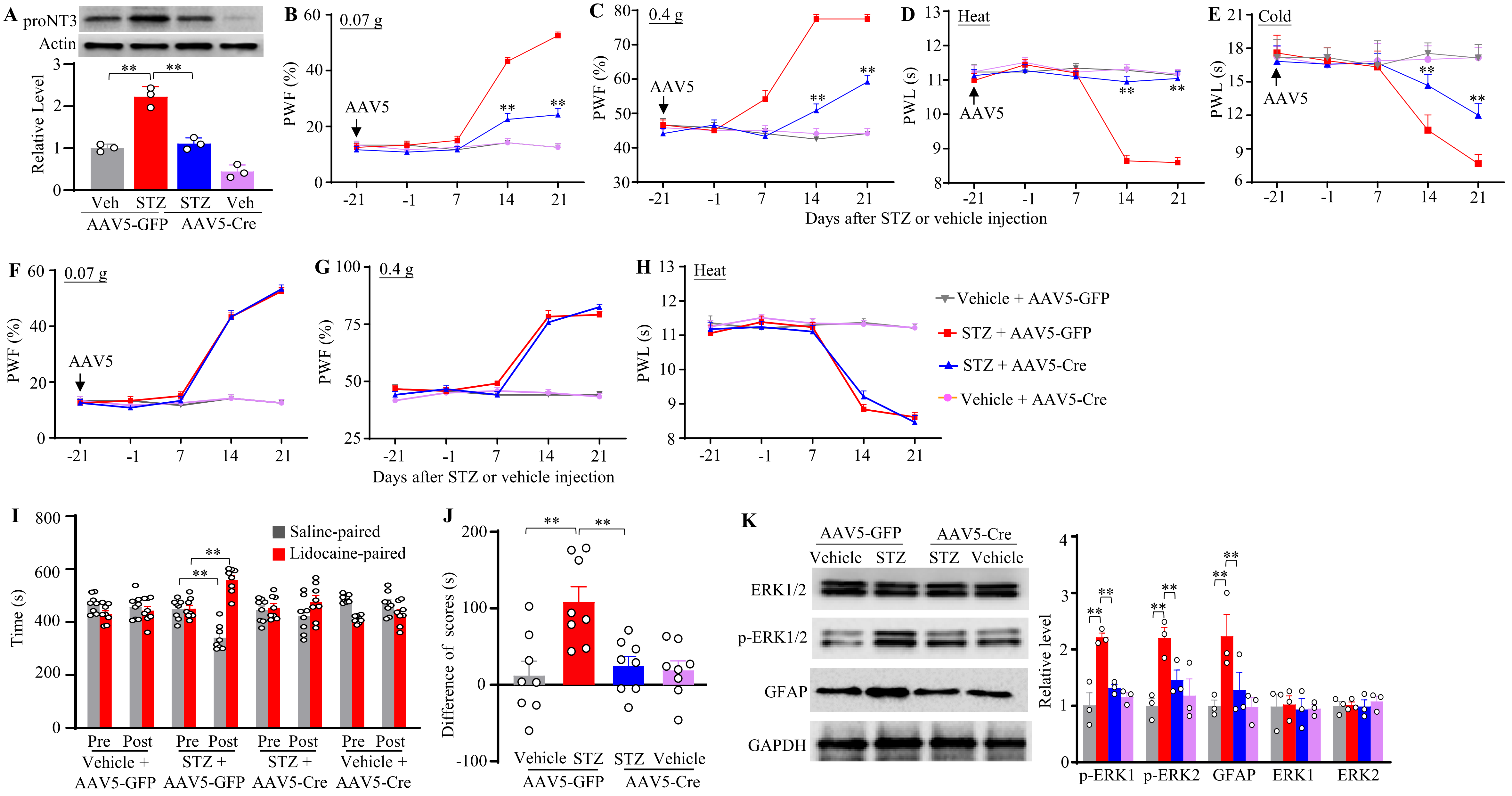
A
proNT3
Actin
Relative Level
Veh
STZ
STZ
Veh
AAV5-GFP
AAV5-Cre
B
0.07 g
PWF (%)
AAV5
C
0.4 g
D
Heat
PWL (s)
E
Cold
Days after STZ or vehicle injection
F
G
H
Vehicle + AAV5-GFP
STZ + AAV5-GFP
STZ + AAV5-Cre
Vehicle + AAV5-Cre
I
Time (s)
Saline-paired
Lidocaine-paired
Pre
Post
Vehicle + AAV5-GFP
STZ + AAV5-GFP
STZ + AAV5-Cre
Vehicle + AAV5-Cre
J
Difference of scores (s)
Vehicle
STZ
K
ERK1/2
p-ERK1/2
GFAP
GAPDH
Relative level
p-ERK1
p-ERK2
ERK1
ERK2

**Supplementary Fig. 15**. Blocking increases of *Nt3* mRNA and proNT3 protein in the DRG attenuates STZ-induced diabetic neuropathic pain in female *Nt3*$^{f/f}$ mice. (A) Expression level of proNT3 protein in the ipsilateral L3/L4 DRGs 21 days after STZ or vehicle injection in female *Nt3*$^{f/f}$ mice receiving pre-microinjection of AAV5-Cre o AAV5-GFP into the unilateral L3/L4 DRGs 21 days before STZ or vehicle injection. *n* = 3 replicates (2 mice/replicate)/group. ***P* < 0.01, two-way ANOVA followed by Tukey's post hoc test. (B-H) Effects of microinjection of AAV5-Cre or AAV5-GFP 21 days before STZ or vehicle injection on paw withdrawal frequency (PWF) to 0.07-g (B and F) and 0.4-g (C and G) von Frey filament stimuli and paw withdrawal latency (PWL) to heat (D and H) and cold (E) stimuli at the indicated days after STZ or vehicle injection on the ipsilateral (B-E) and contralateral (F-H) sides in female *Nt3*$^{f/f}$ mice. *n* = 12 mice/group. ***P* < 0.01 versus the STZ plus AAV5-GFP group at the corresponding time points; three-way repeated-measures ANOVA followed by Tukey's post hoc test. (I, J) Effects of microinjection of AAV5-Cre or AAV5-GFP 21 days before STZ or vehicle injection on spontaneous ongoing pain, assessed using the CPP paradigm 14 days post-STZ or vehicle injection in female *Nt3*$^{f/f}$ mice. Pre, preconditioning; Post, post-conditioning. *n* = 8 mice/group. ***P* < 0.01, two- (I) or three- (J) way ANOVA followed by Tukey's post hoc test. (K) Effects of microinjection of AAV5-Cre or AAV5-GFP 21 days before STZ or vehicle injection on the levels of p-ERK1/2, ERK1/2, and GFAP in the ipsilateral L3/4 dorsal horn of female *Nt3*$^{f/f}$ mice 21 days after STZ or vehicle injection. *n* = 3 mice/group. ***P* < 0.01, 2-way ANOVA followed by Tukey's post hoc test.

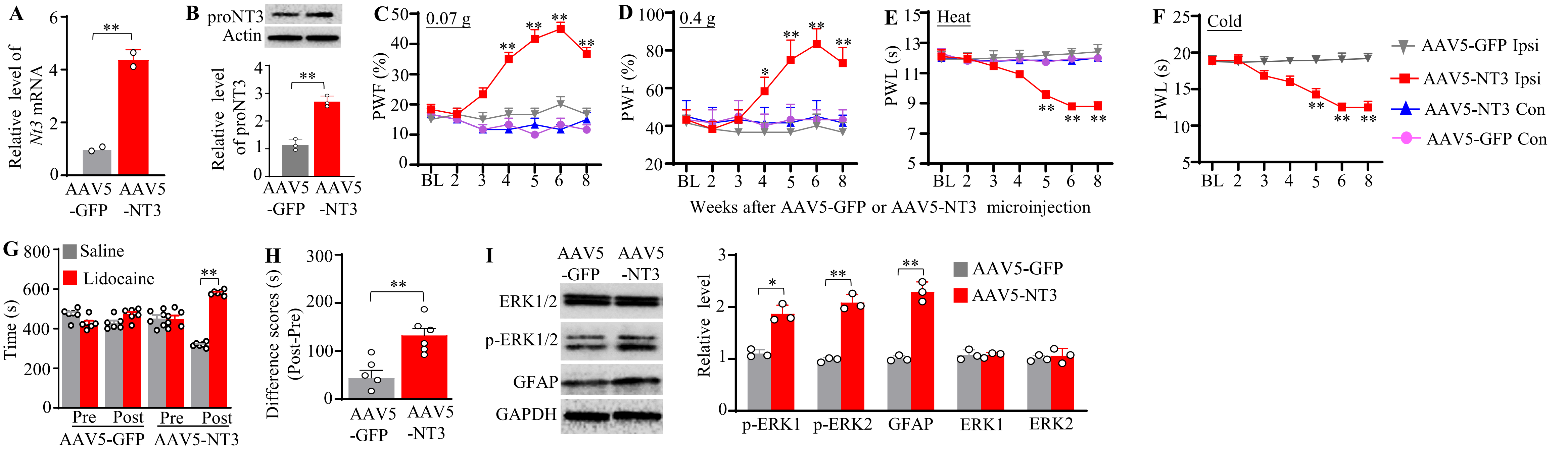

**Supplementary Fig. 16**. (A, B) Expression levels of *Nt3* mRNA (A) and proNT3 protein (B) in the ipsilateral L3/L4 DRGs of female mice 8 weeks after unilateral microinjection of AAV5-NT3 or AAV5-GFP into L3/4 DRGs. n = 3 replicates (2 mice/replicate)/group. $^{**}P < 0.01$, two-tailed unpaired Student's t test. (C–F) Effects of unilateral L3/L4 DRG microinjection of AAV5-NT3 or AAV5-GFP on paw withdrawal frequency (PWF) to 0.07-g (C) and 0.4-g (D) von Frey filament stimuli and paw withdrawal latency (PWL) to heat (E) and cold (F) stimuli at the indicated weeks after viral microinjection on the ipsilateral (Ipsi) and contralateral (Con) sides in female mice. $n = 6$ mice/group. $^{*}P < 0.05$, $^{**}P < 0.01$ versus the AAV5-GFP group at the corresponding time points; two-way repeated-measures ANOVA followed by Tukey's post hoc test. (G, H) Effects of unilateral L3/L4 DRG microinjection of AAV5-NT3 or AAV5-GFP on spontaneous ongoing pain, assessed using the CPP paradigm 7 weeks after viral microinjection in female mice. Pre, preconditioning; Post, post-conditioning. $n = 6$ mice/group. $^{**}P < 0.01$, two-way repeated-measures ANOVA followed by Tukey's post hoc test (G) or two-tailed unpaired Student's *t*-test (H). (I) Effects of unilateral L3/L4 DRG microinjection of AAV5-NT3 or AAV5-GFP on levels of p-ERK1/2, ERK1/2, and GFAP in the ipsilateral L3/4 dorsal horn 8 weeks after microinjection in female mice. $n = 3$ mice/group. $^{**}P < 0.01$, two-tailed unpaired Student's *t* test.

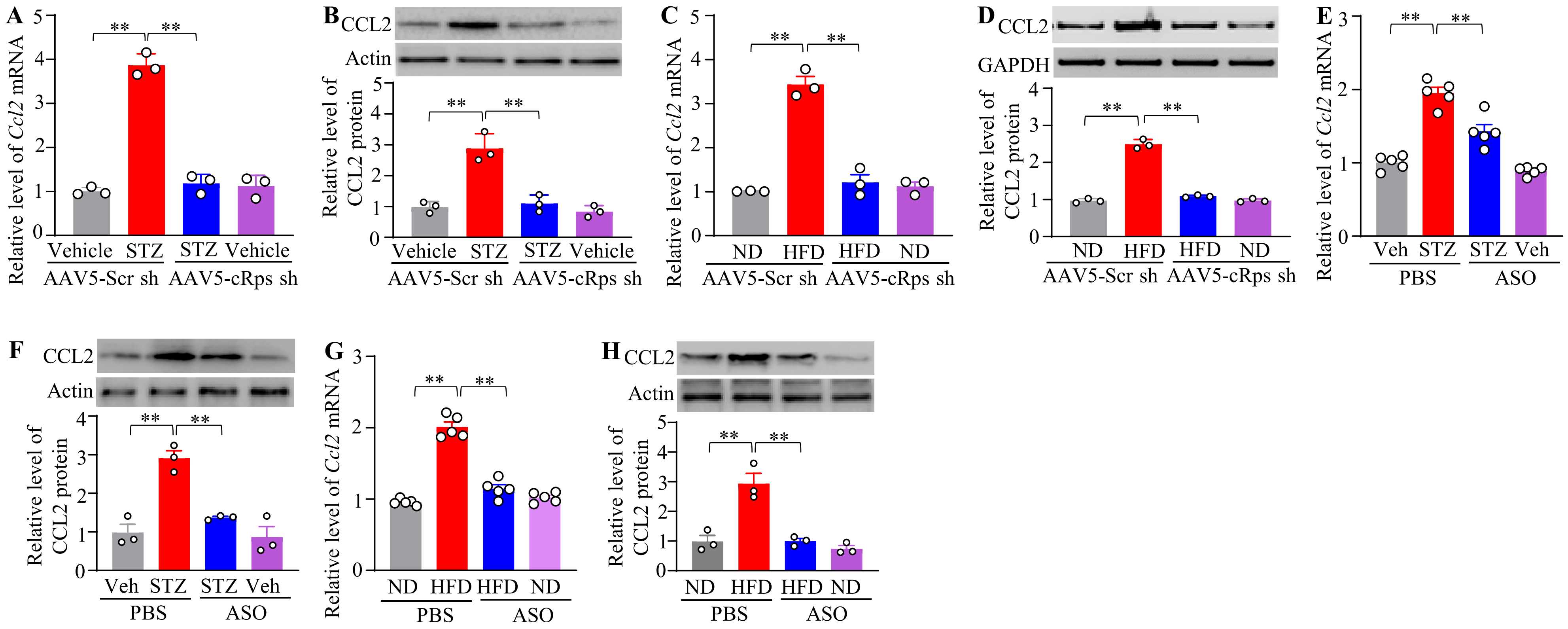

**Supplementary Fig. 17.** Blocking DRG *cRps* upregulation attenuates increases in *Ccl2* mRNA and CCL2 protein levels in DRGs under T1DM and T2DM conditions. (A, B) Expression levels of *Ccl2* mRNA (A) and CCL2 protein (B) in the ipsilateral L3/4 DRGs 35 days after unilateral microinjection of AAV5-cRps sh or AAV5-Scr sh into the L3/4 DRGs in male mice receiving STZ or vehicle injection 3 days before viral microinjection. n = 3 replicates (6 mice)/group/assay. ***P* < 0.01, 2-way ANOVA followed by Tukey's post hoc test. (C, D) Expression levels of *Ccl2* mRNA (C) and CCL2 protein (D) in the ipsilateral L3/4 DRGs 15 weeks after initiation of ND or HFD feeding in male mice receiving unilateral microinjection of AAV5-cRps sh or AAV5-Scr sh 9 weeks after initiation of ND or HFD feeding. n = 3 replicates (6 mice)/group/assay. ***P* < 0.01, 2-way ANOVA followed by Tukey's post hoc test. (E, F) Expression levels of *Ccl2* mRNA (E) and CCL2 protein (F) in the bilateral L3/4 DRGs 49 days after STZ or vehicle injection in male mice receiving intrathecal administration of PBS or ASO (100 µg) 2 weeks post-STZ or vehicle injection. n = 3 mice/group/assay. ***P* < 0.01, 2-way ANOVA followed by Tukey's post hoc test. (G, H) Expression levels of *Ccl2* mRNA (G) and CCL2 protein (H) in the bilateral L3/4 DRGs 15 weeks after initiation of ND or HFD feeding in male mice receiving intrathecal administration of PBS or ASO (100 µg) 10 weeks after initiation of ND or HFD feeding. n = 3 mice/group/assay. ***P* < 0.01, 2-way ANOVA followed by Tukey's post hoc test.

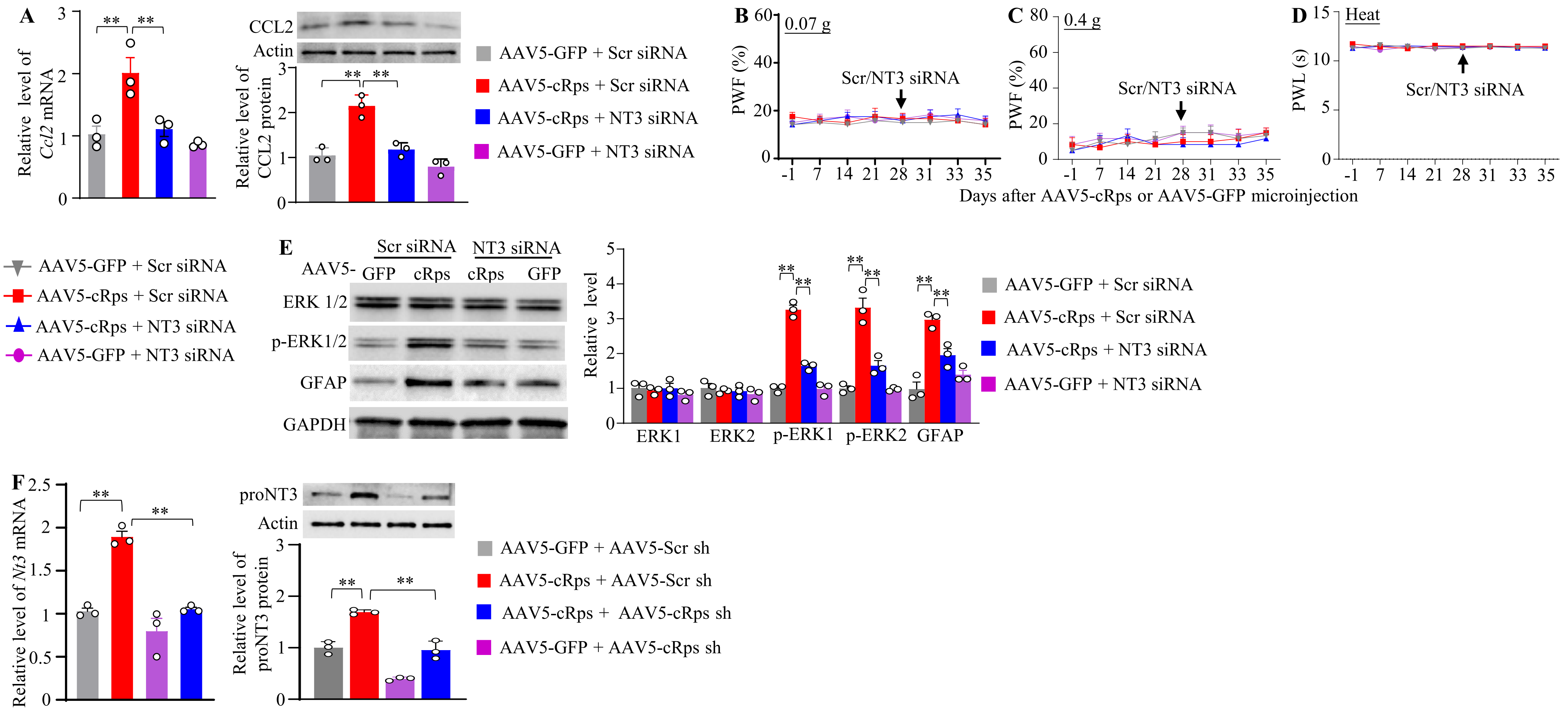
A
Relative level of Ccl2 mRNA
CCL2
Actin
Relative level of CCL2 protein
AAV5-GFP + Scr siRNA
AAV5-cRps + Scr siRNA
AAV5-cRps + NT3 siRNA
AAV5-GFP + NT3 siRNA
B
0.07 g
PWF (%)
Scr/NT3 siRNA
C
0.4 g
D
Heat
PWL (s)
Days after AAV5-cRps or AAV5-GFP microinjection
E
Scr siRNA
NT3 siRNA
AAV5- GFP cRps cRps GFP
ERK 1/2
p-ERK1/2
GFAP
GAPDH
Relative level
ERK1
ERK2
p-ERK1
p-ERK2
F
Relative level of Nt3 mRNA
proNT3
Relative level of proNT3 protein
AAV5-GFP + AAV5-Scr sh
AAV5-cRps + AAV5-Scr sh
AAV5-cRps + AAV5-cRps sh
AAV5-GFP + AAV5-cRps sh

**Supplementary Fig. 18.** (A) Expression levels of *Ccl2* mRNA (A) and CCL2 protein (B) in the ipsilateral L3/4 DRGs 35 days after unilateral microinjection of AAV5-cRps or AAV5-GFP into the L3/4 DRGs in male mice receiving microinjection of *Nt3* siRNA or control scrambled (Scr) siRNA into the ipsilateral L3/4 DRGs 28 days post viral microinjection. n = 3 replicates (6 mice)/group/assay. $^{**}P < 0.01$, one-way ANOVA followed by Tukey's post hoc test. (B-D) Effect of microinjection of *Nt3* siRNA or control scrambled (Scr) siRNA into the ipsilateral L3/4 DRGs 28 days post viral microinjection on paw withdrawal frequency (PWF) to 0.07-g (B) and 0.4-g (C) von Frey filament stimuli and paw withdrawal latency (PWL) to heat stimulation (D) on the days as indicated post-viral microinjection on the contralateral side in male mice. They received unilateral microinjection of AAV5-cRps or AAV5-GFP into the L3/4 DRGs. n = 12 mice/group. Two-way repeated-measures ANOVA followed by Tukey's post hoc test. (E) Effect of microinjection of *Nt3* siRNA or control scrambled (Scr) siRNA into the ipsilateral L3/4 DRGs 28 days post viral microinjection on levels of p-ERK1/2, total ERK1/2 and GFAP in the ipsilateral L3/4 dorsal horn 35 days after viral microinjection. They received unilateral microinjection of AAV5-cRps or AAV5-GFP into the L3/4 DRGs. n = 3 mice/group. One-way repeated-measures ANOVA followed by Tukey's post hoc test. (F) Expression levels of *Nt3* mRNA and proNT3 protein in the ipsilateral L3/4 DRGs 35 days after unilateral co-microinjection of AAV5-cRps plus AAV5-cRps shRNA or AAV5-Scr shRNA as well as of AAV5-GFP plus AAV5-cRps shRNA or AAV5-Scr shRNA in the L3/4 DRGs. n = 3 repeats (6 mice)/group/assay. $^{**}P < 0.01$, one-way ANOVA followed by Tukey's post hoc test.

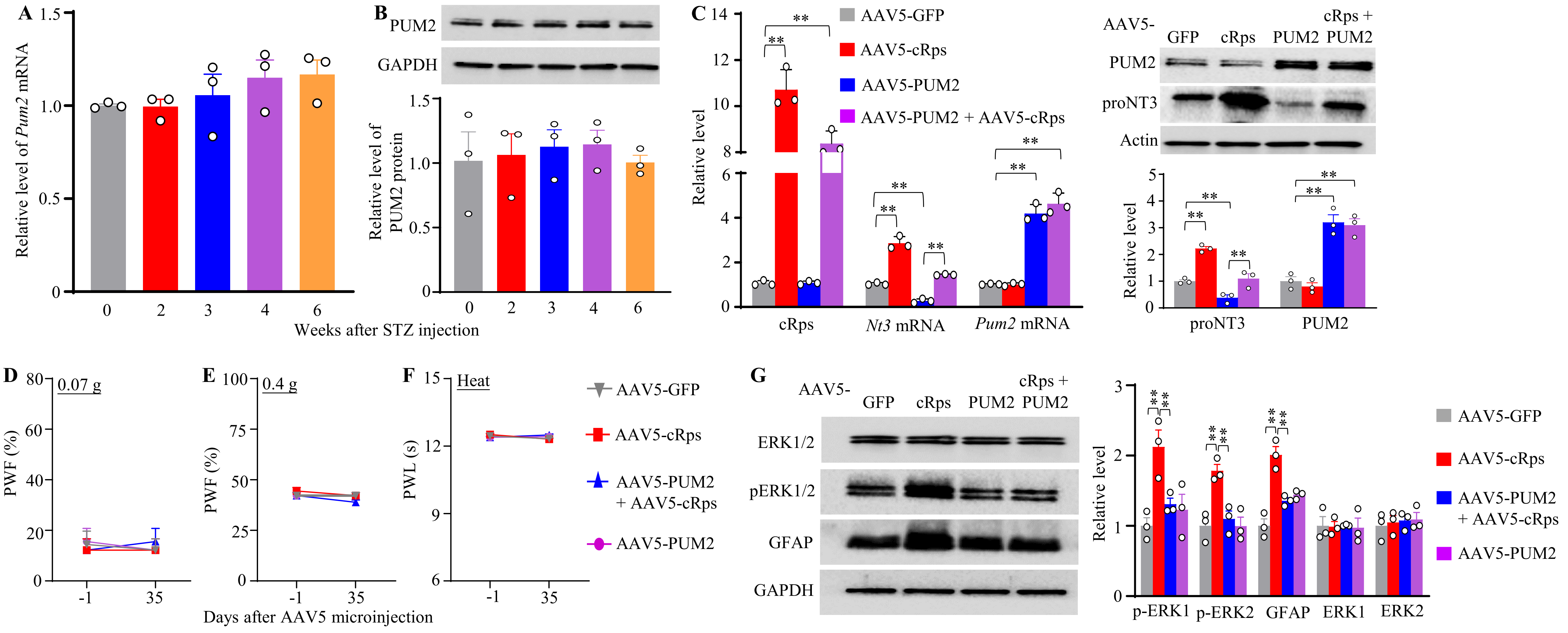

A
Relative level of Pum2 mRNA
Weeks after STZ injection
B
PUM2
GAPDH
Relative level of PUM2 protein
C
AAV5-GFP
AAV5-cRps
AAV5-PUM2
AAV5-PUM2 + AAV5-cRps
Relative level
cRps
Nt3 mRNA
Pum2 mRNA
AAV5-
GFP
cRps
PUM2
cRps + PUM2
PUM2
proNT3
Actin
proNT3
PUM2
D
0.07 g
PWF (%)
E
0.4 g
PWF (%)
F
Heat
PWL (s)
Days after AAV5 microinjection
AAV5-GFP
AAV5-cRps
AAV5-PUM2 + AAV5-cRps
AAV5-PUM2
G
AAV5-
GFP
cRps
PUM2
cRps + PUM2
ERK1/2
pERK1/2
GFAP
GAPDH
Relative level
p-ERK1
p-ERK2
GFAP
ERK1
ERK2
AAV5-GFP
AAV5-cRps
AAV5-PUM2 + AAV5-cRps
AAV5-PUM2

**Supplementary Fig. 19.** (A, B) Expression levels of *Pum2* mRNA (A) and PUM2 protein (B) in lumbar DRGs of male mice 0, 2, 3, 4 and 6 weeks after STZ injection. n = 3 replicates (6 mice)/time point/assay. One-way ANOVA followed by Tukey's post hoc test. (C) Expression levels of *cRps, Nt3* mRNA, *Pum2* mRNA, PUM2 and proNT3 in the ipsilateral L3/4 DRGs 35 days after unilateral microinjection of AAV5-GFP, AAV5-cRps, AAV5-PUM2 and AAV5-PUM2 plus AAV5-cRps, respectively, into the L3/4 DRGs. n = 3 replicates (6 mice)/time point/assay. ***P* < 0.01, one-way ANOVA followed by Tukey's post hoc test.. (D-F) Effect of after unilateral microinjection of AAV5-GFP, AAV5-cRps, AAV5-PUM2 and AAV5-PUM2 plus AAV5-cRps, respectively, into the L3/4 DRGs on paw withdrawal frequency (PWF) to 0.07-g (D) and 0.4-g (E) von Frey filament stimuli and paw withdrawal latency (PWL) to heat stimulation (F) on days as indicated, measured on the contralateral side. n = 11 mice/group. Two-way repeated-measures ANOVA followed by Tukey's post hoc test. (G) Effect of after unilateral microinjection of AAV5-GFP, AAV5-cRps, AAV5-PUM2 and AAV5-PUM2 plus AAV5-cRps, respectively, into the L3/4 DRGs on levels of p-ERK1/2, total ERK1/2 and GFAP in the ipsilateral L3/4 spinal cord dorsal horn 35 days after viral microinjection. n = 3 mice/group. ***P* < 0.01, one-way ANOVA followed by Tukey's post hoc test.

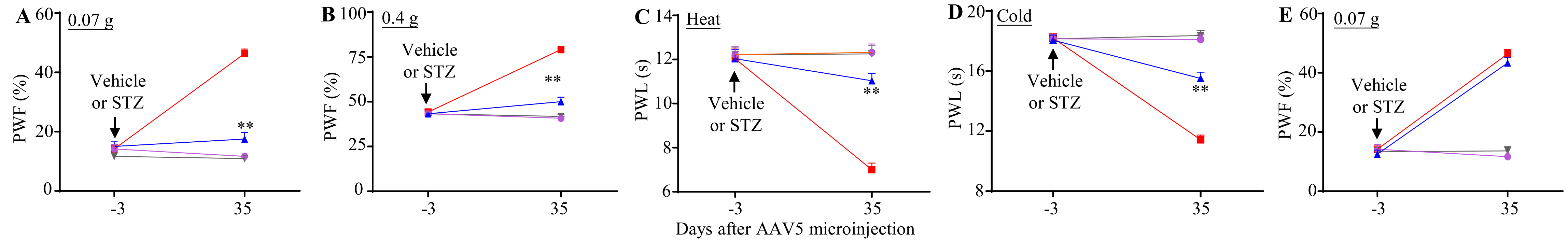

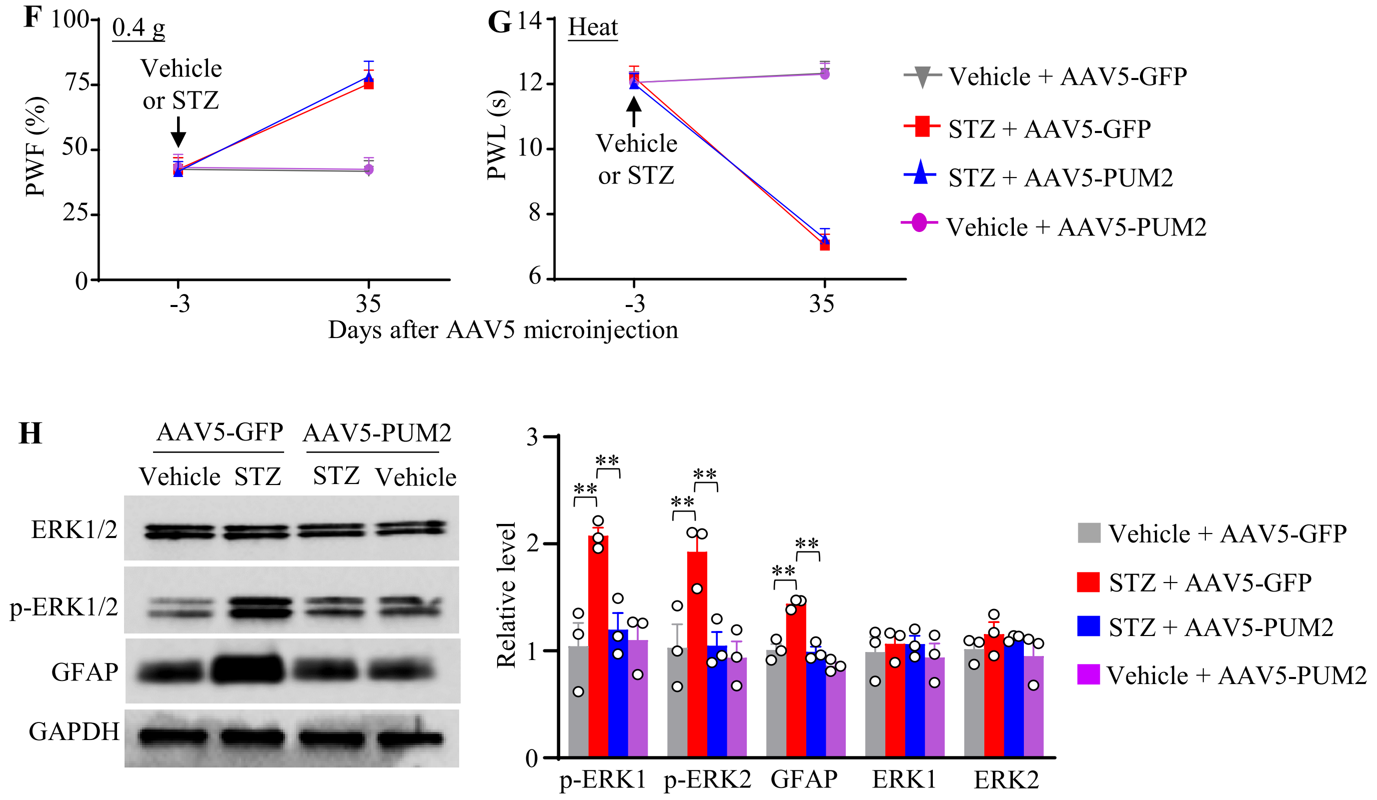

**Supplementary Fig. 20.** DRG overexpression of PUM2 attenuates the STZ-induced nociceptive hypersensitivity. (A-G) Effect of unilateral microinjection of AAV5-PUM2 or AAV5-GFP into the L3/4 DRGs on paw withdrawal frequency (PWF) to 0.07-g (A and E) and 0.4-g (B and F) von Frey filament stimuli and paw withdrawal latency (PWL) to heat (C and G) and cold (D) stimuli on the indicated days after viral microinjection in male mice that received STZ or vehicle injection 3 days before viral microinjection, assessed on the ipsilateral (A–D) and contralateral (E–G) sides. n = 11 mice/group. $^{**}P < 0.01$, three-way repeated-measures ANOVA followed by Tukey's post hoc test. (H) Effect of unilateral microinjection of AAV5-PUM2 or AAV5-GFP into the L3/4 DRGs on levels of p-ERK1/2, total ERK1/2 and GFAP in the ipsilateral L3/4 spinal cord dorsal horn 35 days after viral microinjection in male mice that received STZ or vehicle injection 3 days before viral microinjection. n = 3 mice/group. $^{**}P < 0.01$, two-way ANOVA followed by Tukey's post hoc test.

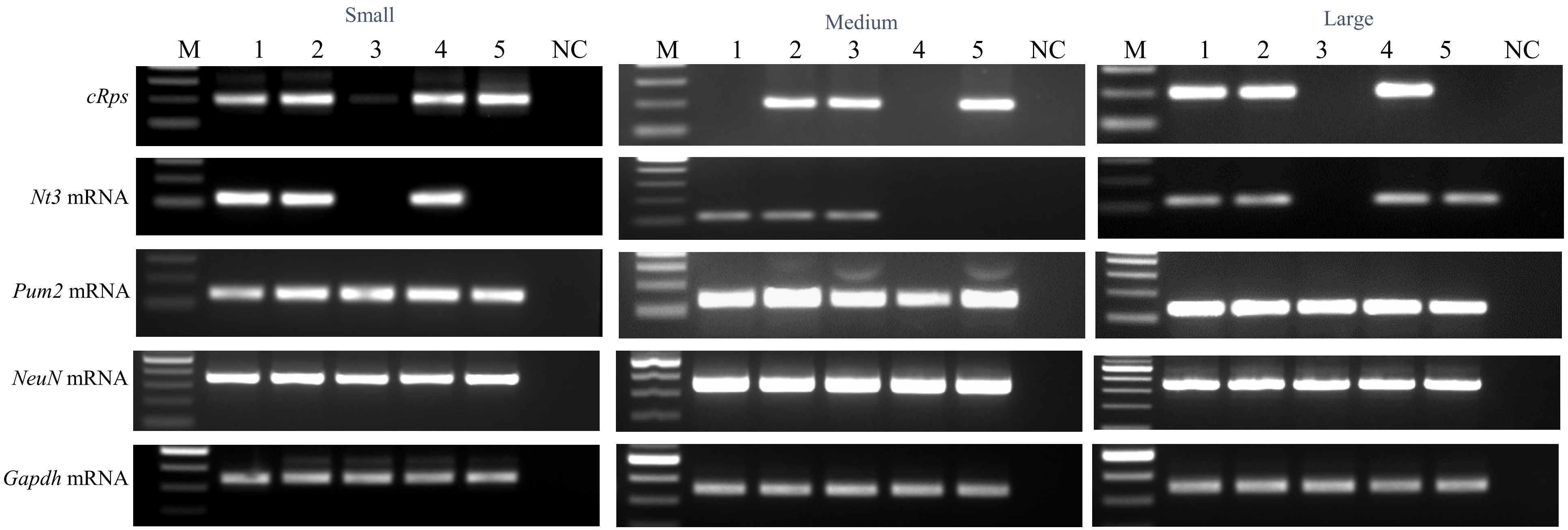

**Supplementary Fig. 21.** Single cell RT-PCR assay shows co-expression of *cRps*, *Nt3* mRNA, and *Pum2* mRNA in individual dorsal root ganglion (DRG) neurons. *NeuN* mRNA is used as a neuronal marker. *Gapdh* mRNA was used as a loading control. Large DRG neuron: > 35 µm in diameter. Medium DRG neuron: 25-35 µm in diameter. Small DRG neuron: < 25 µm in diameter. n = 5 neurons/size. Number 1–5 indicates five different neurons. M: DNA ladder marker. NC: a negative control (without cDNA).

**Supplementary Table 1**: Locomotor function

| Treatment groups | Placing | Grasping | Righting |
|---|---|---|---|
| Vehicle | 5 (0) | 5 (0) | 5 (0) |
| STZ | 5 (0) | 5 (0) | 5 (0) |
| Normal Diet +Vehicle | 5 (0) | 5 (0) | 5 (0) |
| High Fat Diet + STZ | 5 (0) | 5 (0) | 5 (0) |
| AAV5-*Scr shRNA* + Vehicle | 5 (0) | 5 (0) | 5 (0) |
| AAV5-*cRps6kb1 shRNA* + Vehicle | 5 (0) | 5 (0) | 5 (0) |
| AAV5-*Scr shRNA* + STZ | 5 (0) | 5 (0) | 5 (0) |
| AAV5-*cRps6kb1 shRNA* + STZ | 5 (0) | 5 (0) | 5 (0) |
| *cRps6kb1* Blocker + Vehicle | 5 (0) | 5 (0) | 5 (0) |
| *cRps6kb1* Blocker + STZ | 5 (0) | 5 (0) | 5 (0) |
| Scr siRNA + Vehicle | 5 (0) | 5 (0) | 5 (0) |
| *Nt3* siRNA + Vehicle | 5 (0) | 5 (0) | 5 (0) |
| Scr siRNA + STZ | 5 (0) | 5 (0) | 5 (0) |
| *Nt3* siRNA + STZ | 5 (0) | 5 (0) | 5 (0) |
| AAV5-*Gfp* | 5 (0) | 5 (0) | 5 (0) |
| AAV5-*cRps6kb1* | 5 (0) | 5 (0) | 5 (0) |
| AAV5-*Nt3* | 5 (0) | 5 (0) | 5 (0) |
| AAV5-*Gfp* + Scr siRNA | 5 (0) | 5 (0) | 5 (0) |
| AAV5-*Gfp* + *Nt3* siRNA | 5 (0) | 5 (0) | 5 (0) |
| AAV5-*cRps6kb1* + Scr siRNA | 5 (0) | 5 (0) | 5 (0) |
| AAV5-*cRps6kb1* + *Nt3* siRNA | 5 (0) | 5 (0) | 5 (0) |
| $Nt3^{fl/fl}$ + Vehicle + AAV5-*Gfp* | 5 (0) | 5 (0) | 5 (0) |
| $Nt3^{fl/fl}$ + Vehicle + AAV5-*Cre* | 5 (0) | 5 (0) | 5 (0) |
| $Nt3^{fl/fl}$ + STZ + AAV5-*Gfp* | 5 (0) | 5 (0) | 5 (0) |
| $Nt3^{fl/fl}$ + STZ + AAV5-*Cre* | 5 (0) | 5 (0) | 5 (0) |
| AAV5-*Gfp* + Vehicle | 5 (0) | 5 (0) | 5 (0) |
| AAV5-*Gfp* + STZ | 5 (0) | 5 (0) | 5 (0) |
| AAV5-*Pum2* + Vehicle | 5 (0) | 5 (0) | 5 (0) |
| AAV5-*Pum2* + STZ | 5 (0) | 5 (0) | 5 (0) |
| AAV5-*Gfp* + AAV5-*cRps6kb1* | 5 (0) | 5 (0) | 5 (0) |
| AAV5-*Gfp* + AAV5-*Pum2* | 5 (0) | 5 (0) | 5 (0) |
| AAV5-*cRps6kb1* + AAV5-*Pum2* | 5 (0) | 5 (0) | 5 (0) |
| ASO + STZ | 5 (0) | 5 (0) | 5 (0) |
| ASO + Vehicle | 5 (0) | 5 (0) | 5 (0) |
| MSO + STZ | 5 (0) | 5 (0) | 5 (0) |
| ASO + HFD | 5 (0) | 5 (0) | 5 (0) |
| ASO + ND | 5 (0) | 5 (0) | 5 (0) |
| AAV5-*cRps6kb1 shRNA* + HFD | 5 (0) | 5 (0) | 5 (0) |
| AAV5-*cRps6kb1 shRNA* + ND | 5 (0) | 5 (0) | 5 (0) |

All values are Mean (SEM). n = 8-12 mice/group. 5 trials. STZ – Streptozocin. Scr – Scrambled

ASO: Antisense Oligonucleotide. MSO: Missense Oligonucleotide.

**Supplementary Table 2:** Potential *cRps6kb1* binding proteins identified by using ChIRP-MS assay

| Gene Symbol | Accession | Coverage [%] | # PSMs | MW [kDa] | calc. pI | Ratio |
|---|---|---|---|---|---|---|
| EIF4AIII | E9PV04 | 27.01 | 16 | 46.8 | 7.43 | 1.9 |
| SNRPA | Q62189 | 36.93 | 23 | 31.8 | 9.8 | 1.7 |
| HEXIM1 | Q8R409 | 21.35 | 4 | 40.2 | 5.38 | 1.7 |
| RPL13A | P19253 | 34.48 | 17 | 23.4 | 11.02 | 1.6 |
| RPL23 | P62830 | 37.86 | 9 | 14.9 | 10.51 | 1.6 |
| SRSF5 | O35326 | 30.11 | 11 | 30.9 | 11.56 | 1.5 |
| RBM8A | Q9CWZ3 | 31.61 | 6 | 19.9 | 5.72 | 1.5 |
| LSM4 | D3YTP8 | 38.54 | 7 | 11.1 | 8.88 | 1.8 |
| PUM2 | Q3TQ29 | 11.84 | 21 | 105.6 | 6.86 | 1.5 |
| O70435 | | 32.16 | 9 | 28.4 | 5.44 | 2.25 |
| RBM8A | Q9CWZ3 | 31.61 | 6 | 19.9 | 5.72 | 1.5 |
| RNPS1 | Q99M28 | 12.79 | 6 | 34.2 | 11.84 | 1.5 |
| Ago2 | Q8CJG0 | 56.40 | 85 | 97.2 | 9.19 | 1.1 |

*cRps6kb1:* biotinylated full-length sense *cRps6kb1* RNA, *mRps6kb1:* biotinylated full-length sense *mRps6kb1* RNA

Table 3. All primers, probes and siRNAs used

| Names | Sequences |
|---|---|
| *RT-PCR* | |
| m/h-cRps6kb1 F | 5′-TAAAGGGGGCTATGGAAAGG -3′ |
| m/h-cRps6kb1 R | 5′- TTTTCTGGCCCTCTGTTCAC -3′ |
| m-lin-Rps6kb1 F | 5′- TCAGCCCAGTCAAATTCTCTC -3′ |
| m-lin-Rps6kb1 R | 5′- GTCACATCCATCTGCTCTATCC -3′ |
| m-NT3 F | 5′- CTGCCACGATCTTACAGGTG -3′ |
| m-NT3 R | 5′- CTTGGATGCCACGGAGATAAG -3′ |
| m-PUM2 F | 5′- GTATTCTCAGCAGGCCTTGC -3′ |
| m-PUM2 R | 5′- AACCACTGCTGGACCTGCTAA -3′ |
| m-GAPDH F | 5′- TTGTGGAAGGGCTCATGACC- 3′ |
| m-GAPDH R | 5′-GTTGAAGTCGCAGGAGACAA -3′ |
| m-Tubulin F | 5′- GTGCATCTCCATCCATGTTG-3′ |
| m-Tubulin R | 5′- GTGGGTTCCAGGTCTACGAA-3′ |
| m-NT3-3’UTR F | 5′- CAGGCCTTTCCCATCTGTTA -3′ |
| m-NT3-3’UTR R | 5′- CATGGCCTTGACAATACTGAATG -3′ |
| m-H19 F | 5′- CATTCTAGGCTGGGGTCAAA-3′ |
| m-H19 R | 5′- GCCCTTCTTTTCCATTCTCC-3′ |
| m-Malat1 F | 5′- TCATACCTAACCAGGCATAACA-3′ |
| m-Malat1 R | 5′- AAGTGCTCACAAGGCAAATC-3′ |
| m-cPlekha5 F | 5′- GCTCATGCAGCTGAAACTTG-3′ |
| m-cPlekha5 R | 5′- CCGTCCTTCTGAAATCCGTA-3′ |
| m-cRps6kb1 F | 5′- GCCCGGAGAATCTGAAGAC-3′ |
| m-cRps6kb1 R | 5′- GATGTGTTCCTCCCGAATGT-3′ |
| h-GAPDH F | 5′- TCACCATCTTCCAGGAGCG -3′ |
| h-GAPDH R | 5′- CTGCTTCACCACCTTCTTGA -3′ |
| h-NT3 F | 5′- TACAGGTGAACAAGGTGATGTC -3′ |
| h-NT3 R | 5′- GAGGGAATTGAGCGAGTCTTC -3′ |
| h-Rps6kb1 F | 5′- GGCATGGAACATTGTGAGAAAT -3′ |
| h-Rps6kb1 R | 5′- CCTTTACCAAGTACCCGAAGTAG -3′ |
| *siRNA* | |
| si-mNT3 –S1 | 5′- CAACAGAGACGCUACAAUUtt -3′ |
| si-mNT3 –AS1 | 5′- AAUUGUAGCGUCUCUGUUGtt -3′ |
| si-mNT3 –S2 | 5′- GUAACUCUCCUGUGAAACAtt -3′ |
| si-mNT3 –AS2 | 5′- UGUUUCACAGGAGAGUUACtt -3′ |
| si-mNT3 –S3 | 5′- CCAUCUGUUAACCUUGUUUtt -3′ |
| si-mNT3 –AS3 | 5′- AAACAAGGUUAACAGAUGGtt -3′ |
| *Vector construction* | |
| AAV-cRps6kb1 F1 | 5′- TGACCTCCATAGAAGACACC -3′ |
| AAV-cRps6kb1 R1 | 5′- ATGCTTTCATTTAACTGACCCTGAAGTATAAAAAAAAAGTCA -3′ |
| AAV-cRps6kb1 F2 | 5′- GGTCAGTTAAATGAAAGCAT -3′ |
| AAV-cRps6kb1 R2 | 5′- CTTTTTAAGCACCTTCATGG -3′ |
| AAV-cRps6kb1 F3 | 5′- CCATGAAGGTGCTTAAAAAGGTAAGAAGCAAGGTTTCATT -3′ |
| AAV-cRps6kb1 R3 | 5′- AACTCCATCACTAGGGGTTC -3′ |
| AAV-NT3 F | 5′- TCCCCGCGGGCCACCATGAATCATGATTTTCAAGCTC-3′ |
| AAV-NT3 R | 5′- GGCCTCGAGATGTCCATCTTGTTTTATG -3′ |
| AAV-PUM2 F | 5′- TCCCCGCGGGCCACCATGAATCATGATTTTCAAGCTC-3′ |
| AAV-PUM2 R | 5′- CCGCTCGAGTTACAGCATCCCATTTGGTGGTC-3′ |
| AAV-sh-cRps6kb1 F | 5′- GATCCGCTTAAAAAGGGTCAGTTAGAAGCTTGTAACTGACCCTTTTTAAGCTTTTTTT -3′ |
| AAV-sh-cRps6kb1 R | 5′- CTAGAAAAAAAGCTTAAAAAGGGTCAGTTACAAGCTTCTAACTGACCCTTTTTAAGCG -3′ |
| *psiCheck*-NT3-3’UTR F | 5′- CCGCTCGAGATTGGCATCTGTCCCCACATA -3′ |

| | |
|---|---|
| *psiCheck*-NT3-3’UTR R | 5′- GCACGCGGCCGCTCCGATTTTAAGAAAGTTTACTG -3′ |
| *Single cell RT-PCR* | |
| m-cRps6kb1 F | 5′- TAAAGGGGGCTATGGAAAGG -3′ |
| m-cRps6kb1 R | 5′- TTTTCTGGCCCTCTGTTCAC -3′ |
| mNT3 F | 5′- CTGCCACGATCTTACAGGTG -3′ |
| mNT3 R | 5′- CTTGGATGCCACGGAGATAAG -3′ |
| mPUM2 F | 5′- GTATTCTCAGCAGGCCTTGC -3′ |
| mPUM2 R | 5′- AACCACTGCTGGACCTGCTAA -3′ |
| mRbfox3(NeuN) F | 5′- ACATTTCGAGCTGCACCA-3′ |
| mRbfox3(NeuN) R | 5′- CCGATGGTGTGATGGTAAGG-3′ |
| mGAPDH F | 5′- TTGTGGAAGGGCTCATGACC- 3′ |
| mGAPDH R | 5′- GTTGAAGTCGCAGGAGACAA -3′ |
| *RIP-PCR* | |
| RIP-RPS-I1-R1 F | 5′- TGAAGCACACACAGGATTACT-3′ |
| RIP-RPS-I1-R1 R | 5′- TCTGGCTCTACTCCAAAACCACT-3′ |
| RIP-RPS-I1-R2 F | 5′- TTGTCATTGATCACACAAATA -3′ |
| RIP-RPS-I1-R2 R | 5′- ATTAACTACATAAAGCCTATACT -3′ |
| RIP-RPS-I1-R3 F | 5′- GCTGTCTTTAGACACACCAGAG -3′ |
| RIP-RPS-I1-R3 R | 5′- CTCTTCCAAAGGTCCTGAGTTCA3′ |
| RIP-RPS-I4-R1 F | 5′- AGATAGATATGGCTGCGTG -3′ |
| RIP-RPS-I4-R1 R | 5′- GACTTAAATATTAGTGACTCTG -3′ |
| RIP-RPS-I4-R2 F | 5′- TCGAACTCAAAGGGGATT -3′ |
| RIP-RPS-I4-R2 R | 5′- ATTTCTCCTTTAAGTGTGTG -3′ |
| m-NT3-3’UTR F | 5′- CAGGCCTTTCCCATCTGTTA -3′ |
| m-NT3-3’UTR R | 5′- CATGGCCTTGACAATACTGAATG -3′ |
| m-cRps6kb1 F | 5′-TAAAGGGGGCTATGGAAAGG -3′ |
| m-cRps6kb1 R | 5′- TTTTCTGGCCCTCTGTTCAC -3′ |
| *Insitu Hybridization* | |
| NT3-PRF | 5′- ACCACGGAGGAAACGCTATGTGTGGGATCAACATTCCCAAC-3’ |
| NT3-PRR | 5′- CCCCGAATGTCAATGGCTGA -3′ |
| *Northern blot* | |
| F | 5′- TAATACGACTCACTATAGGATATGAACTTGGCATGGAACATTGTG-3’ |
| R | 5′- GGTCCAACTCCCCCATGGTCCATG -3′ |
| *Mass Spec Probe* | |
| F | 5′- TAATACGACTCACTATAGGGAGAGGTCAGTTAAATGAAAGC-3’ |
| R | 5′- CTTTTTAAGCACCTTCATGGC -3′ |

RT: Reverse-transcription; F, Forward; R, Reverse. m-, mice. h-, human. *cRps6kb1*, *circRps6kb1*. *mRps6kb1*, *Rps6kb1* mRNA. sh, short hairpin RNA. Mass Spec, Mass Spectrometry
